# An initial genome editing toolset for *Caldimonas* t*hermodepolymerans*, the first model of thermophilic polyhydroxyalkanoates producer

**DOI:** 10.1101/2024.09.22.614348

**Authors:** Anastasiia Grybchuk-Ieremenko, Kristýna Lipovská, Xenie Kouřilová, Stanislav Obruča, Pavel Dvořák

## Abstract

The limited number of well-characterized model bacteria cannot address all the challenges in a circular bioeconomy. Therefore, there is a growing demand for new production strains with enhanced resistance to extreme conditions, versatile metabolic capabilities, and the ability to utilize cost-effective renewable resources while efficiently generating attractive biobased products. Particular thermophilic microorganisms fulfill these requirements. Non-virulent Gram-negative *Caldimonas thermodepolymerans* DSM15344 is one such attractive thermophile that efficiently converts a spectrum of plant biomass sugars into high quantities of polyhydroxyalkanoates (PHA) - a fully biodegradable substitutes for synthetic plastics. However, to enhance its biotechnological potential, the bacterium needs to be “domesticated”. In this study we established effective homologous recombination and transposon-based genome editing systems for *C. thermodepolymerans*. By optimizing the electroporation protocol and refining counterselection methods, we achieved significant improvements in genetic manipulation and constructed the AI01 chassis strain with improved transformation efficiency and a Δ*phaC* mutant that will be used to study the importance of PHA synthesis in *Caldimonas*. The advances described herein highlight the need for tailored approaches when working with thermophilic bacteria and provide a springboard for further genetic and metabolic engineering of *C. thermodepolymerans*, which can be considered the first model of thermophilic PHA producer.

## 1. Introduction

Nearly 400 million tons of petroleum-based plastics are produced annually, and global demand for plastics is projected to double by 2050 (Stegmann *et al*., 2022). Even if we soon learn how to recycle most of these synthetic polymers, this will not solve the phenomenal problem of microplastic pollution caused by the slow erosion of non-biodegradable plastics and our dependence on fossil resources for *de novo* synthesis of these chemicals (Lim, 2021). Petroplastics such as polyethylene or polypropylene can be replaced in many applications by polyhydroxyalkanoates (PHA) - biocompatible, non-toxic, recyclable and fully biodegradable polyesters that are synthesized by many bacteria and archaea as a storage material making up to 90% of cell dry mass (CDM) (Choi *et al*., 2020; Dvořák *et al*., 2020; Meereboer *et al*., 2020). When produced from renewable feedstocks such as lignocellulosic residues, PHA can also reduce the carbon footprint of plastics production (Yu and Chen, 2008; Stegmann *et al*., 2022). However, in 2023 PHA will still account for less than 0.02% of annual industrial production of plastics, mainly because their biomanufacturing is several times more expensive than the chemical synthesis of petroleum-based plastics (Choi *et al*., 2020; Palmeiro-Sánchez *et al*., 2022). A substantial portion of the cost of PHA is due to expensive substrates used to feed production strains (mainly starch-derived glucose) and operational costs including material and media sterilization to prevent process contamination. The concept of Next Generation Industrial Biotechnology (NGIB), based on the use of extremophilic microorganisms, has brought some promise to the field in the last decade with increased bioprocess robustness (Chen and Jiang, 2018; Ma *et al*., 2020; Zheng *et al*., 2020). The use of engineered *Halomonas* strains capable of producing tailor-made PHA in open, non-sterile conditions in saline water has enabled the emergence of several new ventures that are attacking the global market with PHA at prices approaching those of synthetic plastics (Ye *et al*., 2018; Zheng *et al*., 2020). The introduction of robust extremophiles can indeed be a real game-changer for the field, but it is still not clear whether halophiles are the only best choice for PHA biotechnology.

We have recently carried out a thorough search on the thermophilic PHA producers that are attractive candidates for NGIB due to high PHA yields and range of utilizable substrates (Obruča *et al*., 2022). Thermophile cultures do not generate saltwater waste, which is a significant environmental concern associated with halophilic cultures (Oren, 2011). The safe Gram-negative (G-) bacterium *Caldimonas thermodepolymerans* DSM15344 (formerly *Schlegelella thermodepolymerans*) emerged as a very promising thermophilic PHA producer in the following experimental trials by several laboratories (Kourilova *et al*., 2020, 2021; Zhou *et al*., 2023; Bertran-Llorens *et al*., 2024) *C. thermodepolymerans* was found to accumulate unprecedented amounts of poly(3-hydroxybutyrate) (PHB, up to 87% of cell dry weight) from D-xylose at its optimum growth temperature of 50 °C in inexpensive minimal medium (Kourilova *et al*., 2020; Zhou *et al*., 2023). DSM15344 grows well and produces PHB on other attractive sugars available from renewable lignocellulose - L-arabinose, D-mannose, D-galactose or D-cellobiose (dimer of D-glucose) (Kourilova *et al*., 2020). *C. thermodepolymerans* DSM15344 is also well resistant to the major inhibitors in lignocellulosic hydrolysates such as gallic acid, furfural, levulinic acid, or ferulic acid (Kourilova *et al*., 2021). These properties are unique among characterized extremophiles and desirable in PHA biotechnology (Dietrich *et al*., 2019). Moreover, *Caldimonas*’ optimum growth temperature makes it more compatible with (hemi)cellulose saccharification protocols, allows better dissolution of sugar substrates and prevents contamination with mesophilic microflora (Bosma *et al*., 2013; Obruča *et al*., 2022; Ye *et al*., 2022). Moderate thermophiles can also save energy in biotechnologies, as in industrial fermentations more energy is used for cooling than heating (Turner *et al*., 2007). The whole genome sequence of DSM15344 and several other thermophilic *Caldimonas* strains is now available, as well as a basic metabolic map of the central carbon metabolism of DSM15344 (Musilova *et al*., 2023). Thorough searches of the genomic data provided some insights into the carbohydrate and PHA metabolism, restriction-modification systems or CRISPR arrays in DSM15344 and several related *Caldimonas* strains (Musilova *et al*., 2023). Recently, it was reported that *C. thermodepolymerans* is also capable of efficient and quick conversion of ferulic acid into valuable products such as vanillyl alcohol and vanillic acid (Hrabalova et al. 2024). *C. thermodepolymerans* can be a true turning-point host, but its potential is not fully exploited due to limited growth on glucose, a product portfolio restricted to simple PHB homopolymer (which, unlike other PHA polymers, is limited in applications by mechanical and technological properties), or suboptimal PHA productivity. The wild-type bacterium needs an engineering push to reach the level of PHA super-producer and game changer. However, compared to model mesophilic bacteria, the genetic and metabolic engineering toolkits for Gram-negative thermophiles are practically non-existent. Rare examples of attempts to engineer G-thermophiles include studies on *Thermus thermophilus* (Carr *et al*., 2015) or *Caldimonas manganoxidans* (Arai *et al*., 2022). But there are no available engineering plasmid-based tools validated for G-thermophilic PHA producers. In this study, we aimed to develop and test an initial genome editing toolbox for *C. thermodepolymerans*. Present work includes optimization of the transformation and genome editing protocols as well as testing of prepared mutants for their ability to produce PHA. The study pioneers the biotechnological domestication of *C. thermodepolymerans* and potentially other G-thermophiles and paves the way for further boosting of their bioproduction potential by metabolic engineering.

## 2. Materials and Methods

### 2.1 Bacterial strains and cultivation conditions

*C. thermodepolymerans* DSM 15344 and *Escherichia coli* strains were routinely cultured either in 50 mL of lysogeny broth (LB, Serva) in shake flasks (250 mL) or on agar (1.6 % w/v) plates containing LB. For *C. thermodepolymerans*, growth conditions included incubation at 50 °C, 42 °C, or 37 °C, while *E. coli* was cultured at 37°C. Antibiotics were added into the medium, when necessary, with kanamycin (Km) at a concentration of 12.5 µg/mL and gentamicin (Gm) at 10 µg/mL for *Caldimonas*. For *E. coli*, Km at 50 µg/mL and ampicillin (Amp) at 150 µg/mL were used.

Growth of *Caldimonas* at 37 °C and 50 °C in LB medium was tested using a Cell Growth Quantifier (CGQ) optical sensor-based technology for non-invasive online biomass monitoring in shake flasks by Aquila Biolabs / Scientific Bioprocessing. Cultivation at each temperature lasted 24 h and was conducted in biological triplicates.

### 2.2 Transformation of bacterial cells with plasmid DNA

The delivery of plasmid DNA into *Caldimonas* cells was done by electroporation and triparental mating, while heat-shock transformation was employed for *E. coli*. Electrocompetent *Caldimonas* cells were prepared using a previously established protocol for *C. manganoxidans* (Arai *et al*., 2022), with following modifications that helped to improve transformation efficiency for *C. thermodepolymerans*. Bacterial cells from the night culture were inoculated into 100 mL of LB medium to the starting OD_600_ of 0.05 and shaken at 220 rpm (ES-20/60, Shaker-incubator, Biosan) at 37°C. Upon reaching an OD_600_ of ∼0.5, the bacterial culture was rapidly cooled on ice, and three rounds of washing with 10 % glycerol in milli-Q water followed. The centrifugation in each round was done at 4,000 x g for 10 min for rounds 1 and 2 and 20 min for round 3. The cells were then resuspended in 1.5 mL of ice-cold 10 % glycerol, were aliquoted and frozen at −60 °C. In the electroporation protocol, Eppendorf tubes containing 100 µl of cells were melted on ice and mixed with 100 ng of plasmid DNA. Electroporation was performed using a Biorad Gene Pulser Xcell (2,500 V, 25 μF, 200 Ω) with a 2 mm gap cuvette. Immediately following the pulse, 1 mL of 37 °C prewarmed LB was added to the cells, which were then allowed to recover at 37 °C for 2 h. Subsequently, 100 µl of the cell suspension was plated on agar plates containing selective antibiotic, and cells were cultivated at 37°C, 42°C, or 50°C. Electroporation efficiency was quantified as the number of colony forming units (CFU) per 1 µg of plasmid DNA.

In the triparental mating protocol, three distinct strains were employed: the donor strain (*E. coli* CC118 λpir) carrying the mobilizable plasmid, the helper strain (*E. coli* HB101 with plasmid pRK600), and the recipient strain (*C. thermodepolymerans*). These strains were cultured for 24 h with appropriate antibiotics at 37 °C. Then, 2 mL of cells were collected and diluted in 1 mL of 10 mM MgSO_4_. The ODs at 600 nm of all three strains were measured. Subsequently, the three bacterial cultures were mixed in 1 mL of MgSO4, ensuring a 1:1:1 ratio of strains in the mixture. The resulting cell suspension (10 µl) was plated as separate droplets onto an LB agar plate without antibiotics. The mixed culture was then incubated for 48 h at 37°C. The cells were delicately scraped across the droplet with a sterile tip and resuspended in 1 mL of 10 mM MgSO_4_. The resulting cell suspension was serially diluted in MgSO_4_, with 100 µl of the diluted suspension plated onto LB agar plates supplemented with Km and incubated at 50 °C overnight. Single colonies obtained were subject to two additional restreakings to obtain pure clones.

Plasmid isolation was conducted with an E.Z.N.A. Plasmid DNA Mini Kit (Omega Bio-tek). The standard isolation protocol was used for *E. coli*. For *Caldimonas*, the centrifugation time with the neutralization buffer was extended from the standard 10 minutes to 20 minutes to achieve better purity and higher concentration.

### 2.3 Testing of the influence of methylation pattern on transformation efficiency

*C. thermodepolymerans* cells were electroporated with pSEVA238. Plasmid DNA derived from distinct isolation sources: (i) standard *E. coli* strain CC118, (ii) *E. coli* strain JM110 (*dam*^-^ *dcm*^-^) characterized by the absence of adenosine and cytosine methyltransferases, and (iii) the type *Caldimonas* strain DSM15344. Transformants were incubated on LB agar plates supplemented with Km for 36 h. Transformation efficiencies were calculated per 1 µg of transformed DNA

### 2.4 Targeted gene deletions in C. thermodepolymerans chromosome using homologous recombination-based method

We intended to adopt for genomic editing of *C. thermodepolymerans* two previously reported homologous recombination-based protocols. Both protocols use a single plasmid for targeted edits. The first one takes advantage of green fluorescent protein gene *gfp* as a selectable marker (Volke et al., 2020), while the second one uses levan sucrase gene *sacB* (Ried and Collmer, 1987). Both protocols were modified to enable seamless deletions in the chromosome of *C. thermodepolymerans*.

All primers used in this study are listed in Table S1. The primer annealing temperature was estimated with NEB Tm Calculator. The deletion vector, pSNW2_HA2 (Table 1), was assembled through restriction cloning, utilizing the pSNW2 (Volke *et al*., 2020) as a backbone. Overlap extension PCR (Higuchi *et al*., 1988) was used to fuse homologous regions (including start and stop codons) immediately adjacent to the *phaC* gene open reading frame. The primers HAN1, HR1rev, HR2for, HAN2, HA1for, HA2_XbaIrev were employed. The PCR was performed using Q5 polymerase (New England Biolabs) following the manufactureŕs standard protocol. The PCR product was purified with NucleoSpin Gel and PCR Clean-up kit (Macherey-Nagel) and cloned into the pSNW2 plasmid (Table 1) between *Eco*RI and *Xba*I restriction sites. The CC118 λπ chemically competent *E. coli* cells were used for the plasmid propagation. Insertion of homologous arms into the pSNW2 plasmid and the integration of the entire vector construct into the *C. thermodepolymerans* chromosome were verified by PCR using primers pSNW2 flank for, pSNW2 flank rev, Gen1for, and Gen2 rev. To conduct colony PCR on *Caldimonas* transformants, we utilized PCRBIO Taq Mix Red and VeriFi® Mix Red (PCR Biosystems, United Kingdom). The PCR reactions were carried out in a 10 µl volume, following the manufacturer’s protocol, and additionally supplemented with 5x Q5 High GC Enhancer (NEB). The correct assembly of the vector and the deletion of the *phaC* gene were confirmed through Sanger sequencing. The vector pSNW2_HA2 was introduced into *C. thermodepolymerans* through electroporation and triparental mating. Following the isolation of individual colonies, the isolates were transferred to 3 mL of LB medium in 15 mL Falcon tubes and incubated at 50°C with shaking at 220 rpm (ES-20/60, Shaker-incubator, Biosan). The cells were passaged three times, with 500 µl inoculated into the next passage every 24 h. Subsequently, liquid cultures were plated onto LB agar to obtain single colonies. The deletion in isolates that exhibited a loss of fluorescence was confirmed by PCR using the primers Gen1 for, Gen2 rev and then by the Sanger sequencing.

**Table 1.** Strains and plasmids used in this work.

| Strain/Plasmid | Characteristics | Source |
| --- | --- | --- |
| **Strains** |  |  |
| *Escherichia coli* |  |  |
| **CC118λpir** | Cloning strain: *araD139* $\Delta$(*ara-leu*)7697 Δ*lacX74 galE galK phoA20 thi-1 rpsE rpoB(Rif<sup>R</sup>) argE(Am) recA1* | (Manoil and Beckwith, 1985) |
| **Dh5a** | Cloning strain: *dlacZ* $\Delta$ *M15* $\Delta$ (*lacZYA-argF*) U169 *recA1 endA1 hsdR17* (rK-mK+) *supE44 thi-1 gyrA96 relA1* | (Taylor *et al.*, 1993) |
| **JM110** | Cloning strain: *rpsL, thr, leu, thi, lacY, galK, galT, ara, tomA, tsx, dam-, dcm-, supE44, \Delta(lac-proAB), F' traD36, proAB, lacIqZ\Delta M15* | (Yanisch-Perron *et al.*, 1985) |
| **HB101** | Helper strain for triparental mating: *F- λ-hsdS20(rB-mB-) recA13 leuB6(Am) araC14 \Delta(gpt-proA)62 lacY1 galK2(OC) xyl-5 mtl-1 thiE1 rpsL20(Sm<sup>R</sup>) glnX44(AS)* | (Boyer and Roulland-dussoix, 1969) |
| **One Shot™ PIR1** | Cloning strain: *F- \Delta lac169 rpoS(Am) robA1 creC510 hsdR514 endA recA1 uidA(\Delta MLul)::pir-116* | Invitrogen |
| ***C. thermodepolymerans* DSM 15344** | Type strain | (Elbanna *et al.* 2003) |
| ***C. thermodepolymerans* Δ*phaC*** | Derivative of DSM 15344 with deleted gene of PHA synthase (IS481_08630) | This study |
| ***C. thermodepolymerans* AI01** | Derivative of DSM 15344: ΔIS481_08585, ΔIS481_14855, ΔIS481_14025 (deleted genes of restriction endonucleases from innate R-M systems) | This study |
| **Plasmids** |  |  |
| **pSEVA238** | Medium-copy-number expression plasmid; *oriV* (pBBR1), *XylS/Pm* expression system; Km<sup>R</sup> | (Silva-Rocha *et al.*, 2013) |
| **pSEVA238_ *gfp*** | Derivative of pSEVA238 with *gfp* gene | Laboratory stocks |
| **pSEVA631** | Medium-copy-number expression plasmid; *oriV* (pBBR1), Gm<sup>R</sup> | (Silva-Rocha *et al.*, 2013) |
| **pSNW2** | Derivative of vector pGNW2 with P14g(BCD2)→msfGFP; Km<sup>R</sup> | (Volke *et al.* 2020) |
| **pCJ012** | Plasmid for deletion of *lldD* gene of lactate dehydrogenase in *P. putida* via homologous recombination and SacB-mediated sucrose counter-selection | (Johnson and Beckham, 2015) |
| **pRK600** | Helper plasmid for tri-parental mating: *oriV*(ColE1) RK2 *tra<sup>+</sup> mob<sup>+</sup>*, Cm<sup>R</sup> | (Kessler *et al.*, 1994) |
| **pBAMD1-6_ *glf*** | Mini-Tn5 delivery plasmid with subcloned *glf* gene of glucose facilitator from *Zymomonas mobilis*: *ori*(R6K) *bla aacC1*, Amp<sup>R</sup> Gm<sup>R</sup> | (Martnez-Garca *et al.*, 2014; Bujdos *et al.*, 2023) |
| **pBAMD_gfp** | pBAMD1-6 with *gfp* gene of monomeric superfolder green fluorescent protein cloned into *EcoRI* and *HindIII* | This study |
| **pAI012_HA2** | Derivative plasmid of pCJ012 for deletion of *phaC* gene via homologous recombination and mesophilic SacB-mediated sucrose counter-selection | This study |
| **pSNW2_HA2** | Derivative plasmid of pSNW2 for deletion of *phaC* gene via homologous recombination and GFP-mediated selection | This study |
| **pSNW2_8585** | Derivative plasmid of pSNW2 for deletion of IS481_08585 restriction endonuclease | This study |
| **pSNW2_14855** | Derivative plasmid of pSNW2 for deletion of IS481_14855 restriction endonuclease | This study |
| **pSNW2_14025** | Derivative plasmid of pSNW2 for deletion of IS481_14025 restriction endonuclease | This study |
| **pAI_ThSacB** | Derivative plasmid of pSNW2 for deletion of *phaC* gene via homologous recombination and thermostable SacB-mediated sucrose counter-selection | This study |

Vectors pSNW2_8585, pSNW2_14855, and pSNW2_14025 for the seamless deletions of IS481_08585, IS481_14855 and IS481_14025 restriction endonuclease (RE) genes were constructed in a similar way as described above. The pSNW2 vector was used as a backbone. Homologous arms of the IS481_08585 RE were amplified with the primers 08585_HomS1 for, 08585_HomS1 rev, 08585_HomS2 for, 08585_HomS2 rev, EcoRI _08585_HomS for and XbaI _08585_HomS rev. Amplification of homologous arms for IS481_14855 RE was performed with primers 14855_HomS1 for, 14855_HomS1 rev, 14855_HomS2 for, 14855_HomS2 rev, KpnI _14855_HomS for and XbaI _14855_HomS rev. Arms of homology for IS481_14025 RE were amplified using primers 14025_HomS1 for, 14025_HomS1 rev, 14025_HomS2 for, 14025_HomS2 rev, EcoRI _14025_HomS for and XbaI _14025_HomS rev. The propagation of the constructs was carried out in DH5α λπ chemically competent *E. coli* cells. The correct assembly of the vectors was checked with pSNW2 flank for and pSNW2 flank rev primers. The confirmation of the targeted insertion of the construct into the *C. thermodepolymerans* chromosome was conducted using the primers 8585_Gen1 for, 8585_Gen2 rev, 14855_Gen1 for, 14855_Gen2 rev, 14025_Gen1 for, 14025_Gen2 rev for IS481_08585, IS481_14855 and IS481_14025 RE, respectively. The mentioned primers were used in combination with the primers pSNW2 flank for and pSNW2 flank rev, which annealed to the plasmid backbone. The following steps of obtaining the mutants were the same as described for the *phaC* gene. The presence of the gene deletion was confirmed with colony PCR using 8585_Gen1 for, 8585_Gen2 rev, 14855_Gen1 for, 14855_Gen2 rev, 14025_Gen1 for, 14025_Gen2 rev for IS481_08585, IS481_14855 and IS481_14025 RE, respectively. Sanger sequencing was also performed to validate the deletion of RE genes.

The vector pAI012_HA2 (Table 1) used for the *phaC* gene deletion was constructed based on the pCJ012 plasmid backbone (Johnson and Beckham, 2015) that contained a *sacB* gene originating from *B. subtilis*. *In vivo* cloning was performed to replace the PMB1 origin of replication in pCJ012 plasmid with R6K from the pSNW2 plasmid. Amplification of R6K origin was done from pSNW2 plasmid using R6K for new and R6K rev primers. The amplification of the pCJ012 backbone was conducted with Pl for new and Pl rev new pair of primers. The R6K and pCJ012 purified PCR products were digested with *Dpn*I to get rid of the plasmid templates. *E. coli* One Shot™ PIR1 (Invitrogen) competent cells were transformed with the resulting R6K_pCJ012 plasmid. Cloning was verified by PCR using Ori check for and Ori check rev primers and was further confirmed by Plasmidsaurus (U.S.A.) sequencing. The fused homologous arms for the recombination in *C. thermodepolymerans* were amplified with the primers HAN1 for, HR1 rev, HR2 for, HAN2 rev, HA1 for, and HA2_HindIII rev. The homologous sequence was introduced into the vector R6K_pCJ012 via restriction cloning using *Eco*RI and *Hind*III. The ligation mix was transformed into DH5α λπ chemically competent *E. coli* cells using the heat-shock transformation. The assembly of the pAI012_HA2 vector and the right integration into the bacterial chromosome were confirmed using the primers pCJ012 flank for, pCJ012 flank rev and pCJ012 gen flank for, pCJ012 gen flank rev, correspondingly.

The vector pAI_ThSacB was built based on the pSNW2_HA2 backbone, where the *gfp* gene was substituted for the *sacB* gene (GenBank: ACI15886.1) from *B. licheniformis*. The gene was codon-optimized and used with its native ribosome binding site (RBS). The primers ThSacB for, ThSacB rev and pSNW2_sac for, pSNW2_sac rev were used for the amplification of the *sacB* gene and the pSNW2_HA2 backbone, correspondingly. The cloning was performed via NEBuilder (New England Biolabs) assembly according to the manufactureŕs protocol. The ligation mix was transformed into CC118 λπ chemically competent *E. coli* cells using the heat-shock transformation. The correctness of the assembly was confirmed by Sanger sequencing with the primers SacB pl ch for and SacB pl ch rev.

Vectors pAI012_HA2 or pAI_ThSacB were introduced into *C. thermodepolymerans* via conjugation. Transformants were cultivated at 37 for 48 h on LB plates. Selection of the transformants was performed at 50°C on LB agar plates with Km. The pAI012_HA2 clones were passaged three times in LB with 10 % sucrose for 24 h at 37 °C, while pAI_ThSacB isolates were cultivated at 50 °C. Transformants were spread on LB agar plates with 10 % sucrose to obtain single colonies, which were further cultured at 37 °C (pAI012_HA2 transformants) or 50 °C (pAI_ThSacB transformants). Single colonies were streaked on fresh LB agar plates with 10 % sucrose. Colonies were checked for the presence of the deletion by PCR using Gen1 for and Gen2 rev primers.

### 2.5 Random integration of pBAMD_gfp vector into C. thermodepolymerans chromosome

The pBAMD_*gfp* vector was constructed using restriction cloning. For this purpose, the pSEVA238_*gfp* (Table 1) and pBAMD1-6_*glf* (Martinez-Garcia *et al*., 2014; Bujdoš *et al*., 2023) plasmids were digested with *Eco*RI and *Hind*III. The *gfp* gene, along with the synthetic RBS, was ligated into the pBAMD1-6 backbone using T4 ligase (NEB) according to the standard protocol. The propagation of the pBAMD_*gfp* vector was carried out in DH5α λπ chemically competent *E. coli* cells. *E. coli* cells harboring the pBAMD_*gfp* vector were cultivated in LB medium supplemented with Gm and Amp. The isolated pBAMD_*gfp* vector was introduced into *Caldimonas* cells via electroporation and triparental mating. *Caldimonas* transformants were selected on LB agar plates containing Gm at both 37 °C and 50 °C in parallel experiments. Twenty colonies exhibiting varying fluorescence intensities were selected for further analysis. These isolates were cultivated in a 48-well plate using the Infinite® 200 PRO (Tecan) platform. The cultivation was conducted in 600 μL of LB medium at 42 °C for 24 h. The OD_600_ was measured every 15 min. Orbital shaking with a 2.5 μm amplitude was applied between measurements, and linear shaking with a 3 μm amplitude was performed for 10 s before each measurement. To prevent water condensation on the plate lid, a Triton solution (0.05 % v/v Triton X-100 in 20 % v/v ethanol) was poured onto the lid to cover the entire surface (Brewster, 2003). The lid was then dried and sterilized using UV light.

### 2.6 Production and quantification of PHA in C. thermodepolymerans strains

To verify and compare the PHA production ability of the mutant strains with the wild-type strain, several cultivations were carried out in Erlenmeyer flasks. In the first step, the culture was cultivated for 20 h in 50 mL of nutrient-rich complex medium (Nutrient Broth w/ 1% Peptone, HiMedia) at 50 °C with constant shaking (160 rpm) in 100 mL Erlenmeyer flasks. After this period, the basic mineral salt medium (MSM) was inoculated with 5 vol.% (100 mL of MSM in 250 mL Erlenmeyer flasks). The composition of the MSM was 9.0 g/L Na_2_HPO_4_ · 12 H_2_O, 1.5 g/L KH_2_PO_4_, 1.0 g/L NH_4_Cl, 0.2 g/L MgSO_4_ · 7 H_2_O, 0.02 g/L CaCl_2_ · 2 H_2_O, 0.0012 g/L Fe^(III)^NH_4_citrate, 0.5 g/L yeast extract, 1 mL/L of microelements solution (50.0 g/L EDTA, 13.8 g/L FeCl_3_ · 6 H_2_O, 0.84 g/L ZnCl_2_, 0.13 g/L CuCl_2_ · 2 H_2_O, 0.1 g/L CoCl_2_ · 6 H_2_O, 0.016 g/L MnCl_2_ · 6 H_2_O, 0.1 g/L H_3_BO_3_, dissolved in distilled water) and carbon source – xylose (20 g/L) was added. After 72 h of cultivation (50 °C, 160 rpm), bacterial cells were harvested (10 ml of bacterial suspension per sample) by centrifugation (3,700 × g, 5 min.) washed with distilled water, centrifuged again, the supernatant was discarded, the cell pellet was dried to constant weight. The biomass was determined gravimetrically in at least three biological replicates. The dried biomass was forwarded for sample preparation for analysis and quantification of PHA. To approximately 8 mg of dried biomass was added 1 mL of chloroform and 0.8 mL of a solution containing 15 % sulfuric acid dissolved in methanol with benzoic acid (internal standard applied at concentration of 5 mg/mL). This mixture was left for 3 h at 94 °C in a thermoblock, at which time methanolysis of the PHA polymer occurred. Subsequently, the content of the vial was added to 0.5 mL NaOH and shaken vigorously. After the phases had settled, 50 µl of organic phase was transferred into 900 µl of isopropyl alcohol. The samples thus prepared were analyzed using Trace GC Ultra with flame ionization detector as reported previously (Obruca *et al*., 2013).

### 2.7 Determination of cell numbers by flow cytometry

Cell numbers were determined by flow cytometry (Cytek Biosciences, NL-2000). Bacterial suspensions were diluted as required. In case of *C. thermodepolymerans* WT and Δ*phaC* mutant strain, 1000× and 100× dilutions, respectively, were made using decimal dilution. The resulting number of cells was calculated from the number of 10,000 events (ROI was selected from scategram FSC vs. SSC) and the volume of diluted bacterial suspension used. The experiment was performed in triplicate.

### 2.8 Statistical analyses

The number of repeated experiments or biological replicates is specified in figure and table legends. The mean values and corresponding standard deviations are presented. When appropriate, data were treated with a two-tailed Student’s t-test in Microsoft Office Excel 2013 (Microsoft) and confidence intervals were calculated for the given parameters to test a statistically significant difference in means between two experimental datasets.

## 3. Results and discussion

The process of domestication of a wild-type bacterium into a new chassis strain for biotechnology and synthetic biology endeavors involves several fundamental stages (Fig. 1) (Obruča *et al*., 2022; Pal *et al*., 2024). In previous studies of *C. thermodepolymerans,* we described the whole genome sequence, antibiotic sensitivity and presence of R-M systems (Musilova *et al*., 2021, 2023). In this work, we focus on several remaining steps in stages I and II.

**Fig. 1.**
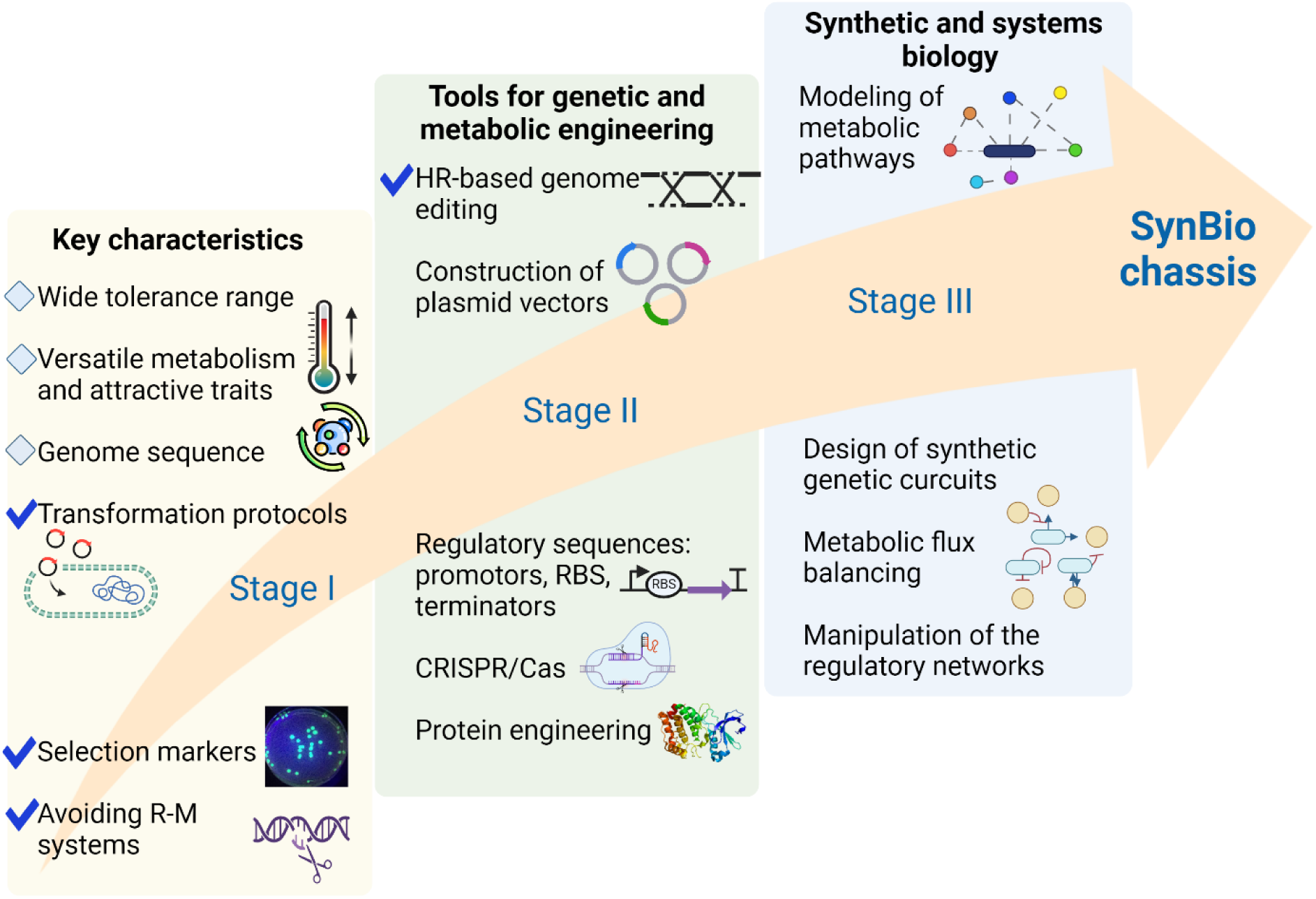
Roadmap of SynBio chassis development with three fundamental stages. The rhombuses represent the results from previous research, while the check marks indicate the accomplishments of this study.

### 3.1 Growth tests at different temperatures and optimization of the electroporation protocol

The growth of *C. thermodepolymerans* in rich LB medium was assessed at both 50 °C and 37 °C (Fig. 2). The bacterium demonstrated robust growth at both temperatures, indicating its facultative thermophilic nature. At 37 °C, the growth exhibited a biphasic pattern, characterized by a shorter initial exponential phase followed by a longer second exponential phase. Notably, the maximum growth rate of approximately 0.5 h^-1^ at 50 °C and 0.2 at 37 °C (during the second exponential phase) was achieved twice as quickly at the optimal temperature of 50 °C (within 4 h) compared to 37 °C (within 8 h). Additionally, the highest OD_600_ recorded was slightly higher at 50 °C (∼ 2.4) than at 37 °C (∼ 2.1). The ability of *C. thermodepolymerans* to grow at 37 °C offers flexibility in manipulation, which is advantageous for genetic engineering.

**Fig. 2.**
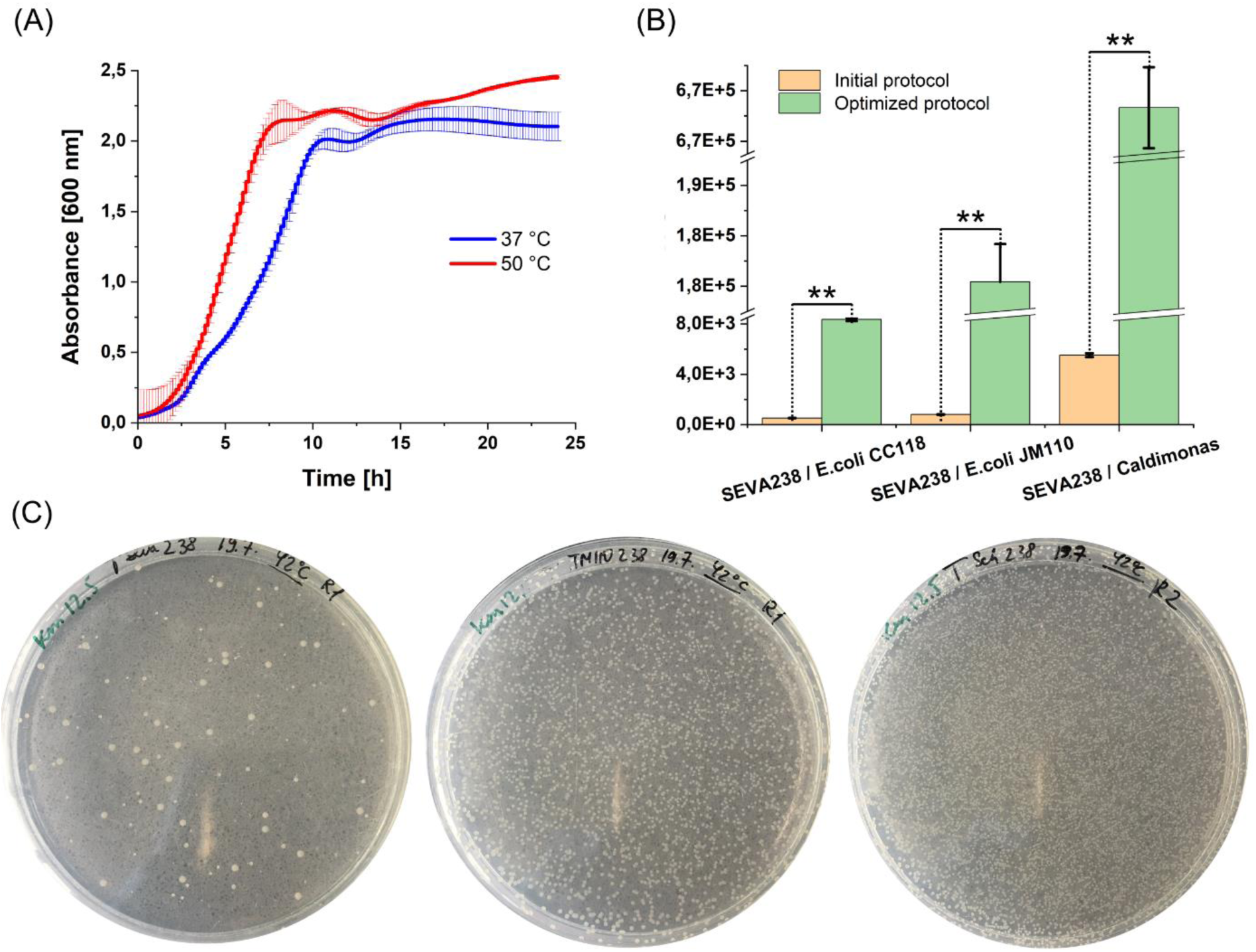
The growth and transformation efficiency of *C. thermodepolymerans*. (A) Growth of *C. thermodepolymerans* at different temperatures. (B, C) Transformation efficiency depending on the electroporation protocol (B) and plasmid isolation source (B and C). Columns in (B) represent means ± standard deviations calculated from three (n = 3) independent experiments. CFU is the abbreviation for colony forming unit. Asterisks (**) denote the significance of the difference between the two means at p < 0.01 calculated using two-tailed Student *t* test (p values from left to right = 4.6 x 10^-8^, 4.8 x 10^-8^, 3.7 x 10^-10^). Plates (C) show *C. thermodepolymerans* pSEVA238 transformants using plasmids isolated from different sources, listed from left to right: *E. coli* CC118, *E. coli* JM110, and *C. thermodepolymerans* DSM 15344. Conditions of the experiment are described in the Methods section.

The cellular uptake of foreign DNA is one of the key prerequisites for a successful transformation process and entire genome-editing protocol. Electroporation is routinely used to transform thermophilic bacteria as heat shock is not an option and conjugation protocols are established for mesophilic hosts (Kong *et al*., 2021, Averhoff and Friedrich, 2003). To assess the efficacy of electroporation of *C. thermodepolymerans*, plasmid pSEVA238 (medium-copy number) was selected from the Standard European Vector Architecture collection (Martínez-García *et al*., 2023) This plasmid harbors a broad-host-range origin of replication and a Km resistance cassette. The sensitivity of *C. thermodepolymerans* to Km, streptomycin, Gm, tetracycline, and chloramphenicol was examined previously (Musilova *et al*., 2023). The Km was confirmed as a suitable antibiotic with a minimal inhibitory concentration of 12.5 µg/mL for selection purposes in *C. thermodepolymerans.* The electrotransformation protocol previously established for the related organism *C. manganoxidans* served as the initial blueprint (Arai *et al*., 2022). According to this protocol, bacteria were cultivated at an optimal growth temperature of 50 °C, followed by competent cell preparation. The prepared competent cells were used immediately. However, when applied to *C. thermodepolymerans*, this protocol yielded suboptimal results (Fig. 2). Moreover, competent cells prepared under this protocol could not be frozen because they were not viable upon thawing. To address this, the growth temperature for competent cell preparation was reduced from 50 °C to 37 °C. This modification led to a substantial improvement in transformation efficiency and enabled the freezing of cells without compromising viability. The reduction in growth temperature is aimed to create an optimal physiological state in the bacterial cells that enhances their ability to take up foreign DNA during the transformation process (Zhou *et al*., 2018). Consequently, under the revised conditions, the transformation efficiency of *C. thermodepolymerans* increased by up to two orders of magnitude and ranged from 8.3 x 10^3^ to 6.7 x 10^5^ CFUs depending on the source strain of isolated plasmid DNA (Fig. 2B and C). This result underscores the importance of optimization for obtaining robust and reproducible electroporation protocol.

### 3.2 Influence of restriction-modification systems on transformation efficiency

Bacteria possess various defense mechanisms, such as native restriction-modification systems (R-M), to protect against foreign DNA intrusion. R-M systems present a significant barrier to genetic manipulation, particularly in genetically intractable bacteria. This challenge has necessitated the development of methods to overcome R-M barriers, enabling advancements in bacterial genetics and synthetic biology (Sánchez-Romero *et al*., 2015).

Key strategies include: (i) altering the target DNA sequences to avoid recognition by the restriction enzymes (Hu *et al*., 2023), (ii) introducing genes encoding methyltransferases into host bacteria (Ren *et al*., 2022), (iii) using certain proteins that inhibit restriction enzymes (Kudryavtseva *et al*., 2023), (iv) using bacterial strains that have been genetically engineered to lack functional restriction endonucleases (Ishikawa and Hori, 2024), and (v) choosing bacterial strains that are naturally deficient in specific R-M systems or that have evolved to have less restrictive R-M activity (Suzuki, 2012).

The type strain of *C. thermodepolymerans* DSM15344 displays one type I and two type II R-M systems (Musilova *et al*., 2023). To study their influence on transformation efficiency, *C. thermodepolymerans* was electroporated with the same plasmid pSEVA238 isolated from three source bacterial strains and, therefore, having different methylation patterns. These were: 1) routinely used cloning strain of *E. coli* CC118, 2) methyltransferase-deficient *E. coli* JM110, and 3) *C. thermodepolymerans* itself (Fig. 2B and C). The lowest number of transformants − 8.3 x 10^3^ CFUs / 1 µg of plasmid DNA - was observed when using *E. coli* CC118 as the isolation source. Electroporation with pSEVA238 from *E. coli* JM110 resulted in a significantly higher transformation efficiency (1.8 x 10^5^ CFUs / 1 µg of plasmid DNA). The highest efficiency - 6.7 x 10^5^ CFUs / 1 µg of plasmid DNA - was obtained using *C. thermodepolymerans* as the plasmid isolation source, giving rise to 3.7-fold more transformants than *E. coli* JM110 used as a plasmid isolation source and two orders of magnitude more than *E. coli* CC118 used as a plasmid isolation source. Employing an optimized electroporation protocol consistently yielded better results across all cases compared to the previously described protocol (Fig. 2B and C) (Arai *et al*., 2022). These findings indicate the significant impact of R-M systems in *C. thermodepolymerans* on transformation efficiency. Utilizing methyltransferase-deficient cloning strains, such as *E. coli* JM110, substantially reduces foreign DNA recognition and digestion by restriction endonucleases (Kolek *et al*., 2016).

### 3.3 Initial testing of three genome editing protocols in C. thermodepolymerans

The PHA biosynthetic pathway in *C. thermodepolymerans* consists of three enzymes (acetyl-CoA C-acetyltransferase IS481_08635, acetoacetyl-CoA reductase IS481_08640, polyhydroxyalkanoate synthase IS481_08630) that convert acetyl-CoA into PHB. We aimed to use a homologous recombination-based protocol for the deletion of the *phaC* gene (IS481_08630) and thus verify the essentiality of this gene for PHB biosynthesis. The process, in general, employs a non-replicative plasmid vector (Fig. 3) (Hmelo et al., 2015). In the initial phase, the suicide vector integrates into the bacterial chromosome through the first crossover event. The selection of the single crossover recombinants is based on antibiotic resistance gene in plasmid backbone. Moving on to the second phase, double crossovers should arise during a subsequent infrequent homologous recombination event (Fig. 4).

**Fig. 3.**
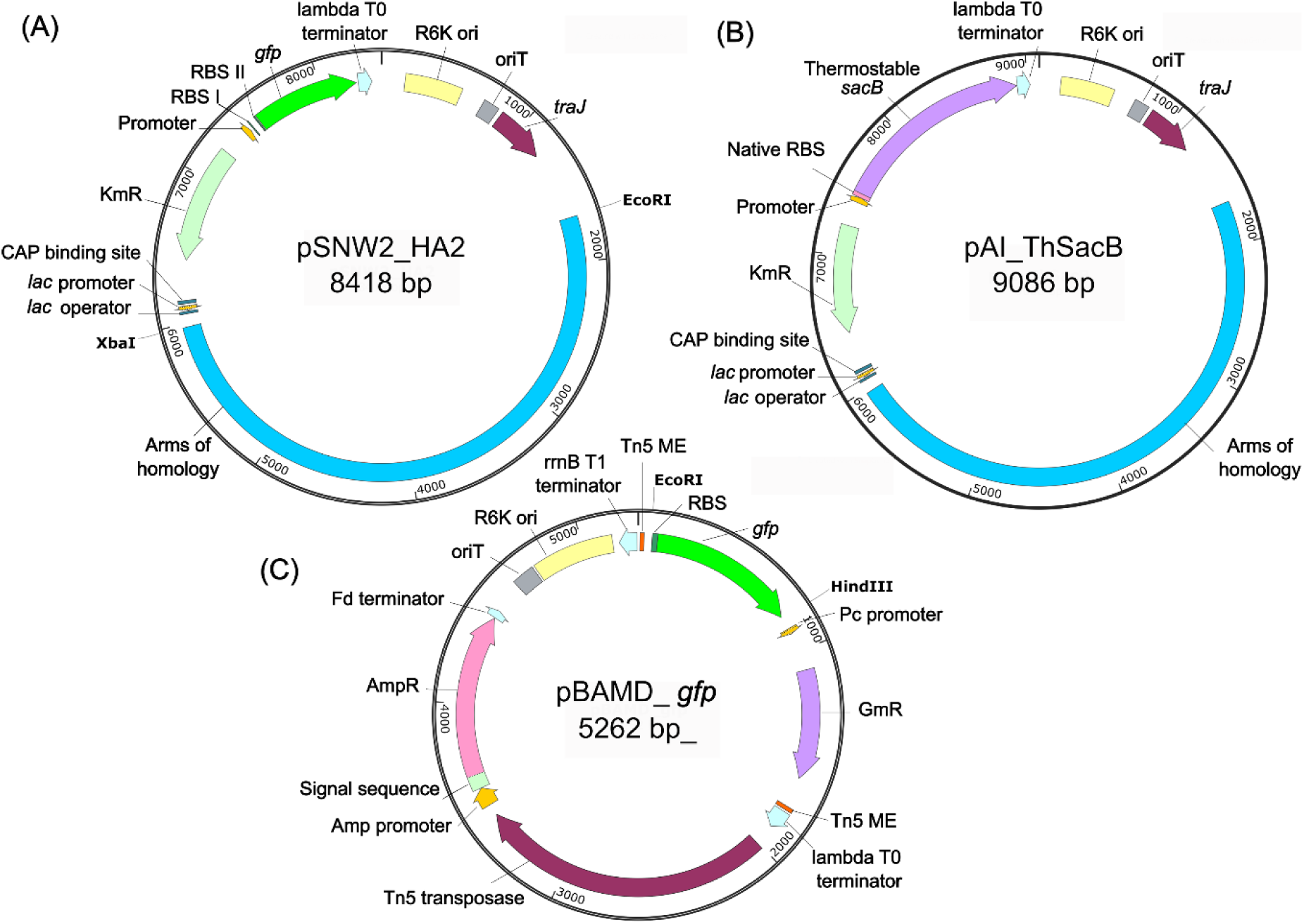
Maps of the plasmids used for homologous-recombination and transposition-based protocols. Abbreviations: Tn5 ME - hyperactive mosaic end for Tn5 transposase recognition, GmR - Gm acetyltransferase, AmpR - β-lactamase, oriT - origin of transfer, TraJ - oriT-recognizing protein, KmR-aminoglycoside phosphotransferase, msfGFP - monomeric superfolder green fluorescent protein. The plasmid maps were prepared using SnapGene Viewer.

**Fig. 4.**
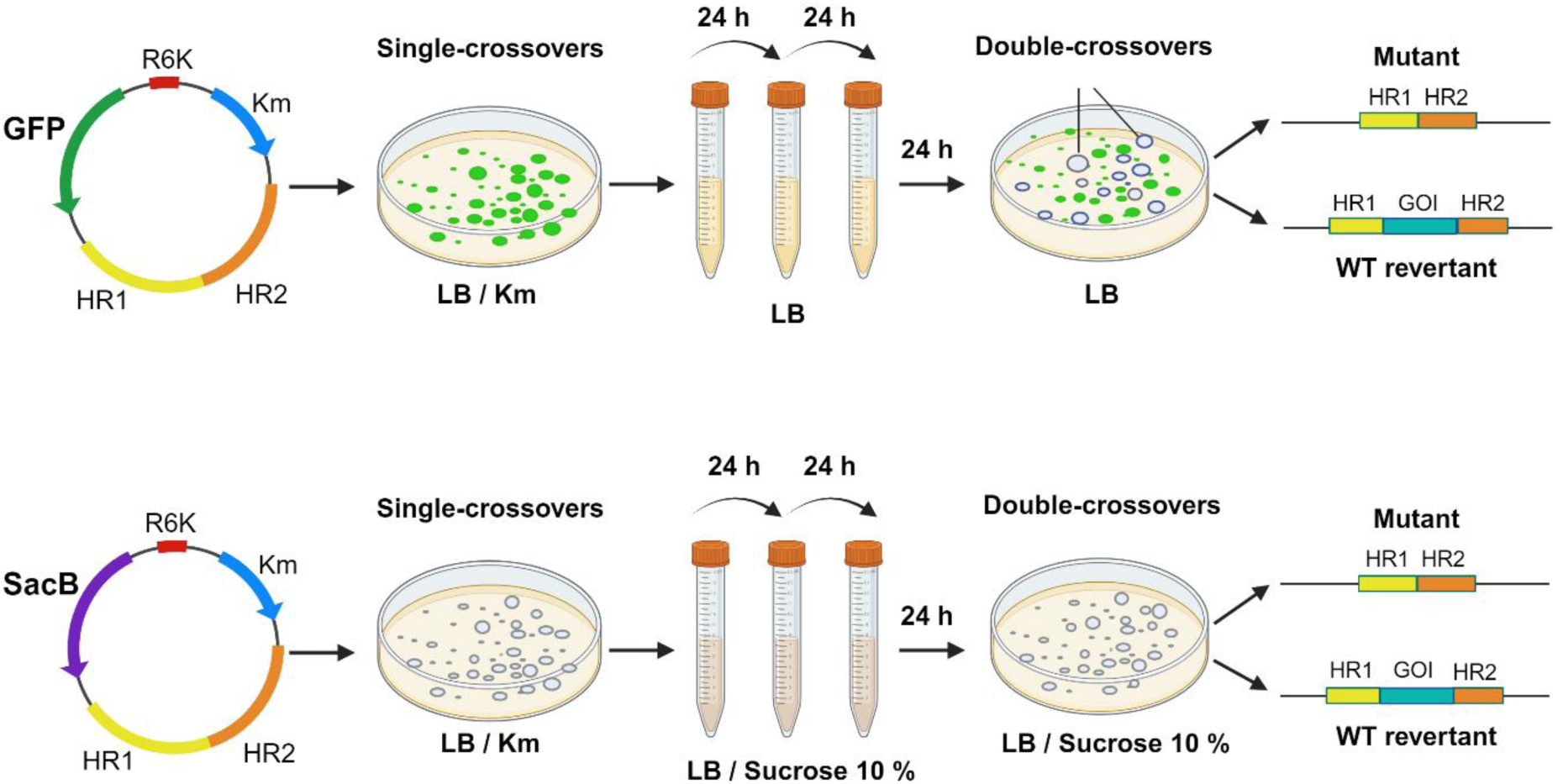
Genome editing of *C. thermodepolymerans* using GFP- and SacB-based homologous recombination protocols for targeted gene deletion.

To conduct the deletion, we constructed suicide vectors pSNW2_HA2 and pAI012_HA2 (Fig. 3), which differed by the selection marker genes – monomeric superfolder *gfp* or *sacB*, respectively. These constructs also included the Km resistance gene and a suicide λpir-dependent origin of replication, R6K, ensuring replication in *E. coli* but preventing replication in *Caldimonas* (Johnson and Beckham, 2015; Volke *et al*., 2020). The deletion vector pSNW2_HA2 was constructed based on pSNW2, a derivative of vector pGNW2 (Volke *et al*., 2020). The stability and functionality of the GFP protein at 50 °C were previously validated (Millgaard et al., 2023). The vector pAI012_HA2 was derived from pCJ012, a plasmid designed for the deletion of the *lldD* gene encoding L-lactate dehydrogenase in *P. putida* through homologous recombination and *sacB*-mediated sucrose counter-selection (Johnson and Beckham, 2015). The deletion vectors were introduced into *C. thermodepolymerans* through electroporation. No pSNW2_HA2 single-crossover recombinants grew at 50°C. After 48 h of growth at lower temperature of 42 °C, colonies appeared, although they did not exhibit fluorescence and were very small. This indicated that the nonspecific growth on the initial Km LB plates was possibly due to trace amounts of the vector and antibiotic resistance protein that took hold in the cells but was gradually diluted over subsequent bacterial generations. Although the number of biological replicates was increased, only two fluorescent colonies were successfully recovered on agar plates supplied with Km (Fig. 5B). These colonies were further streaked onto the second LB plates with Km, displaying growth at both 42°C and 50°C. The integration of the vector into the bacterial chromosome was confirmed with PCR. Subsequently, two isolates with confirmed integration underwent three passages in liquid LB at 42°C and 50°C. They were then spread on LB plates and single clones that lost fluorescence were examined for the presence of the *phaC* gene deletion. Despite checking over three hundred clones, all proved to be revertants to the wild type, indicating the need for further optimization in the experimental design.

**Fig. 5.**
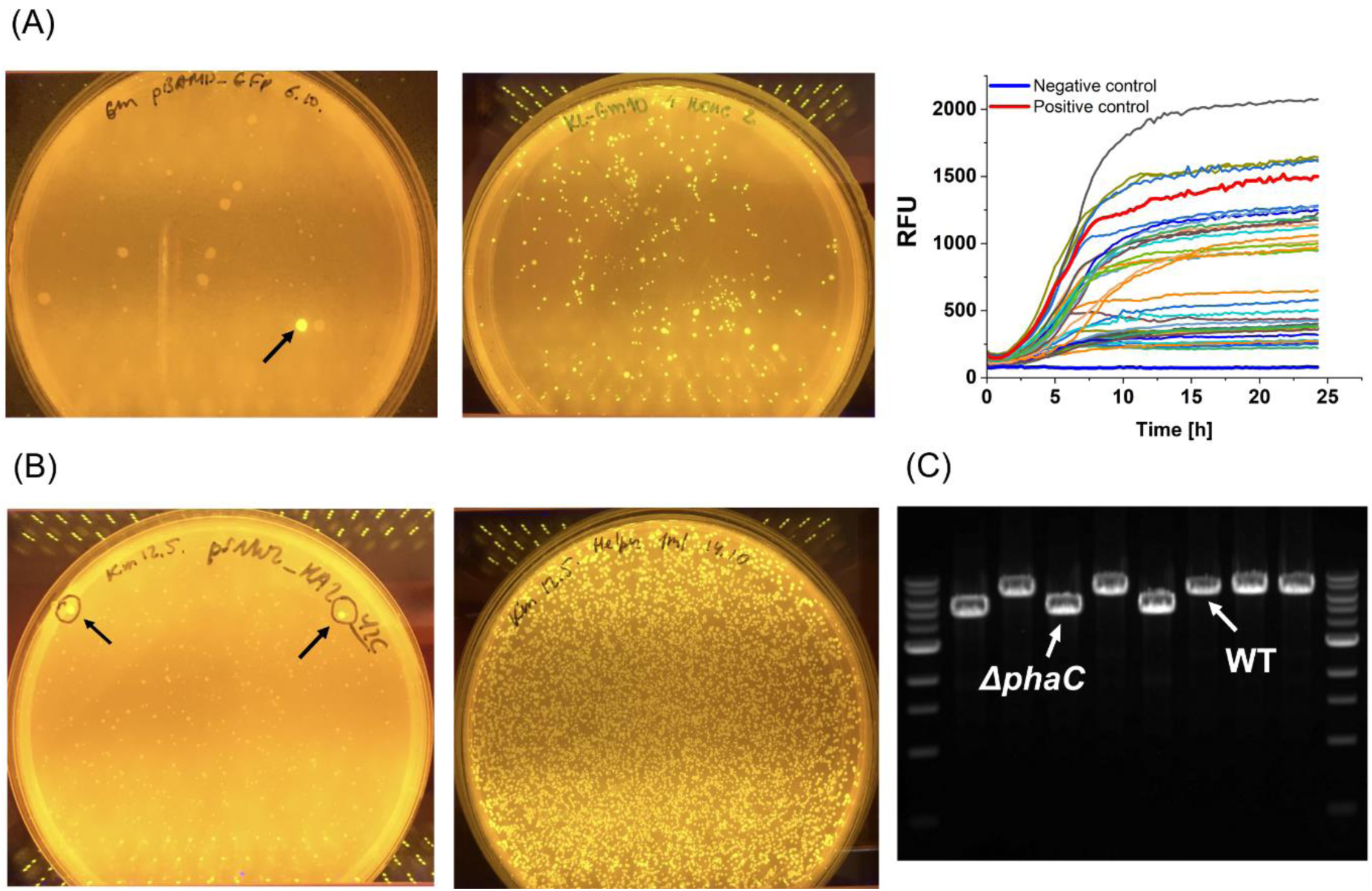
Genome editing of *C. thermodepolymerans* using transposition-based and homologous-recombination based protocols. (A) Transformation of *C. thermodepolymerans* with pBAMD_*gfp* using electroporation (left plate) or conjugation (right plate), and testing of fluorescence intensity of isolated clones in a 48-well plate liquid culture, RFU - relative fluorescence units. (B) Transformation of *C. thermodepolymerans* with pSNW2_HA2 using electroporation (left plate) or conjugation (right plate). (C) PCR verification of double crossovers during the *phaC* gene deletion (the ∼ 6,300 bp products corresponded to Δ*phaC* mutants, ∼ 4,600 bp products – to wild-type revertants, 1 kb DNA ladder from NEB was used). Black arrows in (A) and (B) indicate the single crossover transformants obtained after electroporation.

After unsuccessful attempts to prepare the *phaC* deletion using the GFP selection, the other system based on SacB counterselection was utilized. Gene *sacB* codes for the levansucrase. This enzyme converts sucrose into levans, leading to a toxic effect in Gram-negative bacteria (Ried and Collmer, 1987). Vector pAI012_HA2 was electroporated into *Caldimonas*. The selection of pAI012_HA2 single-crossover recombinants was performed at 42 °C and 50 °C in parallel. However, only two colonies were obtained after 48 h of incubation at 42 °C on LB plates with Km. Then the targeted integration into the chromosome was confirmed followed by a counterselection process in LB medium supplied with 10 % sucrose at 42°C. We tested 5, 10 and 15 % sucrose concentrations and identified that 10 % sucrose was maximum that type *C. thermodepolymerans* strain could withstand. In medium with 15 % sucrose the bacterial growth was absent. The counterselection on sucrose was conducted at 42 °C because *sacB* originated from a mesophilic bacterium *B. subtilis* (Zhang *et al*., 2021). Over two hundred colonies were checked after counterselection, but none fully lost the integration cassette. The majority of the isolates remained single crossovers, some represented the mix of the WT of mutant genotypes.

The notably low occurrence of the initial recombination event pointed to the possibly low level of recombination frequency in *C. thermodepolymerans*. Therefore, we decided to test a Tn5-based mini-transposon system pBAMD for random integration of a gene of interest into the genome that did not encompass a homologous recombination step (Martinez-Garcia *et al*., 2014; Bujdoš *et al*., 2023). *C. thermodepolymerans* was electroporated with a constructed pBAMD_*gfp* vector to: i) check if the other mechanism of integration into the chromosome worked more efficiently than HR; ii) to prepare the library of the clones with *gfp* under the innate *C. thermodepolymerans* promoters (transcriptional fusions) with varying strengths. Nevertheless, we again failed to obtain clones with visible green fluorescence even after ten independent transformations, except for one fluorescent colony, which was recovered (Fig. 5A). This result together with the previous difficulties in obtaining cointegrates via homologous recombination suggested that the challenge might lie in the insufficient entry of plasmid DNA into the cell.

### 3.4 Conjugation as an alternative transformation method for C. thermodepolymerans

An alternative method for bacterial transformation is conjugation, often used in the form of triparental mating (TPM) (Glenn *et al*., 1992, Samperio *et al*., 2021). Compared to electroporation, TPM is generally more time-demanding and three different strains are needed (Lin and Xu, 2013). It is less commonly used for thermophilic bacteria due to temperature incompatibility between the donor, helper, and acceptor strains and thermal instability of the conjugation machinery and other genetic elements from mesophilic bacteria (Averhoff and Friedrich, 2003). Consequently, we did not initially consider TPM as the most suitable method for transforming *Caldimonas*. However, the ability of *C. thermodepolymerans* to grow at mesophilic temperatures allowed us to use 37°C for the first step of conjugation and the transfer of suicide plasmid vector using *E. coli* donor and helper strains and 50°C for the selection of *Caldimonas* recombinants with chromosomal integration. We first tested the TPM protocol for the random integration of *gfp* through the Tn5-based transposition (pBAMD1-4_*gfp*). During the initial stage of conjugation, the donor (*E. coli* CC118), helper (*E. coli* HB101), and recipient (*C. thermodepolymerans* DSM 15344) bacterial strains were cultured at 37°C for two days. Subsequently, for the selection of transconjugants with *gfp*, the strain mixture was streaked on LB agar plates with Gm and incubated at 50°C. Unlike electroporation, the conjugation process resulted in hundreds of fluorescent colonies (Fig. 5A). Twenty isolates were picked, grown in liquid LB medium in microtiter plate format, and tested for GFP fluorescence intensity (Fig. 5A). The GFP fluorescence differed significantly between the individual clones pointing to the variability of the promoter strength under which the *gfp* gene was randomly integrated. The isolates were frozen for future analysis and characterization of the promoters.

### 3.5 Deletion of phaC and genes of three restriction endonucleases with modified genome-editing protocols

Following the successful application of TPM for random genome integration, this method was chosen over electroporation for the deletion of the *phaC* gene using the pSNW2_HA2 vector. After the conjugation, thousands of potential single crossover recombinants with visible GFP fluorescence were recovered on Km plates (Fig. 5B). Sixteen colonies were screened using PCR to verify the targeted integration of the deletion vector. Of these, seven isolates (∼ 43%) demonstrated full-size targeted integration into the chromosome. Individual seven cointegrates were passaged three times in liquid LB at 50°C, with each passage lasting 24 hours. Subsequently, samples of cultures were plated onto LB plates for single colony isolation and incubated at 50°C. Single colonies lacking fluorescence were streaked on fresh plates and analyzed via PCR to confirm the deletion. Over 5,000 colonies were obtained on LB plates, with 16 (∼0.3%) identified as double crossovers. Among the 16 analyzed isolates, 6 (∼37%) were confirmed as deletion mutants, while 10 (∼ 62%) reverted to the wild type (Fig. 5C). PCR analysis for the presence of the *phaC* gene and Sanger sequencing of the target region confirmed the seamless deletion of the gene in all six clones.

To validate the GFP-based protocol and potentially further enhance the transformation efficiency and plasmid maintenance in *C. thermodepolymerans*, we decided to delete genes of three previously identified restriction endonucleases: IS481_08585, IS481_14855, and IS481_14025 (Musilova et al., 2023). The deletion procedure was the same as described for the *phaC* gene. The protocol involved conjugation followed by Km and GFP fluorescence selection for getting single and double crossovers, respectively. Following TPM, sixteen colonies were screened for the cointegration event. All examined colonies were identified as single crossovers with integrated deletion plasmid. These single crossovers were then passaged in liquid LB, and isolates that lost fluorescence were tested for the presence of the deletion with PCR and Sanger sequencing. For IS481_08585, five double crossovers (∼0.13 %) lacking GFP fluorescence were obtained from 3,810 colonies on plates, with three of these (∼60 %) being deletion mutants. For IS481_14855, four double crossovers (∼0.09 %) were obtained from 4,662 colonies, with one (25 %) being a deletion mutant. For IS481_14025, six double crossovers (∼0.13 %) were obtained from 4,550 colonies, with two (33.3 %) being deletion mutants. Thus, the rate of the second recombination event was low, ranging from 0.09 % to 0.3 %.

The transformation efficiency of the constructed triple-deletion *C. thermodepolymerans* strain designated AI01, lacking the RE, was compared to the WT strain using standard plasmids pSEVA238 and pSEVA631 from the SEVA database (Fig. 6A, Table 1). The plasmids were introduced into *Caldimonas* via electroporation. Mutant stain AI01 gave fifty times more pSEVA238 transformants and forty times more pSEVA631 transformants than the WT strain at 37 °C. At 50 °C, only pSEVA631 transformants could be recovered, although at a considerably lower number than at 37 °C (25 times less for WT, 43 times less for AI01 strain). At the same time, the number of pSEVA631 transformants generated by the AI01 strain at 50 °C was still twenty times higher than that of the WT strain. The failure of *Caldimonas* transformants to grow at 50 °C indicates that the pBBR1 origin of replication is thermally unstable. The broad-host-range plasmid BBR1 was initially isolated from *Bordetella bronchiseptica*, a bacterium that grows optimally at temperatures between 35 and 37 °C (Antoine and Locht, 1992), and is not optimized for use in thermophilic bacteria. For thermophilic bacteria, specialized origins of replication are required to ensure that plasmids can replicate efficiently at the high temperatures (Olson and Lynd, 2012). But most shuttle vectors with thermostable origin of replication have been developed for Gram-positive bacteria (Chung *et al*., 2013 Groom *et al*., 2016). Since the BBR1 replicon is likely almost not functional in *C. thermodepolymerans* at 50 °C it is essential to identify a suitable thermostable origin of replication that operates effectively at high temperatures. In a parallel study, we focus on identifying native plasmids in closely related thermophilic bacteria and analyzing their replicons.

**Fig. 6.**
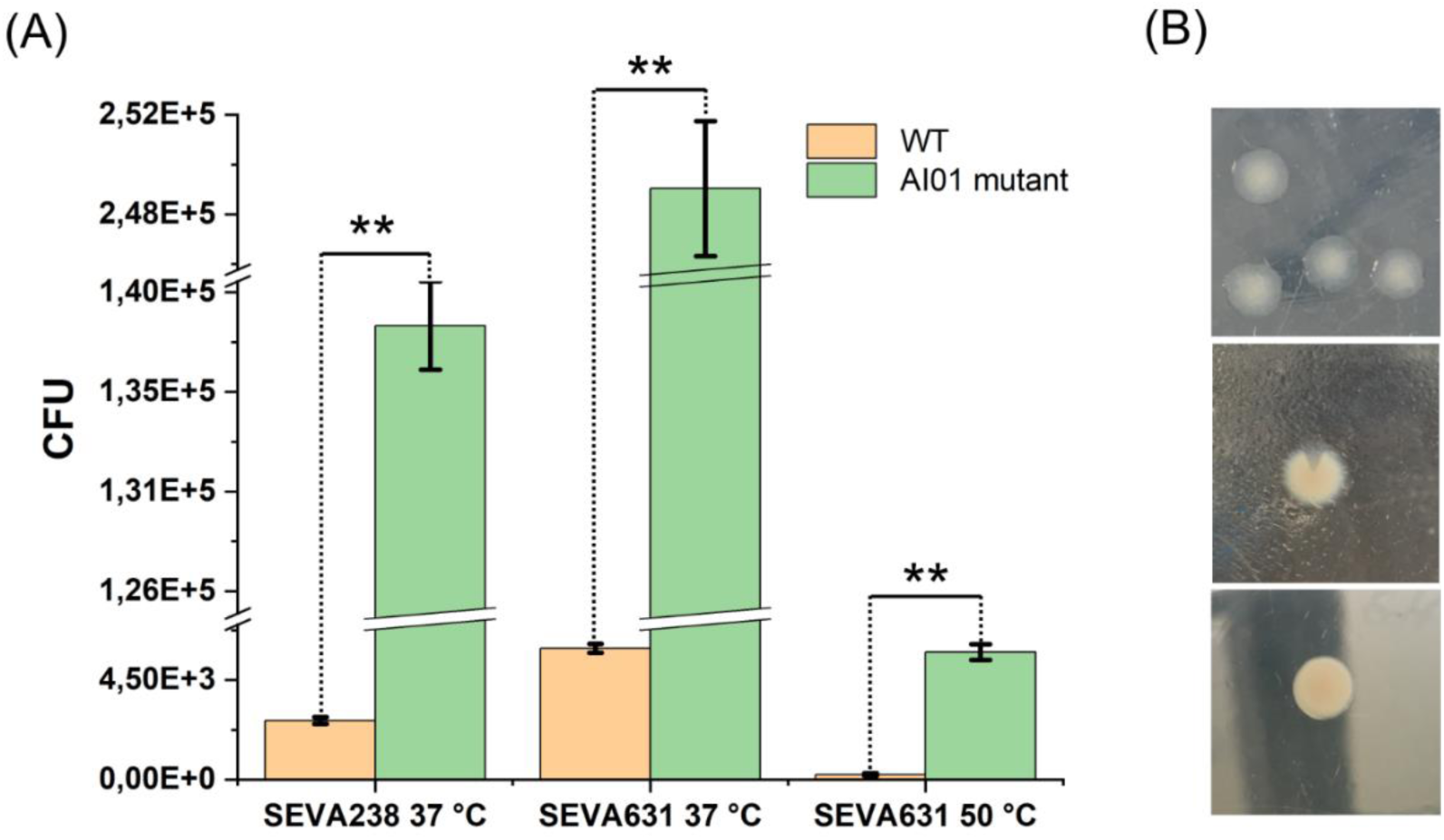
Application of the triple-deletion *C. thermodepolymerans* strain AI01. (A) Transformation efficiency of AI01 strain transformed with pSEVA238 or pSEVA631. Columns in (A) represent means ± standard deviations calculated from three (n = 3) independent experiments. Asterisks (**) denote the significance of the difference between the two means at p < 0.01 calculated using two-tailed Student *t* test (p values from left to right = 3.2 x 10^-8^, 1.2 x 10^-12^, 1.1 x 10^-5^). CFU is the abbreviation for colony forming unit. (B) Morphologies of *C. thermodepolymerans* AI01 colonies observed during the search for double crossovers using SacB-based protocol. Transparent portions of the colonies contained double crossover recombinants, while the non-transparent colonies were a mix of single and double crossovers.

### 3.6 New RE deficient AI01 strain enables the use of a thermostable SacB counterselection protocol for phaC gene deletion

In the previous section, we validated the GFP-selection-based protocol as a suitable methodology for targeted genome editing in *C. thermodepolymerans*. While GFP facilitates visual monitoring during the deletion procedure, it does not provide pressure for the selection of mutant genotypes. Therefore, in the next step, we optimized also the system utilizing SacB counterselection for genome editing in *C. thermodepolymerans*. Vector pAI012_HA2, used for deleting the *phaC* gene in this study, carried the *sacB* gene from the mesophilic bacterium *B. subtilis*. However, the functionality of the mesophilic SacB enzyme diminishes with increasing temperature (Sangiliyandi and Gunasekaran, 1998). To address this issue, we replaced the gene with a thermostable variant sourced from the thermophilic bacterium *B. licheniformis* (Nakapong et al., 2013). A thermostable SacB enzyme was identified through a literature search and subsequently validated using EnzymeMiner software (Hon *et al*., 2020). We substituted the *gfp* gene in pSNW2_*gfp* with the thermostable *sacB* resulting in the pAI_ThSacB construct. This vector was electroporated into *C. thermodepolymerans* AI01 competent cells. All steps following electroporation were conducted at an optimal temperature of 50 °C.

Sixteen examined colonies, selected for single crossovers on Km plates, were confirmed as cointegrates. Interestingly, after 72 h of cultivation on agar plates, the colonies exhibited three different phenotypes (Fig. 6B): i) colonies with an opaque center and transparent edges, ii) opaque colonies with one triangle-shaped transparent sector, iii) entirely opaque colonies. PCR analysis revealed that the transparent portions of the colonies contained double crossover recombinants, while the non-transparent colonies were a mix of single and double crossovers. Of the 16 colonies tested, 5 (∼ 31 %) were identified as pure double crossovers, while 9 (∼ 56 %) showed a mix of single and double crossovers. Among the double crossovers, approximately 80 % were mutants, and 20 % were WT revertants. The primary advantage of the GFP-based system is the ability to visually identify successful recombinants. Fluorescent colonies indicate effective plasmid integration into the chromosome, while the loss of fluorescence during subsequent cultivation signals the occurrence of double crossovers (Fig. 4). Our study revealed that this approach encounters challenges with very low frequencies of double crossovers, ranging from 0.08 % to 0.3 %. In contrast, the SacB system provides a more effective counterselection mechanism, inhibiting bacterial growth in the presence of sucrose unless the necessary recombination events occur. This simplifies the elimination of non-recombinants. By utilizing a thermostable variant of SacB from *B. licheniformis*, we optimized the system, achieving a significantly higher occurrence of double crossovers (∼31 %) at an elevated temperature of 50 °C, thereby improving its effectiveness for thermophilic bacteria. Overall, the SacB-based approach, when optimized for high temperatures, provides better efficiency for gene deletion in *C. thermodepolymerans* due to its strong counterselection mechanism.

### 3.7 Verification of PHA production by ΔphaC and AI01 mutants

The deletion of the *phaC* gene encoding PHA synthase completely disrupted the ability of *C. thermodepolymerans* DSM 15344 to synthesize PHA (Fig. 7A). The lack of PHA synthesis probably affected the fitness of the Δ*phaC* strain compared to the wild type. Flow cytometry analysis verified that the observed lower cell biomass in the culture of the mutant was caused by an order of magnitude lower number of bacterial cells compared to the wild-type culture (3.99 x 10^9^ cells/mL and 2.12 x 10^8^ cells/mL, respectively). This result highlights the importance of PHA metabolism for *C. thermodepolymerans* DSM 15344. A similar effect has been observed for *Cupriavidus necator*, where disruption of PHA metabolism through inactivation of the *phaC* gene also led to impaired growth and reduced stress robustness in the culture (Raberg et al. 2014; Obruca et al. 2017).

**Fig. 7.**
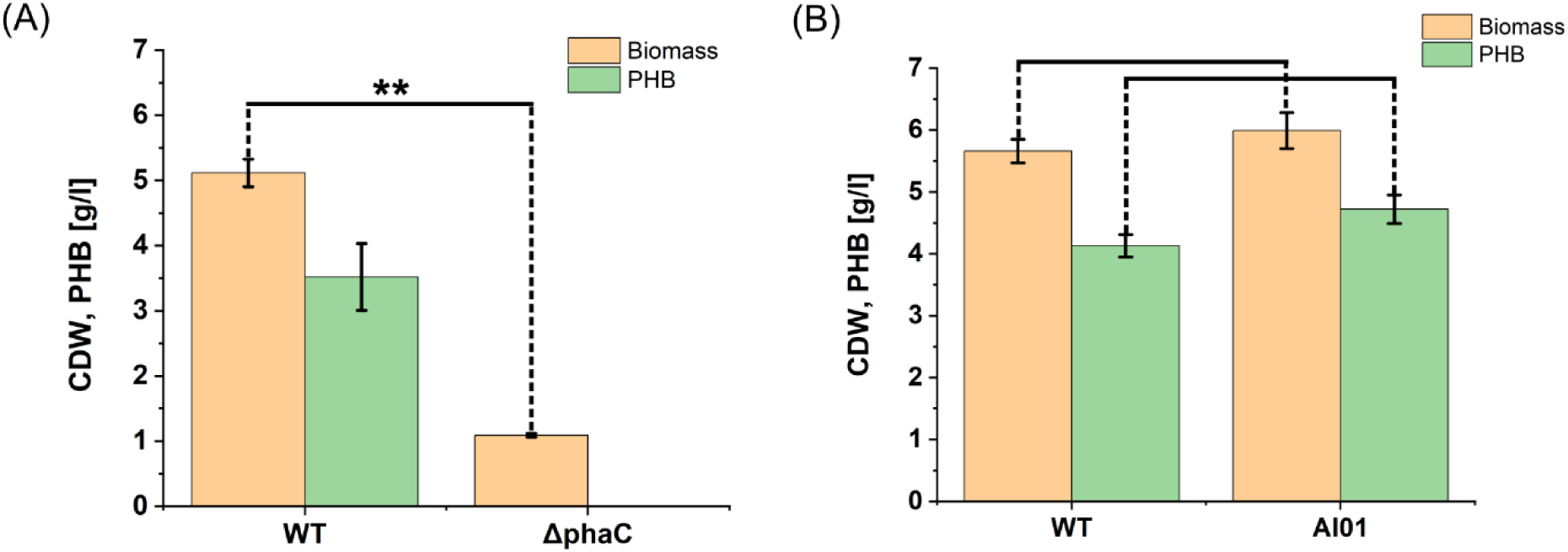
Biomass formation and PHA production by wild type of *C. thermodepolymerans* DSM 15344 and its Δ*phaC* and AI01 mutant strains. (A) The effect of deletion of *phaC* gene, (B) the effect of deletion of three endonuclease genes in *C. thermodepolymerans* AI01 from identified R-M systems. Asterisks (**) denote the significance of the difference between the two means at p < 0.01 calculated using two-tailed Student *t* test, (p-value in (A) = 1.4 x 10^-3^, in (B) for biomass p = 0.6; for PHB production p = 0.07). Columns in (A) and (B) represent means ± standard deviations calculated from two (A) and three (B) independent experiments.

In contrast, the deletion of the endonucleases from R-M systems did not affect the growth or PHA synthesis capacity of the culture, as both biomass and PHA titers were very similar in the wild-type and AI01 mutant strains (Fig. 7B). This is a promising result, indicating that *C. thermodepolymerans* AI01 can serve as a foundation for the development of a thermophilic chassis for advanced PHA synthesis.

## Conclusion

In this study we introduced homologous recombination and transposon-based systems for genome editing in Gram-negative bacterium *Caldimonas thermodepolymerans*. This organism with biotechnologically attractive properties can be considered the first model of thermophilic PHA producer. After initial difficulties, optimized transformation protocol and refined counterselection with a thermostable SacB improved the performance of tested systems. This progress underscores the importance of iterative optimization and the need for tailored approaches when working with thermophilic organisms. The obtained Δ*phaC* mutant confirms the essential role of polyhydroxyalkanoate synthase in the PHA biosynthetic apparatus of *C. thermodepolymerans*. The mutant can serve as a control in studies investigating the significance of PHA synthesis metabolism in *C. thermodepolymerans* DSM 15344 and related thermophilic PHA producers. Alternatively, it can be equipped with a suitable substrate-flexible thermophilic PHA synthase, allowing for the production of superior PHA polymers with tailored monomer compositions. PHA synthases found, e.g., in thermophilic members of the genus *Aneurinibacillus*, are promising candidates (Rehakova *et al*., 2023). The construction of a triple-deletion chassis strain AI01 with improved transformation efficiency and enhanced amenability to genetic manipulations paves the way towards further tailoring of thermophilic *Caldimonas* with genetic and metabolic engineering. In particular, it is attractive to further broaden the bacterium’s substrate scope and its rather narrow PHA product portfolio or to study the relation between PHA biosynthesis and tolerance of bacterial cells to higher temperatures. Our future efforts will also focus on refining of introduced editing techniques (e.g., by increasing the frequency of homologous recombination in *C. thermodepolymerans* via employing exogenous recombinases), exploring alternative plasmid systems with thermostable origins of replication, and leveraging advanced genome-editing tools such as CRISPR/Cas technology to unlock the full biotechnological potential of *C. thermodepolymerans*.

## Acknowledgements

We thank Pablo I. Nikel and Daniel C. Volke (Novo Nordisk Foundation Center for Biosustainability, Kogens Lyngby, Denmark) for providing the pSNW2 plasmid and to Christopher W. Johnson (National Renewable Energy Laboratory, Golden, USA) for providing the pCJ012 plasmid. We also thank Esteban Martínez-García, Víctor de Lorenzo (CNB-CSIC, Madrid, Spain) and the SEVA repository (https://seva-plasmids.com/) for providing us with the SEVA and pBAMD plasmid vectors. This project was funded by the Czech Science Foundation (project registration number 22-10845S) and the Grant Agency of Masaryk University (CAREER RESTART, SYNBIOTHERM, MUNI/R/1266/2022).

## Author contributions

**Anastasiia Grybchuk-Ieremenko**: Investigation; methodology; conceptualization; writing – original draft; writing – review and editing; visualization; formal analysis. **Kristýna Lipovská**: Investigation; methodology; writing – review and editing. **Xenie Kouřilová**: Investigation; methodology; writing – review and editing. **Stanislav Obruča**: Conceptualization; writing – review and editing; supervision. **Pavel Dvořák**: Conceptualization; funding acquisition; supervision; writing – original draft; writing – review and editing. All co-authors read and approved the final version of the manuscript.

## Data Availability Statement

The data that support the findings of this study are available in the supplementary material of this article and from the corresponding author upon request.

## References

Antoine, R. and Locht, C. (1992) Isolation and molecular characterization of a novel broad-host-range plasmid from *Bordetella bronchiseptica* with sequence similarities to plasmids from Gram-positive organisms. Molecular Microbiology 6: 1785–1799.

Arai, T., Aikawa, S., Sudesh, K., Kondo, T., and Kosugi, A. (2022) Electrotransformation of thermophilic bacterium *Caldimonas manganoxidans*. J Microbiol Methods 192: 106375.

Averhoff, B. and Friedrich, A. (2003) Type IV pili-related natural transformation systems: DNA transport in mesophilic and thermophilic bacteria. Archives of Microbiology 180: 385–393.

Bertran-Llorens, S., Zhou, W., Palazzolo, M.A., Colpa, D.L., Euverink, G.-J.W., Krooneman, J., and Deuss, P.J. (2024) ALACEN: A Holistic Herbaceous Biomass Fractionation Process Attaining a Xylose-Rich Stream for Direct Microbial Conversion to Bioplastics. ACS Sustainable Chem Eng 12: 7724–7738.

Boyer, H.W. and Roulland-dussoix, D. (1969) A complementation analysis of the restriction and modification of DNA in *Escherichia coli*. J Mol Biology 41: 459–472.

Brewster, J.D. (2003) A simple micro-growth assay for enumerating bacteria. Journal of Microbiological Methods 53: 77–86.

Bujdoš, D., Popelářová, B., Volke, D.C., Nikel, P.I., Sonnenschein, N., and Dvořák, P. (2023) Engineering of *Pseudomonas putida* for accelerated co-utilization of glucose and cellobiose yields aerobic overproduction of pyruvate explained by an upgraded metabolic model. Metabolic Engineering 75: 29–46.

Carr, J.F., Danziger, M.E., Huang, A.L., Dahlberg, A.E., and Gregory, S.T. (2015) Engineering the Genome of *Thermus thermophilus* Using a Counterselectable Marker. J Bacteriol 197: 1135–1144.

Chen, G.-Q. and Jiang, X.-R. (2018) Next generation industrial biotechnology based on extremophilic bacteria. Current Opinion in Biotechnology 50: 94–100.

Choi, S.Y., Cho, I.J., Lee, Y., Kim, Y.-J., Kim, K.-J., and Lee, S.Y. (2020) Microbial Polyhydroxyalkanoates and Nonnatural Polyesters. Advanced Materials 32: 1907138.

Chung, D., Cha, M., Farkas, J., and Westpheling, J. (2013) Construction of a Stable Replicating Shuttle Vector for *Caldicellulosiruptor* Species: Use for Extending Genetic Methodologies to Other Members of This Genus. PLoS ONE 8: e62881.

Dietrich, K., Dumont, M.-J., Del Rio, L.F., and Orsat, V. (2019) Sustainable PHA production in integrated lignocellulose biorefineries. New Biotechnology 49: 161–168.

Dvořák, P., Kováč, J., and De Lorenzo, V. (2020) Biotransformation of D-xylose to D-xylonate coupled to medium-chain-length polyhydroxyalkanoate production in cellobiose-grown *Pseudomonas putida* EM42. Microbial Biotechnology 13: 1273–1283.

F. Bosma, E., Van Der Oost, J. M. De Vos, W., and Van Kranenburg, R. (2013) Sustainable Production of Bio-Based Chemicals by Extremophiles. CBIOT 2: 360–379.

Glenn, A.W., Roberto, F.F., and Ward, T.E. (1992) Transformation of *Acidiphilium* by electroporation and conjugation. Can J Microbiol 38: 387–393.

Higuchi, R., Krummel, B., and Saiki, R. (1988) A general method of *in vitro* preparation and specific mutagenesis of DNA fragments: study of protein and DNA interactions. Nucl Acids Res 16: 7351–7367.

Hon, J., Borko, S., Stourac, J., Prokop, Z., Zendulka, J., Bednar, D., et al. (2020) EnzymeMiner: automated mining of soluble enzymes with diverse structures, catalytic properties and stabilities. Nucleic Acids Research 48: W104–W109.

Hu, S., Giacopazzi, S., Modlin, R., Karplus, K., Bernick, D.L., and Ottemann, K.M. (2023) Altering under-represented DNA sequences elevates bacterial transformation efficiency. mBio 14: e02105–23.

Ishikawa, M. and Hori, K. (2024) The elimination of two restriction enzyme genes allows for electroporation-based transformation and CRISPR-Cas9-based base editing in the non-competent Gram-negative bacterium Acinetobacter sp. Tol 5. Appl Environ Microbiol 90: e00400–24.

Johnson, C.W. and Beckham, G.T. (2015) Aromatic catabolic pathway selection for optimal production of pyruvate and lactate from lignin. Metabolic Engineering 28: 240–247.

Kessler, B., Herrero, M., Timmis, K.N., and De Lorenzo, V. (1994) Genetic evidence that the XylS regulator of the *Pseudomonas* TOL meta operon controls the Pm promoter through weak DNA-protein interactions. J Bacteriol 176: 3171–3176.

Kolek, J., Sedlar, K., Provaznik, I., and Patakova, P. (2016) Dam and Dcm methylations prevent gene transfer into *Clostridium pasteurianum* NRRL B-598: development of methods for electrotransformation, conjugation, and sonoporation. Biotechnol Biofuels 9: 14.

Kong, L., Xiong, Z., Xia, Y., and Ai, L. (2021) High-efficiency transformation of *Streptococcus thermophilus* using electroporation. J Sci Food Agric 101: 6578–6585.

Kourilova, X., Novackova, I., Koller, M., and Obruca, S. (2021) Evaluation of mesophilic *Burkholderia sacchari*, thermophilic *Schlegelella thermodepolymerans* and halophilic *Halomonas halophila* for polyhydroxyalkanoates production on model media mimicking lignocellulose hydrolysates. Bioresour Technol 325: 124704.

Kourilova, X., Pernicova, I., Sedlar, K., Musilova, J., Sedlacek, P., Kalina, M., et al. (2020) Production of polyhydroxyalkanoates (PHA) by a thermophilic strain of *Schlegelella thermodepolymerans* from xylose rich substrates. Bioresour Technol 315: 123885.

Kudryavtseva, A.A., Cséfalvay, E., Gnuchikh, E.Y., Yanovskaya, D.D., Skutel, M.A., Isaev, A.B., et al. (2023) Broadness and specificity: ArdB, ArdA, and Ocr against various restriction-modification systems. Front Microbiol 14: 1133144.

Lau, W.W.Y., Shiran, Y., Bailey, R.M., Cook, E., Stuchtey, M.R., Koskella, J., et al. (2020) Evaluating scenarios toward zero plastic pollution. Science 369: 1455–1461.s

Lim, X. (2021) Microplastics are everywhere — but are they harmful? Nature 593: 22–25.

Ma, H., Zhao, Y., Huang, W., Zhang, L., Wu, F., Ye, J., and Chen, G.-Q. (2020) Rational flux-tuning of *Halomonas bluephagenesis* for co-production of bioplastic PHB and ectoine. Nat Commun 11: 3313.

Manoil, C. and Beckwith, J. (1985) TnphoA: a transposon probe for protein export signals. Proc Natl Acad Sci USA 82: 8129–8133.

Martínez-García, E., Aparicio, T., De Lorenzo, V., and Nikel, P.I. (2014) New Transposon Tools Tailored for Metabolic Engineering of Gram-Negative Microbial Cell Factories. Front Bioeng Biotechnol 2:.

Martínez-García, E., Fraile, S., Algar, E., Aparicio, T., Velázquez, E., Calles, B., et al. (2023) SEVA 4.0: an update of the Standard European Vector Architecture database for advanced analysis and programming of bacterial phenotypes. Nucleic Acids Research 51: D1558–D1567.

Meereboer, K.W., Misra, M., and Mohanty, A.K. (2020) Review of recent advances in the biodegradability of polyhydroxyalkanoate (PHA) bioplastics and their composites. Green Chem 22: 5519–5558.

Musilova, J., Kourilova, X., Bezdicek, M., Lengerova, M., Obruca, S., Skutkova, H., and Sedlar, K. (2021) First Complete Genome of the Thermophilic Polyhydroxyalkanoates-Producing Bacterium *Schlegelella thermodepolymerans* DSM 15344. Genome Biology and Evolution 13: evab007.

Musilova, J., Kourilova, X., Hermankova, K., Bezdicek, M., Ieremenko, A., Dvorak, P., et al. (2023) Genomic and phenotypic comparison of polyhydroxyalkanoates producing strains of genus *Caldimonas/Schlegelella*. Computational and Structural Biotechnology Journal 21: 5372–5381.

Obruča, S., Dvořák, P., Sedláček, P., Koller, M., Sedlář, K., Pernicová, I., and Šafránek, D. (2022) Polyhydroxyalkanoates synthesis by halophiles and thermophiles: towards sustainable production of microbial bioplastics. Biotechnol Adv 58: 107906.

Obruca, S., Snajdar, O., Svoboda, Z., and Marova, I. (2013) Application of random mutagenesis to enhance the production of polyhydroxyalkanoates by *Cupriavidus necator* H16 on waste frying oil. World J Microbiol Biotechnol 29: 2417–2428.

Olson, D.G. and Lynd, L.R. (2012) Computational design and characterization of a temperature-sensitive plasmid replicon for gram positive thermophiles. J Biol Eng 6: 5.

Oren, A. (2011) Thermodynamic limits to microbial life at high salt concentrations. Environmental Microbiology 13: 1908–1923.

Pal, U., Bachmann, D., Pelzer, C., Christiansen, J., Blank, L.M., and Tiso, T. (2024) A genetic toolbox to empower *Paracoccus pantotrophus* DSM 2944 as a metabolically versatile SynBio chassis. Microb Cell Fact 23: 53.

Palmeiro-Sánchez, T., O’Flaherty, V., and Lens, P.N.L. (2022) Polyhydroxyalkanoate bio-production and its rise as biomaterial of the future. Journal of Biotechnology 348: 10–25.

Rehakova, V., Pernicova, I., Kourilova, X., Sedlacek, P., Musilova, J., Sedlar, K., et al. (2023) Biosynthesis of versatile PHA copolymers by thermophilic members of the genus *Aneurinibacillus*. International Journal of Biological Macromolecules 225: 1588–1598.

Ren, J., Lee, H.-M., Shen, J., and Na, D. (2022) Advanced biotechnology using methyltransferase and its applications in bacteria: a mini review. Biotechnol Lett 44: 33–44.

Ried, J.L. and Collmer, A. (1987) An nptI-sacB-sacR cartridge for constructing directed, unmarked mutations in Gram-negative bacteria by marker exchange-eviction mutagenesis. Gene 57: 239–246.

Samperio, S., Guzmán-Herrador, D.L., May-Cuz, R., Martín, M.C., Álvarez, M.A., and Llosa, M. (2021) Conjugative DNA Transfer From *E. coli* to Transformation-Resistant Lactobacilli. Front Microbiol 12: 606629.

Sánchez-Romero, M.A., Cota, I., and Casadesús, J. (2015) DNA methylation in bacteria: from the methyl group to the methylome. Current Opinion in Microbiology 25: 9–16.

Silva-Rocha, R., Martínez-García, E., Calles, B., Chavarría, M., Arce-Rodríguez, A., de Las Heras, A., et al. (2013) The Standard European Vector Architecture (SEVA): a coherent platform for the analysis and deployment of complex prokaryotic phenotypes. Nucleic Acids Res 41: D666–675.

Stegmann, P., Daioglou, V., Londo, M., van Vuuren, D.P., and Junginger, M. (2022) Plastic futures and their CO2 emissions. Nature 612: 272–276.

Suzuki, H. (2012) Host-Mimicking Strategies in DNA Methylation for Improved Bacterial Transformation. In Methylation - From DNA, RNA and Histones to Diseases and Treatment. Dricu, A. (ed). InTech.

Taylor, R.G., Walker, D.C., and Mclnnes, R.R. (1993) *E.coli* host strains significantly affect the quality of small scale plasmid DNA preparations used for sequencing. Nucl Acids Res 21: 1677–1678.

Turner, P., Mamo, G., and Karlsson, E.N. (2007) Potential and utilization of thermophiles and thermostable enzymes in biorefining. Microb Cell Fact 6: 9.

Volke, D.C., Friis, L., Wirth, N.T., Turlin, J., and Nikel, P.I. (2020) Synthetic control of plasmid replication enables target- and self-curing of vectors and expedites genome engineering of *Pseudomonas putida*. Metabolic Engineering Communications 10: e00126.

Yanisch-Perron, C., Vieira, J., and Messing, J. (1985) Improved M13 phage cloning vectors and host strains: nucleotide sequences of the M13mpl8 and pUC19 vectors. Gene 33: 103–119.

Ye, J., Hu, D., Che, X., Jiang, X., Li, T., Chen, J., et al. (2018) Engineering of *Halomonas bluephagenesis* for low cost production of poly(3-hydroxybutyrate-co-4-hydroxybutyrate) from glucose. Metab Eng 47: 143–152.

Ye, J.-W., Lin, Y.-N., Yi, X.-Q., Yu, Z.-X., Liu, X., and Chen, G.-Q. (2022) Synthetic biology of extremophiles: a new wave of biomanufacturing. Trends in Biotechnology.

Yu, J. and Chen, L.X.L. (2008) The Greenhouse Gas Emissions and Fossil Energy Requirement of Bioplastics from Cradle to Gate of a Biomass Refinery. Environ Sci Technol 42: 6961–6966.

Zhang, X., Liang, Y., Yang, Haibo, Yang, Hui, Chen, S., Huang, F., et al. (2021) A novel fusion levansucrase improves thermostability of polymerization and production of high molecular weight levan. LWT 150: 111951.

Zheng, Y., Chen, J.-C., Ma, Y.-M., and Chen, G.-Q. (2020) Engineering biosynthesis of polyhydroxyalkanoates (PHA) for diversity and cost reduction. Metabolic Engineering 58: 82–93.

Zhou, Jia, Li, X., Xia, J., Wen, Y., Zhou, Jie, Yu, Z., and Tian, B. (2018) The role of temperature and bivalent ions in preparing competent *Escherichia coli*. 3 Biotech 8: 222.

Zhou, W., Colpa, D.I., Permentier, H., Offringa, R.A., Rohrbach, L., Euverink, G.-J.W., and Krooneman, J. (2023) Insight into polyhydroxyalkanoate (PHA) production from xylose and extracellular PHA degradation by a thermophilic *Schlegelella thermodepolymerans*. Resources, Conservation and Recycling 194: 107006.

