## Supplementary Information for "An initial genome editing toolset for *Caldimonas* t*hermodepolymerans*, the first model of thermophilic polyhydroxyalkanoates producer"

by

Anastasiia Grybchuk-Ieremenko[1], Kristýna Lipovská[1], Xenie Kouřilová[2], Stanislav Obruča[2], Pavel Dvořák[1*]

[1] Dpt. of Experimental Biology (Section of Microbiology), Faculty of Science, Masaryk University, Kamenice 5/E25, 62500 Brno, Czech Republic

[2] Institute of Food Science and Biotechnology, Faculty of Chemistry, Brno University of Technology, Purkyňova 464/118, Královo Pole, 61200, Brno, Czech Republic

* Corresponding author:

Pavel Dvořák, Assoc. Prof.

Department of Experimental Biology (Section of Microbiology), Faculty of Science

Masaryk University, Kamenice 735/5, Brno 62500, Czech Republic

### Content


***Supplementary tables***

**Table S1.** Oligonucleotide primers used in this study.

***Supplementary figures***

**Figure S1.** Integration of pSNW2_14025 plasmid into *Caldimonas* chromosome using 14025_HomS1 for, pSNW2 flank rev (A) and pSNW2 flank for, 14025_HomS2 rev (B) pairs of primers.

***Nucleotide sequences of genes and plasmid constructs used in this study***

**Gene of the superfolder GFP**

**Thermostable codon-optimized *sacB* gene**

**Plasmid pBAMD_*gfp***

**Plasmid pAI012_HA2**

**Plasmid pSNW2_HA2**

**Plasmid pSNW2_8585**

**Plasmid pSNW2_14855**

**Plasmid pSNW2_14025**

**Plasmid pAI_ThSacB**

### Supplementary tables

**Table S1.** Oligonucleotide primers used in this study (restriction sites are underlined, and overlapped sequences are in lowercase).

| Primer | Sequence | Usage |
|---|---|---|
| HAN1 | AGAACAGCTCGACGATCACCAGCG | Amplification of *phaC* upstream homologous region, forward primer |
| HR1rev | atgccggaaggcgccttcaCATCCTCTCCTCGTGTCAGTGAACG | Amplification of *phaC* upstream homologous region, reverse primer |
| HR2for | TGAAGGCGCCTTCCGGCAT | Amplification of *phaC* downstream homologous region, forward primer |
| HAN2 | CTGGACAACGTCTGGCCCGAG | Amplification of *phaC* downstream homologous region, reverse primer |
| HA1for | AGAACAGCTCGACGATCACCAGCG | Amplification of overlapped homologous sequence, forward primer (flanking *phaC*) |
| HA2_XbaIrev | ATAT<u>TCTAGA</u>TTCTTCGCATCGGCAGGGCTG | Amplification of overlapped homologous sequence, reverse primer (flanking *phaC*) |
| pSNW2 flank for | CCAGGGTTTTCCCAGTCACGAC | Check of the assembly of the pSNW2_HA2, pAI_ThSacB vector, and target plasmid integration into the chromosome |
| pSNW2 flank rev | CGGATTACCCTGTTATCCCTAGAAG | Check of the assembly of the vector and target plasmid integration into the chromosome |
| Gen1for | AGAACAGCTCGACGATCACCAG | Check of the pSNW2_HA2, pAI012_HA2, pAI_ThSacB plasmid integration into the chromosome (*phaC*) |
| Gen2 rev | CTGGACAACGTCTGGCCC | Check of the plasmid integration into the chromosome (*phaC*) |
| 08585_HomS1 for | GTATAGGGGTTGAAGACGCCATTG | Amplification of IS481_08585 upstream homologous region, forward primer |
| 08585_HomS1 rev | agagatctttgccgcatgTGACCCATGCCCAACGTCTC | Amplification of IS481_08585 upstream homologous region, reverse primer |
| 08585_HomS2 for | CATGCGGCAAAGATCTCTATTGTGAG | Amplification of IS481_08585 downstream homologous region, forward primer |

**Table S1.** Continuing

| | | |
|---|---|---|
| EcoRI _08585_HomS for | TAGAATTCGATGGTGTGCAGGTTGCACTC | Amplification of overlapped homologous sequence, forward primer (flanking IS481_08585) |
| XbaI _08585_HomS rev | TATTCTAGACTTCGGGATCTCGATCAACCAG | Amplification of overlapped homologous sequence, reverse primer (flanking IS481_08585) |
| 14855_HomS1 for | GATAGAGAGATGCTGTGTATAGATACGCTAGTTC | Amplification of IS481_14855 upstream homologous region, forward primer |
| 14855_HomS1 rev | tgcccaggtagctgcatgTAGCCTAGCTGGTTTCGACGT | Amplification of IS481_14855 upstream homologous region, reverse primer |
| 14855_HomS2 for | CATGCAGCTACCTGGGCAC | Amplification of IS481_14855 downstream homologous region, forward primer |
| 14855_HomS2 rev | GTCTGGGACGTACTTCAAAGCC | Amplification of IS481_14855 downstream homologous region, reverse primer |
| KpnI _14855_HomS for | TTGGTACCCATGTCTAGCTGCTCGCTGAAG | Amplification of overlapped homologous sequence, forward primer (flanking IS481_14855) |
| XbaI _14855_HomS rev | TTTCTAGATGCTGCGTCATCTCACGTTC | Amplification of overlapped homologous sequence, reverse primer (flanking IS481_14855) |
| 14025_HomS1 for | CTTGCATCGTTTCGTGTAGG | Amplification of IS481_14025 upstream homologous region, forward primer |
| 14025_HomS1 rev | ggcggcttctttatttcaCATGATTCGAGGTCCTCG | Amplification of IS481_14025 upstream homologous region, reverse primer |
| 14025_HomS2 for | TGAAATAAAGAAGCCGCCCGAGGGCG | Amplification of IS481_14025 downstream homologous region, forward primer |
| 14025_HomS2 rev | CAACAGCCTCTCGAAGCAGATC | Amplification of IS481_14025 downstream homologous region, reverse primer |
| EcoRI _14025_HomS for | TAGAATTCGCAATCTGATCTTCGAGCACGTTG | Amplification of overlapped homologous sequence, forward primer (flanking IS481_14025) |
| XbaI _14025_HomS rev | TATCTAGACTCAAGGCCAAGGGCGAG | Amplification of overlapped continuous homologous sequence, reverse primer (flanking IS481_14025) |

**Table S1.** Continuing

| | | |
|---|---|---|
| 8585_Gen1 for | CTCGTAATACCAGATCTCCTTCGTCG | Check of the pSNW2_8585 plasmid integration into the chromosome (IS481_08585) |
| 8585_Gen2 rev | CCTCTCTTTACAAACCAACCAGAAACTG | Check of the pSNW2_8585 plasmid integration into the chromosome (IS481_08585) |
| 14855_Gen1 for | GTTCATTCATAGAGTCTCCGCAGG | Check of the pSNW2_14855 plasmid integration into the chromosome (IS481_14855) |
| 14855_Gen2 rev | CGTCTCACACCACCAGATGG | Check of the pSNW2_14855 plasmid integration into the chromosome (IS481_14855) |
| 14025_Gen1 for | GGCAGCACCGCAGGAAAAC | Check of the pSNW2_14025 plasmid integration into the chromosome (IS481_14025) |
| 14025_Gen2 rev | GAGGCGATCCAGTCCGAGC | Check of the pSNW2_14025 plasmid integration into the chromosome (IS481_14025) |
| HAN1 for | AGAACAGCTCGACGATCACCAGCG | Amplification of *phaC* upstream homologous region and amplification of overlapped homologous sequence, forward primer |
| HR1 Rev | ATGCCGGAAGGCGCCTTCACATCCTCTCCTCGTGTCAG TGAACG | Amplification of *phaC* upstream homologous region, reverse primer |
| HR2 For | TGAAGGCGCCTTCCGGCAT | Amplification of *phaC* downstream homologous region, forward primer |
| HAN2 | CTGGACAACGTCTGGCCCGAG | Amplification of *phaC* downstream homologous region and amplification of overlapped homologous sequence, reverse primer |
| HA1 For | ATAAGAATTCCATAGACCAGCGGTTCGCCGTC | Amplification of overlapped continuous homologous sequence, forward primer (flanking *phaC*) |
| HA2_HindIII rev | ATATAAGCTTTTCTTCGCATCGGCAGGGCTG | Amplification of overlapped continuous homologous sequence, reverse primer (flanking *phaC*) |

**Table S1.** Continuing

| | | |
|---|---|---|
| pCJ012 flank for | CTCGTATGTTGTGTGGAATTGTG | Check of the assembly of the pAI012_HA2 vector and plasmid integration into the chromosome (*phaC*) |
| pCJ012 flank rev | CGTTGTAAAACGACGGCCAG | Check of the assembly of the pAI012_HA2 vector and plasmid integration into the chromosome (*phaC)* |
| pCJ012 gen flank for | CTTACGATGGGCTGAACCTG | Check of the assembly of the pAI012_HA2 vector and target plasmid integration into the chromosome (*phaC)* |
| pCJ012 gen flank rev | CAGCCCATTTCCTTGTGCAC | Check of the assembly of the pAI012_HA2 vector and target plasmid integration into the chromosome (*phaC)* |
| PI for new | gaactgctgatcttcagatcGCCTTTTGCTCACATGTTCTTTCCT | Amplification of pCJ012 plasmid backbone without PMB1 origin for in vivo cloning, forward primer |
| PI rev new | TTTGATCTTTTCTACGGGGTCTGACG | Amplification of pCJ012 plasmid backbone without PMB1 origin for in vivo cloning, reverse primer |
| R6K for new | ccccgtagaaaagatcaaaCCATGTCAGCCGTTAAGTGTTCC | Amplification of R6K origin from pSNW2 plasmid for in vivo cloning, forward primer |
| R6K rev | GATCTGAAGATCAGCAGTTCAACCTGT | Amplification of R6K origin from pSNW2 plasmid for in vivo cloning, reverse primer |
| Ori check for | CTAGGTGAAGATCCTTTTTGATAATCTCATG | Check for the assembly of the R6K_pCJ012 vector, forward primer |
| Ori check rev | AAAGGCGGTAATACGGTTATCCAC | Check for the assembly of the R6K_pCJ012 vector, reverse primer |
| ThSacB for | ccaggtataattgcacgaATTACATATTGAAAAAGGGAGG | Amplification of *sacB* from commercially ordered plasmid ThSacB_pUC_Amp (Azenta), forward primer |
| ThSacB rev | TCACTTGTTCACGGTCAG | Amplification of *sacB* from commercially ordered plasmid ThSacB_pUC_Amp (Azenta), reverse primer |
| pSNW2_sac for | ctgaccgtgaacaagtgaCTTGGACTCCTGTTGATAGATC | Amplification of pSNW2 backbone from pSNW2_HA2 vector, forward primer |
| pSNW2_sac rev | TCGTGCAATTATACCTGGC | Amplification of pSNW2 from pSNW2_HA2 vector backbone, reverse primer |

**Table S1.** Continuing

| | | |
|---|---|---|
| SacB pl ch for | ACCGCCCAGTCTAGCTATC | Check for the assembly of the pAI_ThSacB vector, reverse primer |
| SacB pl ch rev | GCGTTCTGAACAAATCCAGATGG | Check for the assembly of the pAI_ThSacB vector, reverse primer |

### Supplementary figures

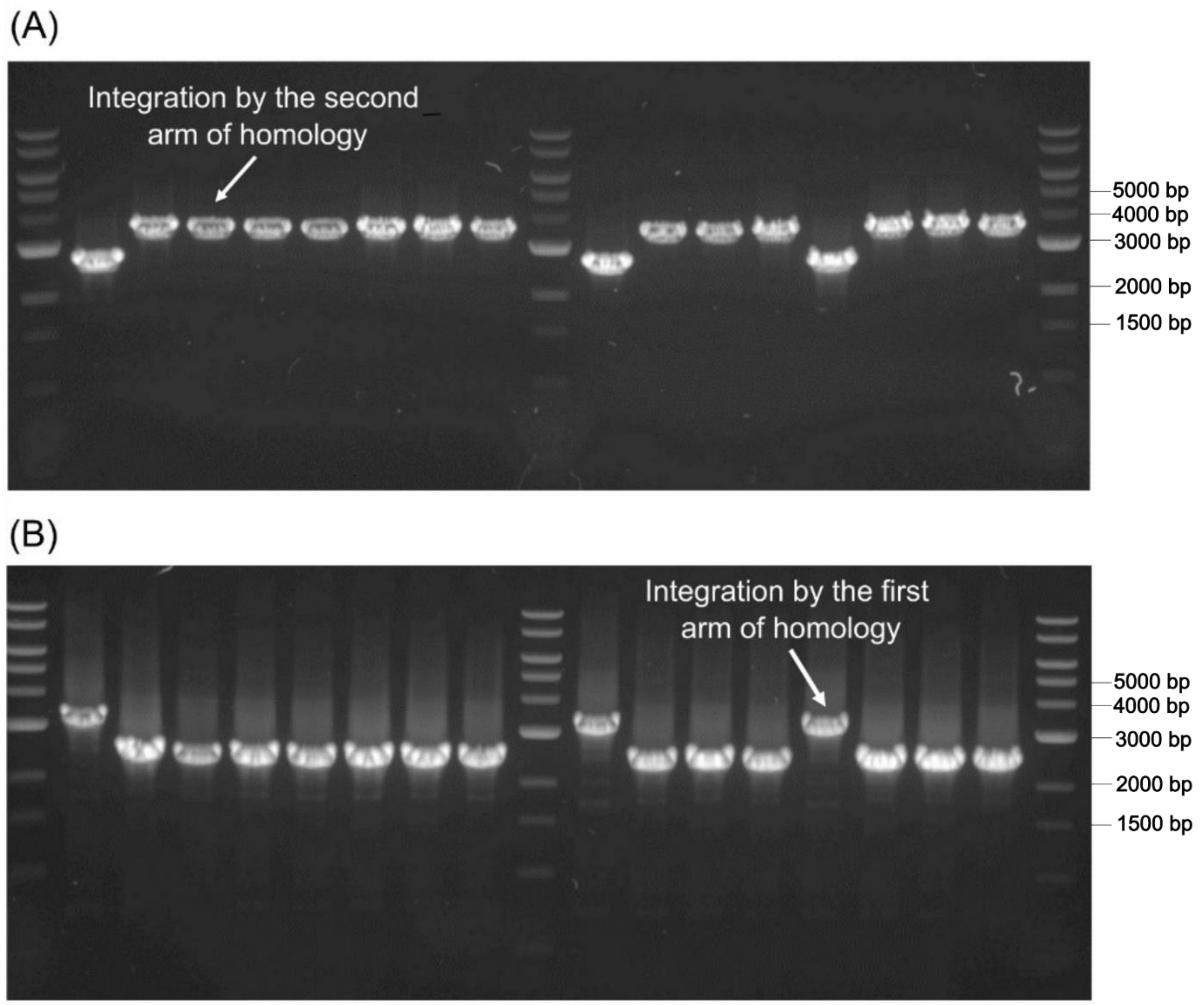


**Figure S1.** PCR products confirming the integration of pSNW2_14025 plasmid into *Caldimonas* chromosome. Primers 14025_HomS1 for and pSNW2 flank rev were used in (**A**) and pSNW2 flank for and 14025_HomS2 rev in (**B**). The expected size of the co-integrate is ~3200 bp.

### Nucleotide sequences of genes and plasmids used in this study

**Gene of the superfolder GFP** with synthetic RBS (underlined, 783 bp).

<u>TCAATTCTAGCAAGAGGAATATACAT</u>ATGGCTAGCCGTAAAGGTGAAGAACTGTTCACCGGT
GTTGTTCCGATCCTGGTTGAACTGGATGGTGATGTTAACGGCCACAAATTCTCTGTTCGTGG
TGAAGGTGAAGGTGATGCAACCAACGGTAAACTGACCCTGAAATTCATCTGCACTACCGGTA
AACTGCCGGTTCCATGGCCGACTCTGGTGACTACCCTGACCTATGGTGTTCAGTGTTTTTCT
CGTTACCCGGATCACATGAAGCAGCATGATTTCTTCAAATCTGCAATGCCGGAAGGTTATGT
ACAGGAGCGCACCATTTCTTTCAAAGACGATGGCACCTACAAAACCCGTGCAGAGGTTAAAT
TTGAAGGTGATACTCTGGTGAACCGTATTGAACTGAAAGGCATTGATTTCAAAGAGGACGGC
AACATCCTGGGCCACAAACTGGAATATAACTTCAACTCCCATAACGTTTACATCACCGCAGA
CAAACAGAAGAACGGTATCAAAGCTAACTTCAAAATTCGCCATAACGTTGAAGACGGTAGCG
TACAGCTGGCGGACCACTACCAGCAGAACACTCCGATCGGTGATGGTCCGGTTCTGCTGCCG
GATAACCACTACCTGTCCACCCAGTCTAAACTGTCCAAAGACCCGAACGAAAAGCGCGACCA
CATGGTGCTGCTGGAGTTCGTTACTGCAGCAGGTATCACGCACGGCATGGATGAACTCTACA
AATAA

**Thermostable codon-optimized *sacB*** gene with native RBS (underlined) from *B. licheniformis* (1,479 bp).

<u>ATTACATATTGAAAAAGGGAGGAAATATTG</u>ATGAACATCAAGAACATCGCCAAGAAGGCCAG
CGCCCTGACCGTGGCCGCCGCCCTGCTGGCCGGCGGCGCCCCGCAGACCTTCGCCAAGGAGA
CCCAGGACTACAAGAAGAGCTACGGCTTCAGCCACATCACCCGCCACGACATGCTGAAGATC
CCGGAGCAGCAGAAGAGCGAGCAGTTCAAGGTGCCGCAGTTCGACCCGAAGACCATCAAGAA
CATCCCGAGCGCCAAGGGCTACAACAAGAACGGCGAGCTGATCGACCTGGACGTGTGGGACA
GCTGGCCGCTGCAGAACGCCGACGGCACCGTGGCCACCTACCACGGCTACAACCTGGTGTTC
GCCCTGGCCGGCGACCCGAAGGACGTGGACGACACCAGCATCTACCTGTTCTACCAGAAGAA
GGGCGAGACCAGCATCGACAGCTGGAAGAACGCCGGCCGCGTGTTCAAGGACAGCGACAAGT
TCGTGCCGGACGACCCGTACCTGAAGCACCAGACCCAGGAGTGGAGCGGCAGCGCCACCCTG
ACCAAGGACGGCAAGGTGCGCCTGTTCTACACCGCCTTCAGCGGCACCCAGTACGGCAAGCA
GACCCTGACCACCGCCCAGGTGAACTTCAGCCAGCCGGACAGCGACACCCTGAAGATCGACG
GCGTGGAGGACCACAAGAGCGTGTTCGACGGCGCCGACGGCACCGTGTACCAGAACGTGCAG
CAGTTCATCGACGAGGGCAACTACAGCAGCGGCGACAACCACACCATGCGCGACCCGCACTA

CGTGGAGGACCGCGGCCACAAGTACCTGGTGTTCGAGGACAACACCGGCACCAAGACCGGCT
ACCAGGGCGAGGACAGCCTGTTCAACCGCGCCTACTACGGCGGCAGCAAGAAGTTCTTCAAG
GAGGAGAGCAGCAAGCTGCTGCAGGGCGCCAACAAGAAGAACGCCAGCCTGGCCAACGGCGC
CCTGGGCATCATCGAGCTGAACAACGACTACACCCTGAAGAAGGTGATGAAGCCGCTGATCG
CCAGCAACACCGTGACCGACGAGATCGAGCGCGCCAACCTGTTCAAGATGAACGGCAAGTGG
TACCTGTTCACCGACAGCCGCGGCAGCAAGATGACCATCGACGGCATCGGCAGCAAGGACAT
CTACATGCTGGGCTACGTGAGCGGCAGCCTGACCGGCCCGTTCAAGCCGCTGAACAAGAGCG
GCCTGGTGCTGCACATGGACCAGGACTACAACGACATCACCTTCACCTACAGCCACTTCGCC
GTGCCGCAGAAGAAGGGCGACGAGGTGGTGATCACCAGCTACATCACCAACCGCGGCATCAG
CAACGAGCACCACGCCACCTTCGCCCCGAGCTTCCTGCTGAAGATCAAGGGCAGCAAGACCA
GCGTGGTGAAGAACAGCATCCTGGAGCAGGGCCAGCTGACCGTGAACAAGTGA

**Plasmid pBAMD_*gfp* -** Tn5-based mini-transposon system for random integration of a gene(s) of interest (the gene of superfolder GFP in this case) into the genome (5,262 bp).

TTTAATTAACTGTCTCTTATACACATCTTTGTGTCTCAGGCCGCCTAGGCCGCGGCCGCGCG
AATTCTCAATTCTAGCAAGAGGAATATACATATGGCTAGCCGTAAAGGTGAAGAACTGTTCA
CCGGTGTTGTTCCGATCCTGGTTGAACTGGATGGTGATGTTAACGGCCACAAATTCTCTGTT
CGTGGTGAAGGTGAAGGTGATGCAACCAACGGTAAACTGACCCTGAAATTCATCTGCACTAC
CGGTAAACTGCCGGTTCCATGGCCGACTCTGGTGACTACCCTGACCTATGGTGTTCAGTGTT
TTTCTCGTTACCCGGATCACATGAAGCAGCATGATTTCTTCAAATCTGCAATGCCGGAAGGT
TATGTACAGGAGCGCACCATTTCTTTCAAAGACGATGGCACCTACAAAACCCGTGCAGAGGT
TAAATTTGAAGGTGATACTCTGGTGAACCGTATTGAACTGAAAGGCATTGATTTCAAAGAGG
ACGGCAACATCCTGGGCCACAAACTGGAATATAACTTCAACTCCCATAACGTTTACATCACC
GCAGACAAACAGAAGAACGGTATCAAAGCTAACTTCAAAATTCGCCATAACGTTGAAGACGG
TAGCGTACAGCTGGCGGACCACTACCAGCAGAACACTCCGATCGGTGATGGTCCGGTTCTGC
TGCCGGATAACCACTACCTGTCCACCCAGTCTAAACTGTCCAAAGACCCGAACGAAAAGCGC
GACCACATGGTGCTGCTGGAGTTCGTTACTGCAGCAGGTATCACGCACGGCATGGATGAACT
CTACAAATAACTGCAGGCATGCAAGCTTGCGGCCGCCAAAGCCCGCCGAAAGGCGGGCTTTT
CTGTATTTAAATTTGACATAAGCCTGTTCGGTTCGTAAACTGTAATGCAAGTAGCGTATGCG
CTCACGCAACTGGTCCAGAACCTTGACCGAACGCAGCGGTGGTAACGGCGCAGTGGCGGTTT
TCATGGCTTGTTATGACTGTTTTTTTGTACAGCCTATGCCTCGGGCATCCAAGCAGCAAGCG
CGTTACGCCGTGGGTCGATGTTTGATGTTATGGAGCAGCAACGATGTTACGCAGCAGCAACG
ATGTTACGCAGCAGGGCAGTCGCCCTAAAACAAAGTTAGGTGGCTCAAGTATGGGCATCATT

CGCACATGTAGGCTCGGCCCTGACCAAGTCAAATCCATGCGGGCTGCTCTTGATCTTTTCGG
TCGTGAGTTCGGAGACGTAGCCACCTACTCCCAACATCAGCCGGACTCCGATTACCTCGGGA
ACTTGCTCCGTAGTAAGACATTCATCGCGCTTGCTGCCTTCGACCAAGAAGCGGTTGTTGGC
GCTCTCGCGGCTTACGTTCTGCCCAAGTTTGAGCAGCCGCGTAGTGAGATCTATATCTATGA
TCTCGCAGTCTCCGGAGAGCACCGGAGGCAGGGCATTGCCACCGCGCTCATCAATCTCCTCA
AGCATGAGGCCAACGCGCTTGGTGCTTATGTGATCTACGTGCAAGCAGATTACGGTGACGAT
CCCGCAGTGGCTCTCTATACAAAGTTGGGCATACGGGAAGAAGTGATGCACTTTGATATCGA
CCCAAGTACCGCCACCTAACAATTCGTTCAAGCCGAGATCGGCTTCCCGGCCGCGGAGTTGT
TCGGTAAATTGGACAATTGTCTAATTAATTGCGGACCCTAGAGGTCCCCTTTTTTATTTTAA
AAATTTTTTCACAAAACGGTTTACAAGCATAAAATCTCTGAAGATGTGTATAAGAGACAGAC
TAGTCTTGGACTCCTGTTGATAGATCCAGTAATGACCTCAGAACTCCATCTGGATTTGTTCA
GAACGCTCGGTTGCCGCCGGGCGTTTTTTATTGGTGAGAATCCAGGGGTCCCCTGGTTTAAA
CTACACAAGTAGCGTCCTGAACGGAACCTTTCCCGTTTTCCAGAATCTGATGTTCCATGTGA
CCTCCTAACATGGTAACGTTCATGATTACCAGTGCACTGCATCGTGCGGCGGATTGGGCGAA
AAGCGTGTTTTCTAGTGCTGCGCTGGGTGATCCGCGTCGTACCGCGCGTCTGGTGAATGTTG
CGGCGCAACTGGCCAAATATAGCGGCAAAAGCATTACCATTAGCAGCGAAGGCAGCAAAGCC
ATGCAGGAAGGCGCGTATCGTTTTATTCGTAATCCGAACGTGAGCGCGGAAGCGATTCGTAA
AGCGGGTGCCATGCAGACCGTGAAACTGGCCCAGGAATTTCCGGAACTGCTGGCAATTGAAG
ATACCACCTCTCTGAGCTATCGTCATCAGGTGGCGGAAGAACTGGGCAAACTGGGTAGCATT
CAGGATAAAAGCCGTGGTTGGTGGGTGCATAGCGTGCTGCTGCTGGAAGCGACCACCTTTCG
TACCGTGGGCCTGCTGCATCAAGAATGGTGGATGCGTCCGGATGATCCGGCGGATGCGGATG
AAAAAGAAAGCGGCAAATGGCTGGCCGCTGCTGCAACTTCGCGTCTGAGAATGGGCAGCATG
ATGAGCAACGTGATTGCGGTGTGCGATCGTGAAGCGGATATTCATGCGTATCTGCAAGATAA
ACTGGCCCATAACGAACGTTTTGTGGTGCGTAGCAAACATCCGCGTAAAGATGTGGAAAGCG
GCCTGTATCTGTATGATCACCTGAAAAACCAGCCGGAACTGGGCGGCTATCAGATTAGCATT
CCGCAGAAAGGCGTGGTGGATAAACGTGGCAAACGTAAAAACCGTCCGGCGCGTAAAGCGAG
CCTGAGCCTGCGTAGCGGCCGTATTACCCTGAAACAGGGCAACATTACCCTGAACGCGGTGC
TGGCCGAAGAAATTAATCCGCCGAAAGGCGAAACCCCGCTGAAATGGCTGCTGCTGACCAGC
GAGCCGGTGGAAAGTCTGGCCCAAGCGCTGCGTGTGATTGATATTTATACCCATCGTTGGCG
CATTGAAGAATTTCACAAAGCGTGGAAAACGGGTGCGGGTGCGGAACGTCAGCGTATGGAAG
AACCGGATAACCTGGAACGTATGGTGAGCATTCTGAGCTTTGTGGCGGTGCGTCTGCTGCAA
CTGCGTGAATCTTTTACTCCGCCGCAAGCACTGCGTGCGCAGGGCCTGCTGAAAGAAGCGGA
ACACGTTGAAAGCCAGAGCGCGGAAACCGTGCTGACCCCGGATGAATGCCAACTGCTGGGCT
ATCTGGATAAAGGCAAACGCAAACGCAAAGAAAAAGCGGGCAGCCTGCAATGGGCGTATATG

GCGATTGCGCGTCTGGGCGGCTTTATGGATAGCAAACGTACCGGCATTGCGAGCTGGGGTGC
GCTGTGGGAAGGTTGGGAAGCGCTGCAAAGCAAACTGGATGGCTTTCTGGCCGCGAAAGACC
TGATGGCGCAGGGCATTAAAATCTAATCGGCGATCGCTAGTAGCCCGCCTAATGAGCGGGCT
TTTTTTTAATTCCCCTATTTGTTTATTTTTCTAAATACATTCAAATATGTATCCGCTCATGA
GACAATAACCCTGATAAATGCTTCAATAATATTGAAAAAGGAAGAGTATGAGCATTCAGCAT
TTTCGTGTGGCGCTGATTCCGTTTTTTGCGGCGTTTTGCCTGCCGGTGTTTGCGCATCCGGA
AACCCTGGTGAAAGTGAAAGATGCGGAAGATCAACTGGGTGCGCGCGTGGGCTATATTGAAC
TGGATCTGAACAGCGGCAAAATTCTGGAATCTTTTCGTCCGGAAGAACGTTTTCCGATGATG
AGCACCTTTAAAGTGCTGCTGTGCGGTGCGGTTCTGAGCCGTGTGGATGCGGGCCAGGAACA
ACTGGGCCGTCGTATTCATTATAGCCAGAACGATCTGGTGGAATATAGCCCGGTGACCGAAA
AACATCTGACCGATGGCATGACCGTGCGTGAACTGTGCAGCGCGGCGATTACCATGAGCGAT
AACACCGCGGCGAACCTGCTGCTGACGACCATTGGCGGTCCGAAAGAACTGACCGCGTTTCT
GCATAACATGGGCGATCATGTGACCCGTCTGGATCGTTGGGAACCGGAACTGAACGAAGCGA
TTCCGAACGATGAACGTGATACCACCATGCCGGCAGCAATGGCGACCACCCTGCGTAAACTG
CTGACGGGTGAGCTGCTGACCCTGGCAAGCCGCCAGCAACTGATTGATTGGATGGAAGCGGA
TAAAGTGGCGGGTCCGCTGCTGCGTAGCGCGCTGCCGGCTGGCTGGTTTATTGCGGATAAAA
GCGGTGCGGGCGAACGTGGCAGCCGTGGCATTATTGCGGCGCTGGGCCCGGATGGTAAACCG
AGCCGTATTGTGGTGATTTATACCACCGGCAGCCAGGCGACGATGGATGAACGTAACCGTCA
GATTGCGGAAATTGGCGCGAGCCTGATTAAACATTGGTAAACCGATACAATTAAAGGCTCCT
TTTGGAGCCTTTTTTTTTGGACACGCGTTTGTCCTTTTCCGCTGCATAACCCTGCTTCGGGG
TCATTATAGCGATTTTTTCGGTATATCCATCCTTTTTCGCACGATATACAGGATTTTGCCAA
AGGGTTCGTGTAGACTTTCCTTGGTGTATCCAACGGCGTCAGCCGGGCAGGATAGGTGAAGT
AGGCCCACCCGCGAGCGGGTGTTCCTTCTTCACTGTCCCTTATTCGCACCTGGCGGTGCTCA
ACGGGAATCCTGCTCTGCGAGGCTGGCCGTAGGCCGGCCGATCTGAAGATCAGCAGTTCAAC
CTGTTGATAGTACGTACTAAGCTCTCATGTTTCACGTACTAAGCTCTCATGTTTAACGTACT
AAGCTCTCATGTTTAACGAACTAAACCCTCATGGCTAACGTACTAAGCTCTCATGGCTAACG
TACTAAGCTCTCATGTTTCACGTACTAAGCTCTCATGTTTGAACAATAAAATTAATATAAAT
CAGCAACTTAAATAGCCTCTAAGGTTTTAAGTTTTATAAGAAAAAAAAGAATATATAAGGCT
TTTAAAGCCTTTAAGGTTTAACGGTTGTGGACAACAAGCCAGGGATGTAACGCACTGAGAAG
CCCTTAGAGCCTCTCAAAGCAATTTTGAGTGACACAGGAACACTTAACGGCTGACATGGGGC
GCGCCCAGCTGTCTAGGGCGGCGGATTTGTCCTACTCAGGAGAGCGTTCACCGACAAACAAC
AGATAAAACGAAAGGCCCAGTCTTTCGACTGAGCCTTTCGTTTTATTTGATGCC

**Plasmid pAI012_HA2 -** plasmid for deletion of *phaC* gene via homologous recombination and SacB-mediated (mesophilic origin) sucrose counter-selection (9,976 bp).

TGCCGCAAGCACTCAGGGCGCAAGGGCTGCTAAAGGAAGCGGAACACGTAGAAAGCCAGTCC
GCAGAAACGGTGCTGACCCCGGATGAATGTCAGCTACTGGGCTATCTGGACAAGGGAAAACG
CAAGCGCAAAGAGAAAGCAGGTAGCTTGCAGTGGGCTTACATGGCGATAGCTAGACTGGGCG
GTTTTATGGACAGCAAGCGAACCGGAATTGCCAGCTGGGGCGCCCTCTGGTAAGGTTGGGAA
GCCCTGCAAAGTAAACTGGATGGCTTTCTTGCCGCCAAGGATCTGATGGCGCAGGGGATCAA
GATCTGATCAAGAGACAGGATGAGGATCGTTTCGCATGATTGAACAAGATGGATTGCACGCA
GGTTCTCCGGCCGCTTGGGTGGAGAGGCTATTCGGCTATGACTGGGCACAACAGACAATCGG
CTGCTCTGATGCCGCCGTGTTCCGGCTGTCAGCGCAGGGGCGCCCGGTTCTTTTTGTCAAGA
CCGACCTGTCCGGTGCCCTGAATGAACTCCAAGACGAGGCAGCGCGGCTATCGTGGCTGGCC
ACGACGGGCGTTCCTTGCGCAGCTGTGCTCGACGTTGTCACTGAAGCGGGAAGGGACTGGCT
GCTATTGGGCGAAGTGCCGGGGCAGGATCTCCTGTCATCTCACCTTGCTCCTGCCGAGAAAG
TATCCATCATGGCTGATGCAATGCGGCGGCTGCATACGCTTGATCCGGCTACCTGCCCATTC
GACCACCAAGCGAAACATCGCATCGAGCGAGCACGTACTCGGATGGAAGCCGGTCTTGTCGA
TCAGGATGATCTGGACGAAGAGCATCAGGGGCTCGCGCCAGCCGAACTGTTCGCCAGGCTCA
AGGCGCGGATGCCCGACGGCGAGGATCTCGTCGTGACCCATGGCGATGCCTGCTTGCCGAAT
ATCATGGTGGAAAATGGCCGCTTTTCTGGATTCATCGACTGTGGCCGGCTGGGTGTGGCGGA
CCGCTATCAGGACATAGCGTTGGCTACCCGTGATATTGCTGAAGAGCTTGGCGGCGAATGGG
CTGACCGCTTCCTCGTGCTTTACGGTATCGCCGCTCCCGATTCGCAGCGCATCGCCTTCTAT
CGCCTTCTTGACGAGTTCTTCTGAGCGGGACTCTGGGGTTCGCTAGAGGATCGATCCTTTTT
AACCCATCACATATACCTGCCGTTCACTATTATTTAGTGAAATGAGATATTATGATATTTTC
TGAATTGTGATTAAAAAGGCAACTTTATGCCCATGCAACAGAAACTATAAAAAATACAGAGA
ATGAAAAGAAACAGATAGATTTTTTAGTTCTTTAGGCCCGTAGTCTGCAAATCCTTTTATGA
TTTTCTATCAAACAAAAGAGGAAAATAGACCAGTTGCAATCCAAACGAGAGTCTAATAGAAT
GAGGTCGAAAAGTAAATCGCGCGGGTTTGTTACTGATAAAGCAGGCAAGACCTAAAATGTGT
AAAGGGCAAAGTGTATACTTTGGCGTCACCCCTTACATATTTTAGGTCTTTTTTTATTGTGC
GTAACTAACTTGCCATCTTCAAACAGGAGGGCTGGAAGAAGCAGACCGCTAACACAGTACAT
AAAAAAGGAGACATGAACGATGAACATCAAAAAGTTTGCAAAACAAGCAACAGTATTAACCT
TTACTACCGCACTGCTGGCAGGAGGCGCAACTCAAGCGTTTGCGAAAGAAACGAACCAAAAG
CCATATAAGGAAACATACGGCATTTCCCATATTACACGCCATGATATGCTGCAAATCCCTGA
ACAGCAAAAAAATGAAAAATATCAAGTTTCTGAATTTGATTCGTCCACAATTAAAAATATCT

CTTCTGCAAAAGGCCTGGACGTTTGGGACAGCTGGCCATTACAAAACGCTGACGGCACTGTC
GCAAACTATCACGGCTACCACATCGTCTTTGCATTAGCCGGAGATCCTAAAAATGCGGATGA
CACATCGATTTACATGTTCTATCAAAAAGTCGGCGAAACTTCTATTGACAGCTGGAAAAACG
CTGGCCGCGTCTTTAAAGACAGCGACAAATTCGATGCAAATGATTCTATCCTAAAAGACCAA
ACACAAGAATGGTCAGGTTCAGCCACATTTACATCTGACGGAAAAATCCGTTTATTCTACAC
TGATTTCTCCGGTAAACATTACGGCAAACAAACACTGACAACTGCACAAGTTAACGTATCAG
CATCAGACAGCTCTTTGAACATCAACGGTGTAGAGGATTATAAATCAATCTTTGACGGTGAC
GGAAAAACGTATCAAAATGTACAGCAGTTCATCGATGAAGGCAACTACAGCTCAGGCGACAA
CCATACGCTGAGAGATCCTCACTACGTAGAAGATAAAGGCCACAAATACTTAGTATTTGAAG
CAAACACTGGAACTGAAGATGGCTACCAAGGCGAAGAATCTTTATTTAACAAAGCATACTAT
GGCAAAAGCACATCATTCTTCCGTCAAGAAAGTCAAAAACTTCTGCAAAGCGATAAAAAACG
CACGGCTGAGTTAGCAAACGGCGCTCTCGGTATGATTGAGCTAAACGATGATTACACACTGA
AAAAAGTGATGAAACCGCTGATTGCATCTAACACAGTAACAGATGAAATTGAACGCGCGAAC
GTCTTTAAAATGAACGGCAAATGGTACCTGTTCACTGACTCCCGCGGATCAAAAATGACGAT
TGACGGCATTACGTCTAACGATATTTACATGCTTGGTTATGTTTCTAATTCTTTAACTGGCC
CATACAAGCCGCTGAACAAAACTGGCCTTGTGTTAAAAATGGATCTTGATCCTAACGATGTA
ACCTTTACTTACTCACACTTCGCTGTACCTCAAGCGAAAGGAAACAATGTCGTGATTACAAG
CTATATGACAAACAGAGGATTCTACGCAGACAAACAATCAACGTTTGCGCCGAGCTTCCTGC
TGAACATCAAAGGCAAGAAAACATCTGTTGTCAAAGACAGCATCCTTGAACAAGGACAATTA
ACAGTTAACAAATAAAAACGCAAAAGAAAATGCCGATGGGTACCGAGCGAAATGACCGACCA
AGCGACGCCCAACCTGCCATCACGAGATTTCGATTCCACCGCCGCCTTCTATGAAAGGTTGG
GCTTCGGAATCGTTTTCCGGGACGCCCTCGCGGACGTGCTCATAGTCCACGACGCCCGTGAT
TTTGTAGCCCTGGCCGACGGCCAGCAGGTAGGCCGACAGGCTCATGCCGGCCGCCGCCGCCT
TTTCCTCAATCGCTCTTCGTTCGTCTGGAAGGCAGTACACCTTGATAGGTGGGCTGCCCTTC
CTGGTTGGCTTGGTTTCATCAGCCATCCGCTTGCCCTCATCTGTTACGCCGGCGGTAGCCGG
CCAGCCTCGCAGAGCAGGATTCCCGTTGAGCACCGCCAGGTGCGAATAAGGGACAGTGAAGA
AGGAACACCCGCTCGCGGGTGGGCCTACTTCACCTATCCTGCCCGGCTGACGCCGTTGGATA
CACCAAGGAAAGTCTACACGAACCCTTTGGCAAAATCCTGTATATCGTGCGAAAAAGGATGG
ATATACCGAAAAAATCGCTATAATGACCCCGAAGCAGGGTTATGCAGCGGAAAAGCGCTGCT
TCCCTGCTGTTTTGTGGAATATCTACCGACTGGAAACAGGCAAATGCAGGAAATTACTGAAC
TGAGGGGACAGGCGAGAGACGATGCCAAAGAGCTCCTGAAAATCTCGATAACTCAAAAAATA
CGCCCGGTAGTGATCTTATTTCATTATGGTGAAAGTTGGAACCTCTTACGTGCCGATCAACG
TCTCATTTTCGCCAAAAGTTGGCCCAGGGCTTCCCGGTATCAACAGGGACACCAGGATTTAT
TTATTCTGCGAAGTGATCTTCCGTCACAGGTATTTATTCGGCGCAAAGTGCGTCGGGTGATG

CTGCCAACTTACTGATTTAGTGTATGATGGTGTTTTTGAGGTGCTCCAGTGGCTTCTGTTTC
TATCAGCTCCTGAAAATCTCGATAACTCAAAAAATACGCCCGGTAGTGATCTTATTTCATTA
TGGTGAAAGTTGGAACCTCTTACGTGCCGATCAACGTCTCATTTTCGCCAAAAGTTGGCCCA
GGGCTTCCCGGTATCAACAGGGACACCAGGATTTATTTATTCTGCGAAGTGATCTTCCGTCA
CAGGTATTTATTCGGCGCAAAGTGCGTCGGGTGATGCTGCCAACTTACTGATTTAGTGTATG
ATGGTGTTTTTGAGGTGCTCCAGTGGCTTCTGTTTCTATCAGGGCTGGATGATCCTCCAGCG
CGGGGATCTCATGCTGGAGTTCTTCGCCCACCCCAAAAGGATCTAGGTGAAGATCCTTTTTG
ATAATCTCATGACCAAAATCCCTTAACGTGAGTTTTCGTTCCACTGAGCGTCAGACCCCGTA
GAAAAGATCAAACCATGTCAGCCGTTAAGTGTTCCTGTGTCACTCAAAATTGCTTTGAGAGG
CTCTAAGGGCTTCTCAGTGCGTTACATCCCTGGCTTGTTGTCCACAACCGTTAAACCTTAAA
AGCTTTAAAAGCCTTATATATTCTTTTTTTTCTTATAAAACTTAAAACCTTAGAGGCTATTT
AAGTTGCTGATTTATATTAATTTTATTGTTCAAACATGAGAGCTTAGTACGTGAAACATGAG
AGCTTAGTACGTTAGCCATGAGAGCTTAGTACGTTAGCCATGAGGGTTTAGTTCGTTAAACA
TGAGAGCTTAGTACGTTAAACATGAGAGCTTAGTACGTGAAACATGAGAGCTTAGTACGTAC
TATCAACAGGTTGAACTGCTGATCTTCAGATCGCCTTTTGCTCACATGTTCTTTCCTGCGTT
ATCCCCTGATTCTGTGGATAACCGTATTACCGCCTTTGAGTGAGCTGATACCGCTCGCCGCA
GCCGAACGACCGAGCGCAGCGAGTCAGTGAGCGAGGAAGCGGAAGAGCGCCCAATACGCAAA
CCGCCTCTCCCCGCGCGTTGGCCGATTCATTAATGCAGCTGGCACGACAGGTTTCCCGACTG
GAAAGCGGGCAGTGAGCGCAACGCAATTAATGTGAGTTAGCTCACTCATTAGGCACCCCAGG
CTTTACACTTTATGCTTCCGGCTCGTATGTTGTGTGGAATTGTGAGCGGATAACAATTTCAC
ACAGGAAACAGCTATGACATGATTACGAATTCCATAGACCAGCGGTTCGCCGTCGCAGCGCA
GCACGACCTCGCGGGCGTGCGTGCGGCCGCGGCAGGCGGGCAGGAGGCTTCGTTCGTTCCGA
CGCAGGGGGCTGCTGCCCTGGCGGACCGGCTGCACGGCGTAGTGCTCGCACACGGCTTGCAG
CCGCGCGCTCAGGGACCCGGCCCCGGTCAGCCAGTGCCGCAGTCGGCACCAGGGCAGGGGAC
GGGCAAACCAGGATGGCATGGACGATGGGAACAGGATGGGGACCGTCAAAGGGGCGCCATCA
TAACGACGGGGTAGGGATCGGCGCGGCCGGCGAAGCTGGCTACACTGCCGCCACATGAAGCT
GCACAACTACTTCCGGTCTTCCGCTTCGTTCCGCGTGCGCATCGCGCTGGCGCTCAAGGGCC
TGGACTACGAGTACGTGCCCGTCCACCTGGTCAAGGGCGAGCAGCTGCAGGCGCCGTTTGCC
CAGCTCTCCCCGGAGCGGCTGGTGCCGGTGCTGCAGGACGGCGACCAGACGCTCTCGCAGTC
GCTGGCCATCATCGAGTACCTCGACGAAACCCACCCCGAGCCGCCGCTGCTGCCGGCCGACC
CGCCGGGCCGCGCGCGGGTGCGGGCGCTGGCGCTGGACATCGCCTGCGAGATCCACCCGCTC
AACAACCTGCGCGTGCTGCGCTACCTGGTGCGCCAGCTGGGCGTGAGCGACGAGGCCAAGAA
CGGCTGGTACCGGCACTGGGTGGAAACCGGCCTGGAGGCGGTGGAGCGCCAGCTGGCCGGGC
ACCCGGCCACCGGCCGCTACTGCCACGGCGACACCCCCACGCTCGCGGACTGCGTGCTGGTG

CCGCAGATCTTCAACGCGCAGCGCTTCGACTGCCGGCTGGACCACGTGCCGACCGTCATGAA
GGTGTTCGAGCACTGCATGCAGCACCCGGCCTTCATCGCCGCGCAGCCGTCGCGCTGCCCCG
ACGCCGAGGCCTGAGGCGGCGATGGGCGAGGTCGACGCCACGCGGCCGGACACGGCCTGGTT
GCGCCCCGAGTGGCCGGCGCCGCCGGGCGTGCGGGCGCTGATGAGCACCCGACAGGGGGGCG
TCAGCCGCCCGCCTTACGATGGGCTGAACCTGGGCGACCACGTCGGCGACGACGCCGAGGCG
GTGCGGCGCAACCGCGAGCGCTTCGTCGCGGCGCTGCAGGCGCAGCCGGTGTTCCTGCAGCA
GGTGCACGGCACCACCGTGGTGCGGCTGGGACCGGACGACCTGCGCCGTGCCCGGCCGCACG
AGGCCGATGCGGCGATCACGACCGAGCCGGGCATCGCCTGCACGGTCATGGTGGCCGACTGC
CTGCCGGTGCTGTTCGCCAGCGCCGACGGCCGCGCGGTGGGTGCCGCCCATGCAGGCTGGCG
CGGGCTGTGCGCCGGCGTGCTGGAGCGCACCGTGCGCGCGCTGTGCGAGGCAGCCGGGTGCG
AGCCGGCGCGGCTGCTCGCCTGGCTCGGACCTTGCATCGGAGCCGATCGGTTCGAGGTGGGC
GACGAGGTGCGCCAGGCGTTCGTCGCGGTGCAGGACCGCGCCGGCGCCCGCTTCCGGCCGGG
GGCGGTGGCGGGCAAGTGGTGGGCCGACCTGCCCGGGCTGGCGAGGGACCGCCTGGCGGCCG
CGGGCGTGACCGCGGTCAGCGGCGGGCACTGGTGCACGGTGTCCGACCGCTCAAGGTTCTTT
TCGTTCCGGCGCGACGGGGTCACGGGGCGCATGGCGGCTGCCGTCTGGCGGGTCGCCGGCGC
CGGGGACTAGCGGGGACGCGGCGGCTTGCGCCTGCTCGGCGCGACGCCGGGCCTTGCGGCGC
GCAGGTGTACCCAACAGATACAGCACCAGTGCGACGGGGCCGAGCCCGTACAGAATGAAGGT
GAAGATCGCGCCGAGCACGGTTCCCTGGCTGCTGAAGGCCTCGGCCACGGCCATCATCAGCG
CGACGTAGAGCCAGGTGATTGCGACGAGATAGAGCACGCGTGGTTCCAGCACCGGGCGGCAG
GCCCGATTGACAACGATCAGAGAGGACCGAGCATTATGGGAGTCAGATGCAGGGCCGCGGGC
AGCGCGGGCGCCAGAGGGAGGCAAACGCCCGGACACCCGGTTTCGGGCGGCCCGGCTCGCGT
TCACTGACACGAGGAGAGGATGTGAAGGCGCCTTCCGGCATGAACAGATACCCGAACCCTTC
CAAGGAGCTTGAACATGTCTGACATCGTCATCGTTTCCGCCGCGCGAACGGCGGTCGGCAAG
TTCGGCGGCACGCTGGCGAAGACGCCGGCTGCCGAGCTGGGGGCCACCGTGATCAAGGAGGT
GCTGCGCCGCGCCGGCCTTTCGGGCGAGCAGGTCAGCGAGGTGATCATGGGCCAGGTGCTGC
AGGCCGGCTGCGGGCAGAACCCGGCGCGACAGGCGGTCATCAAGGCCGGGTTGCCGGAAGGC
GTGCCGGCGATGACCATCAACAAGGTGTGCGGCTCGGGCCTGAAGGCCGTGATGCTGGCGGC
GCAGGCCATCCGCGACGGCGACGCCGACATCGTCGTGGCCGGCGGGCAGGAGAACATGAGCC
TGGCGCCCCACGTGCTGCTCGGCTCGCGCGAGGGCCAGCGCATGGGTGACTGGAAGATGGTC
GACTCGATGATCACCGACGGCCTGTGGGACGTCTACAACCAGTACCACATGGGCATCACCGC
CGAGAACGTCGCGAAGAAGTACGGCATCAGCCGCGAGGAGCAGGACGCGCTGGCGCTGGCCT
CGCAGCAGAAGGCCGCCGCCGCGCAGGACGCCGGGCGCTTCAAGGACGAGATCGTGCCGGTG
GTGATCCCCCAGAGGAAGGGCGACCCGGTGGTGTTCGACACCGACGAGTTCATCAACCGCAA
GACCAGCGCCGAGGCGCTGGCCGGGCTGCGCCCGGCCTTCGACAAGGCGGGCACGGTGACCG

CGGGCAATGCCTCGGGCATCAACGACGGCGCGGCCGCGGTGGTGGTGATGAGCGCGAAGCGC
GCCGAGCAGCTGGGCCTCAAGCCGCTGGCGCGCATCGCCTCCTATGCCAGCGCCGGCCTGGA
TCCGGCCTACATGGGCATGGGCCCGGTGCCGGCGGCGCGCAAGGCGCTGGACCGCGCCGGCT
GGAAGCCGGCCGACCTCGACCTGCTCGAGATCAACGAGGCCTTCGCGGCGCAGGCCTGCGCG
GTGCACAAGGAAATGGGCTGGGACACCAGCAAGGTCAACGTCAACGGCGGCGCGATCGCGAT
CGGGCACCCGATCGGCGCGTCCGGCTGCCGCATCCTGGTCACGCTGCTGCACGAGATGCAGC
GGCGCGACGCCCGCAAGGGCATCGCCTCGCTGTGCATCGGCGGCGGCATGGGCGTGGCACTG
ACCGTCGAGCGCTGAACGTTGCGGCAATGGCCCGTCCGGTGACGGGGGAGGGGGAACCTGGG
CTTGACCCGTGGGGCCTTTGCGGCAACCCTCGTCACCGGCACAGACGAAACAGATACATCAG
GAGCAGAACATGGCACAGAAAGTTGCGTACGTCACCGGCGGCATGGGCGGTATCGGCACCGC
GATCTGCCAGCGCCTGGCACGCGATGGGTTCAAGGTCATCGCCGGCTGCGGGCCGAACCGCG
ACTACCAGAAGTGGCTCGACCAGCAGAAGGAGCTGGGCTACACCTTCTACGCCTCGGTGGGC
AACGTGGCCGACTGGGATTCGACGGTGGCCGCCTTCGCCAAGGCCAAGGCCGAGCACGGGCC
GATCGACGTGCTGGTCAACAACGCCGGCATCACCCGCGACCGCATGTTCCTGAAGATGACGC
CGGAGGACTGGCACGCGGTGATCAACACCAACCTCAACAGCATGTTCAACGTCACCAAGCAG
GTGGTGCCGGACATGGTGGAGCGGGGCTGGGGCCGCATCATCCAGATCTCCTCGGTCAACGG
CGAGAAGGGCCAGGCCGGGCAGACCAACTACTCGGCGGCCAAGGCCGGCATGCACGGCTTCA
CGATGGCGCTGGCGCAGGAGCTGGCCTCCAAGGGCGTGACGGTCAACACCGTGAGCCCCGGC
TACATCGGCACCGACATGGTCCGCGCGATCAAGCCCGAGATCCTGGAGAAGATCATCGCCAC
GATCCCGGTGCGGCGCCTGGGCACGCCGGAGGAAATCGCCTCCATCGTGTCCTGGGTGGCGT
CGGAGGAATCGGGCTTCGCGACCGGTGCCGATTTCTCGATCAACGGCGGCCTGCACATGGGC
TGAAGCTCGCCAGCCCTGCCGATGCGAAGAAGTCTAGAGTCGACCTGCAGGCATGCAAGCTT
GGCACTGGCCGTCGTTTTACAACGTCGTGACTGGGAAAACCCTGGCGTTACCCAACTTAATC
GCCTTGCAGCACATCCCCCTTTCGCCAGCTGGCGTAATAGCGAAGAGGCCCGCACCGATCGC
CCTTCCCAACAGTTGCGCAGCCTGAATGGCGAATGGCGATAAGCTAGCTTCACGC

**Plasmid pSNW2_HA2** - plasmid for deletion of *phaC* gene via homologous recombination and GFP-mediated selection (8,418 bp).

GCTTTCTCTTTGCGCTTGCGTTTTCCCTTGTCCAGATAGCCCAGTAGCTGACATTCATCCGG
GGTCAGCACCGTTTCTGCGGACTGGCTTTCTACGTGTTCCGCTTCCTTTAGCAGCCCTTGCG
CCCTGAGTGCTTGCGGCAGCGTGAAGCTAATTCCCATGTCAGCCGTTAAGTGTTCCTGTGTC
ACTCAAAATTGCTTTGAGAGGCTCTAAGGGCTTCTCAGTGCGTTACATCCCTGGCTTGTTGT
CCACAACCGTTAAACCTTAAAAGCTTTAAAAGCCTTATATATTCTTTTTTTTCTTATAAAAC

TTAAAACCTTAGAGGCTATTTAAGTTGCTGATTTATATTAATTTTATTGTTCAAACATGAGA
GCTTAGTACGTGAAACATGAGAGCTTAGTACGTTAGCCATGAGAGCTTAGTACGTTAGCCAT
GAGGGTTTAGTTCGTTAAACATGAGAGCTTAGTACGTTAAACATGAGAGCTTAGTACGTGAA
ACATGAGAGCTTAGTACGTACTATCAACAGGTTGAACTGCTGATCTTCAGATCCTCTACGCC
GGACGCATCGTGGCCGTTTTCCGCTGCATAACCCTGCTTCGGGGTCATTATAGCGATTTTTT
CGGTATATCCATCCTTTTTCGCACGATATACAGGATTTTGCCAAAGGGTTCGTGTAGACTTT
CCTTGGTGTATCCAACGGCGTCAGCCGGGCAGGATAGGTGAAGTAGGCCCACCCGCGAGCGG
GTGTTCCTTCTTCACTGTCCCTTATTCGCACCTGGCGGTGCTCAACGGGAATCCTGCTCTGC
GAGGCTGGCCGGCTACCGCCGGCGTAACAGATGAGGGCAAGCGGATGGCTGATGAAACCAAG
CCAACCAGGAAGGGCAGCCCACCTATCAAGGTGTACTGCCTTCCAGACGAACGAAGAGCGAT
TGAGGAAAAGGCGGCGGCGGCCGGCATGAGCCTGTCGGCCTACCTGCTGGCCGTCGGCCAGG
GCTACAAAATCACGGGCGTCGTGGACTATGAGCACGTCCGCGAGCTGGCCCGCATCAATGGC
GACCTGGGCCGCCTGGGCGGCCTGCTGAAACTCTGGCTCACCGACGACCCGCGCACGGCGCG
GTTCGGTGATGCCACGATCCTCGCCCTGCTGGCGAAGATCGAAGAGAAGCAGGACGAGCTTG
GCAAGGTCATGATGGGCGTGGTCCGCCCGAGGGCAGAGCCATGACTTTTTTAGCCGCTAAAA
CGGCCGGGGGGTGCGCGTGATTGCCAAGCACGTCCCCATGCGCTCCATCAAGAAGAGCGACT
TCGCGGAGCTGGTGAAGTACATCACCGACGAGCAAGGCAAGACCGACCAAAGCGGCCATCGT
GCCTCCCCACTCCTGCAGTTCGGGGGCATGGATGCGCGGATAGCCGCTGCTGGTTTCCTGGA
TGCCGACGGATTTGCACTGCCGGTAGAACTCCGCGAGGTCGTCCAGCCTCAGGCAGCAGCTG
AACCAACTCGCGAGGGGATCGAGCCCCATTCGCCATTCAGGCTGCGCAACTGTTGGGAAGGG
CGATCGGTGCGGGCCTCTTCGCTATTACGCCAGCTGGCGAAAGGGGGATGTGCTGCAAGGCG
ATTAAGTTGGGTAACGCCAGGGTTTTCCCAGTCACGACGTTGTAAAACGACGGCCAGTATAG
GGATAACAGGGTAATCTGAATTCCATAGACCAGCGGTTCGCCGTCGCAGCGCAGCACGACCT
CGCGGGCGTGCGTGCGGCCGCGGCAGGCGGGCAGGAGGCTTCGTTCGTTCCGACGCAGGGGG
CTGCTGCCCTGGCGGACCGGCTGCACGGCGTAGTGCTCGCACACGGCTTGCAGCCGCGCGCT
CAGGGACCCGGCCCCGGTCAGCCAGTGCCGCAGTCGGCACCAGGGCAGGGGACGGGCAAACC
AGGATGGCATGGACGATGGGAACAGGATGGGGACCGTCAAAGGGGCGCCATCATAACGACGG
GGTAGGGATCGGCGCGGCCGGCGAAGCTGGCTACACTGCCGCCACATGAAGCTGCACAACTA
CTTCCGGTCTTCCGCTTCGTTCCGCGTGCGCATCGCGCTGGCGCTCAAGGGCCTGGACTACG
AGTACGTGCCCGTCCACCTGGTCAAGGGCGAGCAGCTGCAGGCGCCGTTTGCCCAGCTCTCC
CCGGAGCGGCTGGTGCCGGTGCTGCAGGACGGCGACCAGACGCTCTCGCAGTCGCTGGCCAT
CATCGAGTACCTCGACGAAACCCACCCCGAGCCGCCGCTGCTGCCGGCCGACCCGCCGGGCC
GCGCGCGGGTGCGGGCGCTGGCGCTGGACATCGCCTGCGAGATCCACCCGCTCAACAACCTG
CGCGTGCTGCGCTACCTGGTGCGCCAGCTGGGCGTGAGCGACGAGGCCAAGAACGGCTGGTA

CCGGCACTGGGTGGAAACCGGCCTGGAGGCGGTGGAGCGCCAGCTGGCCGGGCACCCGGCCA
CCGGCCGCTACTGCCACGGCGACACCCCCACGCTCGCGGACTGCGTGCTGGTGCCGCAGATC
TTCAACGCGCAGCGCTTCGACTGCCGGCTGGACCACGTGCCGACCGTCATGAAGGTGTTCGA
GCACTGCATGCAGCACCCGGCCTTCATCGCCGCGCAGCCGTCGCGCTGCCCCGACGCCGAGG
CCTGAGGCGGCGATGGGCGAGGTCGACGCCACGCGGCCGGACACGGCCTGGTTGCGCCCCGA
GTGGCCGGCGCCGCCGGGCGTGCGGGCGCTGATGAGCACCCGACAGGGGGGCGTCAGCCGCC
CGCCTTACGATGGGCTGAACCTGGGCGACCACGTCGGCGACGACGCCGAGGCGGTGCGGCGC
AACCGCGAGCGCTTCGTCGCGGCGCTGCAGGCGCAGCCGGTGTTCCTGCAGCAGGTGCACGG
CACCACCGTGGTGCGGCTGGGACCGGACGACCTGCGCCGTGCCCGGCCGCACGAGGCCGATG
CGGCGATCACGACCGAGCCGGGCATCGCCTGCACGGTCATGGTGGCCGACTGCCTGCCGGTG
CTGTTCGCCAGCGCCGACGGCCGCGCGGTGGGTGCCGCCCATGCAGGCTGGCGCGGGCTGTG
CGCCGGCGTGCTGGAGCGCACCGTGCGCGCGCTGTGCGAGGCAGCCGGGTGCGAGCCGGCGC
GGCTGCTCGCCTGGCTCGGACCTTGCATCGGAGCCGATCGGTTCGAGGTGGGCGACGAGGTG
CGCCAGGCGTTCGTCGCGGTGCAGGACCGCGCCGGCGCCCGCTTCCGGCCGGGGGCGGTGGC
GGGCAAGTGGTGGGCCGACCTGCCCGGGCTGGCGAGGGACCGCCTGGCGGCCGCGGGCGTGA
CCGCGGTCAGCGGCGGGCACTGGTGCACGGTGTCCGACCGCTCAAGGTTCTTTTCGTTCCGG
CGCGACGGGGTCACGGGGCGCATGGCGGCTGCCGTCTGGCGGGTCGCCGGCGCCGGGGACTA
GCGGGGACGCGGCGGCTTGCGCCTGCTCGGCGCGACGCCGGGCCTTGCGGCGCGCAGGTGTA
CCCAACAGATACAGCACCAGTGCGACGGGGCCGAGCCCGTACAGAATGAAGGTGAAGATCGC
GCCGAGCACGGTTCCCTGGCTGCTGAAGGCCTCGGCCACGGCCATCATCAGCGCGACGTAGA
GCCAGGTGATTGCGACGAGATAGAGCACGCGTGGTTCCAGCACCGGGCGGCAGGCCCGATTG
ACAACGATCAGAGAGGACCGAGCATTATGGGAGTCAGATGCAGGGCCGCGGGCAGCGCGGGC
GCCAGAGGGAGGCAAACGCCCGGACACCCGGTTTCGGGCGGCCCGGCTCGCGTTCACTGACA
CGAGGAGAGGATGTGAAGGCGCCTTCCGGCATGAACAGATACCCGAACCCTTCCAAGGAGCT
TGAACATGTCTGACATCGTCATCGTTTCCGCCGCGCGAACGGCGGTCGGCAAGTTCGGCGGC
ACGCTGGCGAAGACGCCGGCTGCCGAGCTGGGGGCCACCGTGATCAAGGAGGTGCTGCGCCG
CGCCGGCCTTTCGGGCGAGCAGGTCAGCGAGGTGATCATGGGCCAGGTGCTGCAGGCCGGCT
GCGGGCAGAACCCGGCGCGACAGGCGGTCATCAAGGCCGGGTTGCCGGAAGGCGTGCCGGCG
ATGACCATCAACAAGGTGTGCGGCTCGGGCCTGAAGGCCGTGATGCTGGCGGCGCAGGCCAT
CCGCGACGGCGACGCCGACATCGTCGTGGCCGGCGGGCAGGAGAACATGAGCCTGGCGCCCC
ACGTGCTGCTCGGCTCGCGCGAGGGCCAGCGCATGGGTGACTGGAAGATGGTCGACTCGATG
ATCACCGACGGCCTGTGGGACGTCTACAACCAGTACCACATGGGCATCACCGCCGAGAACGT
CGCGAAGAAGTACGGCATCAGCCGCGAGGAGCAGGACGCGCTGGCGCTGGCCTCGCAGCAGA
AGGCCGCCGCCGCGCAGGACGCCGGGCGCTTCAAGGACGAGATCGTGCCGGTGGTGATCCCC

CAGAGGAAGGGCGACCCGGTGGTGTTCGACACCGACGAGTTCATCAACCGCAAGACCAGCGC
CGAGGCGCTGGCCGGGCTGCGCCCGGCCTTCGACAAGGCGGGCACGGTGACCGCGGGCAATG
CCTCGGGCATCAACGACGGCGCGGCCGCGGTGGTGGTGATGAGCGCGAAGCGCGCCGAGCAG
CTGGGCCTCAAGCCGCTGGCGCGCATCGCCTCCTATGCCAGCGCCGGCCTGGATCCGGCCTA
CATGGGCATGGGCCCGGTGCCGGCGGCGCGCAAGGCGCTGGACCGCGCCGGCTGGAAGCCGG
CCGACCTCGACCTGCTCGAGATCAACGAGGCCTTCGCGGCGCAGGCCTGCGCGGTGCACAAG
GAAATGGGCTGGGACACCAGCAAGGTCAACGTCAACGGCGGCGCGATCGCGATCGGGCACCC
GATCGGCGCGTCCGGCTGCCGCATCCTGGTCACGCTGCTGCACGAGATGCAGCGGCGCGACG
CCCGCAAGGGCATCGCCTCGCTGTGCATCGGCGGCGGCATGGGCGTGGCACTGACCGTCGAG
CGCTGAACGTTGCGGCAATGGCCCGTCCGGTGACGGGGGAGGGGGAACCTGGGCTTGACCCG
TGGGGCCTTTGCGGCAACCCTCGTCACCGGCACAGACGAAACAGATACATCAGGAGCAGAAC
ATGGCACAGAAAGTTGCGTACGTCACCGGCGGCATGGGCGGTATCGGCACCGCGATCTGCCA
GCGCCTGGCACGCGATGGGTTCAAGGTCATCGCCGGCTGCGGGCCGAACCGCGACTACCAGA
AGTGGCTCGACCAGCAGAAGGAGCTGGGCTACACCTTCTACGCCTCGGTGGGCAACGTGGCC
GACTGGGATTCGACGGTGGCCGCCTTCGCCAAGGCCAAGGCCGAGCACGGGCCGATCGACGT
GCTGGTCAACAACGCCGGCATCACCCGCGACCGCATGTTCCTGAAGATGACGCCGGAGGACT
GGCACGCGGTGATCAACACCAACCTCAACAGCATGTTCAACGTCACCAAGCAGGTGGTGCCG
GACATGGTGGAGCGGGGCTGGGGCCGCATCATCCAGATCTCCTCGGTCAACGGCGAGAAGGG
CCAGGCCGGGCAGACCAACTACTCGGCGGCCAAGGCCGGCATGCACGGCTTCACGATGGCGC
TGGCGCAGGAGCTGGCCTCCAAGGGCGTGACGGTCAACACCGTGAGCCCCGGCTACATCGGC
ACCGACATGGTCCGCGCGATCAAGCCCGAGATCCTGGAGAAGATCATCGCCACGATCCCGGT
GCGGCGCCTGGGCACGCCGGAGGAAATCGCCTCCATCGTGTCCTGGGTGGCGTCGGAGGAAT
CGGGCTTCGCGACCGGTGCCGATTTCTCGATCAACGGCGGCCTGCACATGGGCTGAAGCTCG
CCAGCCCTGCCGATGCGAAGAAGTCTAGAGTCGACCTGCAGGCATGCAAGCTTCTAGGGATA
ACAGGGTAATCCGGCGTAATCATGGTCATAGCTGTTTCCTGTGTGAAATTGTTATCCGCTCA
CAATTCCACACAACATACGAGCCGGAAGCATAAAGTGTAAAGCCTGGGGTGCCTAATGAGTG
AGCTAACTCACATTAATTGCGTTGCGCTCACTGCCCGCTTTCCAGTCGGGAAACCTGTCGTG
CCAGCTGCATTAATGAATCGGCCAACGCGCGGGGAGAGGCGGTTTGCGTATTGGGGGGTGGG
CGAAGAACTCCAGCATGAGATCCCCGCGCTGGAGGATCATCCAGCCGGCGTCCCGGAAAACG
ATTCCGAAGCCCAACCTTTCATAGAAGGCGGCGGTGGAATCGAAATCTCGTGATGGCAGGTT
GGGCGTCGCTTGGTCGGTCATTTCGAACCCCAGAGTCCCGCTCAGAAGAACTCGTCAAGAAG
GCGATAGAAGGCGATGCGCTGCGAATCGGGAGCGGCGATACCGTAAAGCACGAGGAAGCGGT
CAGCCCATTCGCCGCCAAGCTCTTCAGCAATATCACGGGTAGCCAACGCTATGTCCTGATAG
CGGTCCGCCACACCCAGCCGGCCACAGTCGATGAATCCAGAAAAGCGGCCATTTTCCACCAT

GATATTCGGCAAGCAGGCATCGCCATGGGTCACGACGAGATCCTCGCCGTCGGGCATGCGCG

CCTTGAGCCTGGCGAACAGTTCGGCTGGCGCGAGCCCCTGATGCTCTTCGTCCAGATCATCC

TGATCGACAAGACCGGCTTCCATCCGAGTACGTGCTCGCTCGATGCGATGTTTCGCTTGGTG

GTCGAATGGGCAGGTAGCCGGATCAAGCGTATGCAGCCGCCGCATTGCATCAGCCATGATGG

ATACTTTCTCGGCAGGAGCAAGGTGAGATGACAGGAGATCCTGCCCCGGCACTTCGCCCAAT

AGCAGCCAGTCCCTTCCCGCTTCAGTGACAACGTCGAGCACAGCTGCGCAAGGAACGCCCGT

CGTGGCCAGCCACGATAGCCGCGCTGCCTCGTCCTGCAGTTCATTCAGGGCACCGGACAGGT

CGGTCTTGACAAAAAGAACCGGGCGCCCCTGCGCTGACAGCCGGAACACGGCGGCATCAGAG

CAGCCGATTGTCTGTTGTGCCCAGTCATAGCCGAATAGCCTCTCCACCCAAGCGGCCGGAGA

ACCTGCGTGCAATCCATCTTGTTCAATCATGCGAAACGATCCTCATCCTGTCTCTTGATCAG

ATCTTGATCCCCTGCGCCATCAGATCCTTGGCGGCAAGAAAGCCATCCAGTTTACTTTGCAG

GGCTTCCCAACCTTACCAGAGGGCGCCCCAGCTGGCAATTCCGGTTCGCTTGCTGTCCATAA

AACCGCCCAGTCTAGCTATCGCCATGCCCATTGACAAGGCTCTCGCGGCCAGGTATAATTGC

ACGAGGGCCCAAGTTCACTTAAAAAGGAGATCAACAATGAAAGCAATTTTCGTACTGAAACA

TCTTAATCATGCTAAGGAGGTTTTCTAATGCGTAAAGGTGAAGAACTGTTCACCGGTGTTGT

TCCGATCCTGGTTGAACTGGATGGTGATGTTAACGGCCACAAATTCTCTGTTCGTGGTGAAG

GTGAAGGTGATGCAACCAACGGTAAACTGACCCTGAAATTCATCTGCACTACCGGTAAACTG

CCGGTTCCATGGCCGACTCTGGTGACTACCCTGACCTATGGTGTTCAGTGTTTTTCTCGTTA

CCCGGATCACATGAAGCAGCATGATTTCTTCAAATCTGCAATGCCGGAAGGTTATGTACAGG

AGCGCACCATTTCTTTCAAAGACGATGGCACCTACAAAACCCGTGCAGAGGTTAAATTTGAA

GGTGATACTCTGGTGAACCGTATTGAACTGAAAGGCATTGATTTCAAAGAGGACGGCAACAT

CCTGGGCCACAAACTGGAATATAACTTCAACTCCCATAACGTTTACATCACCGCAGACAAAC

AGAAGAACGGTATCAAAGCTAACTTCAAAATTCGCCATAACGTTGAAGACGGTAGCGTACAG

CTGGCGGACCACTACCAGCAGAACACTCCGATCGGTGATGGTCCGGTTCTGCTGCCGGATAA

CCACTACCTGTCCACCCAGTCTAAACTGTCCAAAGACCCGAACGAAAAGCGCGACCACATGG

TGCTGCTGGAGTTCGTTACTGCCGCAGGTATCACGCACGGCATGGATGAACTCTACAAATAA

ATGACTAGTCTTGGACTCCTGTTGATAGATCCAGTAATGACCTCAGAACTCCATCTGGATTT

GTTCAGAACGCTCGGTTGCCGCCGGGCGTTTTTTATTGGTGAGAATCCAGGGGTCCCCAATA

ATTACGATTTAAATTTGTGTCTCAAAATCTCTGATGTTACAGCTACCT

**Plasmid pSNW2_8585 -** plasmid for deletion of the gene encoding IS481_08585 restriction endonuclease (5,845 bp).

GCTTTCTCTTTGCGCTTGCGTTTTCCCTTGTCCAGATAGCCCAGTAGCTGACATTCATCCGG
GGTCAGCACCGTTTCTGCGGACTGGCTTTCTACGTGTTCCGCTTCCTTTAGCAGCCCTTGCG
CCCTGAGTGCTTGCGGCAGCGTGAAGCTAATTCCCATGTCAGCCGTTAAGTGTTCCTGTGTC
ACTCAAAATTGCTTTGAGAGGCTCTAAGGGCTTCTCAGTGCGTTACATCCCTGGCTTGTTGT
CCACAACCGTTAAACCTTAAAAGCTTTAAAAGCCTTATATATTCTTTTTTTTCTTATAAAAC
TTAAAACCTTAGAGGCTATTTAAGTTGCTGATTTATATTAATTTTATTGTTCAAACATGAGA
GCTTAGTACGTGAAACATGAGAGCTTAGTACGTTAGCCATGAGAGCTTAGTACGTTAGCCAT
GAGGGTTTAGTTCGTTAAACATGAGAGCTTAGTACGTTAAACATGAGAGCTTAGTACGTGAA
ACATGAGAGCTTAGTACGTACTATCAACAGGTTGAACTGCTGATCTTCAGATCCTCTACGCC
GGACGCATCGTGGCCGTTTTCCGCTGCATAACCCTGCTTCGGGGTCATTATAGCGATTTTTT
CGGTATATCCATCCTTTTTCGCACGATATACAGGATTTTGCCAAAGGGTTCGTGTAGACTTT
CCTTGGTGTATCCAACGGCGTCAGCCGGGCAGGATAGGTGAAGTAGGCCCACCCGCGAGCGG
GTGTTCCTTCTTCACTGTCCCTTATTCGCACCTGGCGGTGCTCAACGGGAATCCTGCTCTGC
GAGGCTGGCCGGCTACCGCCGGCGTAACAGATGAGGGCAAGCGGATGGCTGATGAAACCAAG
CCAACCAGGAAGGGCAGCCCACCTATCAAGGTGTACTGCCTTCCAGACGAACGAAGAGCGAT
TGAGGAAAAGGCGGCGGCGGCCGGCATGAGCCTGTCGGCCTACCTGCTGGCCGTCGGCCAGG
GCTACAAAATCACGGGCGTCGTGGACTATGAGCACGTCCGCGAGCTGGCCCGCATCAATGGC
GACCTGGGCCGCCTGGGCGGCCTGCTGAAACTCTGGCTCACCGACGACCCGCGCACGGCGCG
GTTCGGTGATGCCACGATCCTCGCCCTGCTGGCGAAGATCGAAGAGAAGCAGGACGAGCTTG
GCAAGGTCATGATGGGCGTGGTCCGCCCGAGGGCAGAGCCATGACTTTTTTAGCCGCTAAAA
CGGCCGGGGGGTGCGCGTGATTGCCAAGCACGTCCCCATGCGCTCCATCAAGAAGAGCGACT
TCGCGGAGCTGGTGAAGTACATCACCGACGAGCAAGGCAAGACCGACCAAAGCGGCCATCGT
GCCTCCCCACTCCTGCAGTTCGGGGGCATGGATGCGCGGATAGCCGCTGCTGGTTTCCTGGA
TGCCGACGGATTTGCACTGCCGGTAGAACTCCGCGAGGTCGTCCAGCCTCAGGCAGCAGCTG
AACCAACTCGCGAGGGGATCGAGCCCCATTCGCCATTCAGGCTGCGCAACTGTTGGGAAGGG
CGATCGGTGCGGGCCTCTTCGCTATTACGCCAGCTGGCGAAAGGGGGATGTGCTGCAAGGCG
ATTAAGTTGGGTAACGCCAGGGTTTTCCCAGTCACGACGTTGTAAAACGACGGCCAGTATAG
GGATAACAGGGTAATCTGAATTCGGCTTCGATGCCGTCTTCCTCCATGCCCCCGAAGGGAGG
GTTTGTCACGATCACGTCCACGCGATCCTTCGGCCCCCAGTCGCGCAGCGGACGGGCCAGGG
TGTTGTCGTGGCGGACATTGGAGGGAACGTCGATGCCGTGCAGGAGAAGGTTGGTCATGCAC

AGCACGTGCGGCAGGTGCTTTTTCTCGATGCCGGAGAAGCACTCCTGAATGGTGCGCTCGTC
GGCTTCAGTTCTTGCCTGCTTGCGCAGGTGTTCGATGGCGCACACCAGGAAGCCGCCGGTGC
CACAGGCGGGGTCGAGGATGGTCTCTCCGAGGCGGGGATTGACCTGCTCGACGATAAACTGG
GTGACGGCGCGCGGCGTGTAGAACTCGCCGGCATTGCCCGCGGACTGGAGATCCTTGAGCAG
CTTCTCGTAGATGTCGCCGAACATGTGGCGGTCATCGGAGGCATTGAAGTCGATGCCGTTGA
TCTTGTTGATCACCTGCCGCATCAATGTGCCGTTCTTCATGTAGTTGTTGGCGTCGGCGAAG
GTCGAGCGGATGAGTGCTGACATCGGGTCCGAACCGGGGAGCGATTCCAGCGCCGGAAGCAG
CTCATCATCGACAAAATTCAGAAGCGCATCTCCGGTGATACCTTCCTCGTCAGCCGCCCAGT
TGCGCCAGCGGAACTTCTCGGGGAGGGGCGATTTGTAGTCGTCCTGGAGAAGTTCGAGCTGG
GTCTCGCGGTCGTCGAAGATCTTGAGGAAGAACATCCATGCCATTTGCTCGAGCCGCTGGGC
GTCGCCGTAGGTGCCGGCGTCCTTGCGCATGATGTCCTGGATAGATTTGACGAGATTGGAGA
CGTTGGGCATGGGTCACATGCGGCAAAGATCTCTATTGTGAGGAGGTTGAAGGGCTGACCAT
GCAGTGCCACCATGCGATCTACGAGGTCGGCCCTACTCACGGGGCATGCCTGCGACGCCTGC
CCCCACCACCTCCCGGTTCAGAAGTGCGATCTGCTGATCCGCCGGGTTCCGGGTGGACGGGT
CAATGTCGAGCCCGAGACGCTTGAGCGTGTAGTAGAGCAGCGCCTTGCGGACGGTGACCCTG
GCCTTGCCGTTGCGCAGTCCGTAGTCGAGGGCGATGATCTTCTTCTGCGTGTCGGACAAACC
TGGGTGCGGGCCGATATCCAGCGTGATGTGCGTGTGCCAGTCGGCATCCTGGCTGTCGTCGA
TATCGCTGGGGCGGCTGTCCCGGGTGCCGAGGATGCGCGCCAACAGGAAGTCCTTGAAGCAG
CCGTCGGTGAAGCAGTATGCCCGGGCATGCCAGCGGAAGCCGTCGAAGGCGATGGCGTGCGG
TGCGATCCAGCGCCAGCGTGGGGCGGGGTTGGAAAGGGACCGGTACTCGACTTCAAGCGCTT
GCCTGGTACGAATCGCTTCGATGACGCGGCGCAACGTGTGGGCATCGATGCCGCGAACGGGG
CTGGGGACGATGTCGAAGGCAGGCATCTGGCCGATCCAGGTTTCCTCCTGCGTGGTGATGCC
GTCGGAGATCGACCGCAGCTGGGACAGGTAGCTGGCCGGGTCCGGTTTCAGAAACAGGGGCT
CGAAGCGGTCGCCCCGGACGTAGGTGCGGGCGCTCTTGTCATAGACCATGTTGCCCGGCGCC
ATGCCGAGGTAGCGGTTCAGGTCGGTGGATGCCTGGTTGATCGAGATCCCGAAGTCTAGAGT
CGACCTGCAGGCATGCAAGCTTCTAGGGATAACAGGGTAATCCGGCGTAATCATGGTCATAG
CTGTTTCCTGTGTGAAATTGTTATCCGCTCACAATTCCACACAACATACGAGCCGGAAGCAT
AAAGTGTAAAGCCTGGGGTGCCTAATGAGTGAGCTAACTCACATTAATTGCGTTGCGCTCAC
TGCCCGCTTTCCAGTCGGGAAACCTGTCGTGCCAGCTGCATTAATGAATCGGCCAACGCGCG
GGGAGAGGCGGTTTGCGTATTGGGGGGTGGGCGAAGAACTCCAGCATGAGATCCCCGCGCTG
GAGGATCATCCAGCCGGCGTCCCGGAAAACGATTCCGAAGCCCAACCTTTCATAGAAGGCGG
CGGTGGAATCGAAATCTCGTGATGGCAGGTTGGGCGTCGCTTGGTCGGTCATTTCGAACCCC
AGAGTCCCGCTCAGAAGAACTCGTCAAGAAGGCGATAGAAGGCGATGCGCTGCGAATCGGGA
GCGGCGATACCGTAAAGCACGAGGAAGCGGTCAGCCCATTCGCCGCCAAGCTCTTCAGCAAT

ATCACGGGTAGCCAACGCTATGTCCTGATAGCGGTCCGCCACACCCAGCCGGCCACAGTCGA
TGAATCCAGAAAAGCGGCCATTTTCCACCATGATATTCGGCAAGCAGGCATCGCCATGGGTC
ACGACGAGATCCTCGCCGTCGGGCATGCGCGCCTTGAGCCTGGCGAACAGTTCGGCTGGCGC
GAGCCCCTGATGCTCTTCGTCCAGATCATCCTGATCGACAAGACCGGCTTCCATCCGAGTAC
GTGCTCGCTCGATGCGATGTTTCGCTTGGTGGTCGAATGGGCAGGTAGCCGGATCAAGCGTA
TGCAGCCGCCGCATTGCATCAGCCATGATGGATACTTTCTCGGCAGGAGCAAGGTGAGATGA
CAGGAGATCCTGCCCCGGCACTTCGCCCAATAGCAGCCAGTCCCTTCCCGCTTCAGTGACAA
CGTCGAGCACAGCTGCGCAAGGAACGCCCGTCGTGGCCAGCCACGATAGCCGCGCTGCCTCG
TCCTGCAGTTCATTCAGGGCACCGGACAGGTCGGTCTTGACAAAAAGAACCGGGCGCCCCTG
CGCTGACAGCCGGAACACGGCGGCATCAGAGCAGCCGATTGTCTGTTGTGCCCAGTCATAGC
CGAATAGCCTCTCCACCCAAGCGGCCGGAGAACCTGCGTGCAATCCATCTTGTTCAATCATG
CGAAACGATCCTCATCCTGTCTCTTGATCAGATCTTGATCCCCTGCGCCATCAGATCCTTGG
CGGCAAGAAAGCCATCCAGTTTACTTTGCAGGGCTTCCCAACCTTACCAGAGGGCGCCCCAG
CTGGCAATTCCGGTTCGCTTGCTGTCCATAAAACCGCCCAGTCTAGCTATCGCCATGCCCAT
TGACAAGGCTCTCGCGGCCAGGTATAATTGCACGAGGGCCCAAGTTCACTTAAAAAGGAGAT
CAACAATGAAAGCAATTTTCGTACTGAAACATCTTAATCATGCTAAGGAGGTTTTCTAATGC
GTAAAGGTGAAGAACTGTTCACCGGTGTTGTTCCGATCCTGGTTGAACTGGATGGTGATGTT
AACGGCCACAAATTCTCTGTTCGTGGTGAAGGTGAAGGTGATGCAACCAACGGTAAACTGAC
CCTGAAATTCATCTGCACTACCGGTAAACTGCCGGTTCCATGGCCGACTCTGGTGACTACCC
TGACCTATGGTGTTCAGTGTTTTTCTCGTTACCCGGATCACATGAAGCAGCATGATTTCTTC
AAATCTGCAATGCCGGAAGGTTATGTACAGGAGCGCACCATTTCTTTCAAAGACGATGGCAC
CTACAAAACCCGTGCAGAGGTTAAATTTGAAGGTGATACTCTGGTGAACCGTATTGAACTGA
AAGGCATTGATTTCAAAGAGGACGGCAACATCCTGGGCCACAAACTGGAATATAACTTCAAC
TCCCATAACGTTTACATCACCGCAGACAAACAGAAGAACGGTATCAAAGCTAACTTCAAAAT
TCGCCATAACGTTGAAGACGGTAGCGTACAGCTGGCGGACCACTACCAGCAGAACACTCCGA
TCGGTGATGGTCCGGTTCTGCTGCCGGATAACCACTACCTGTCCACCCAGTCTAAACTGTCC
AAAGACCCGAACGAAAAGCGCGACCACATGGTGCTGCTGGAGTTCGTTACTGCCGCAGGTAT
CACGCACGGCATGGATGAACTCTACAAATAAATGACTAGTCTTGGACTCCTGTTGATAGATC
CAGTAATGACCTCAGAACTCCATCTGGATTTGTTCAGAACGCTCGGTTGCCGCCGGGCGTTT
TTTATTGGTGAGAATCCAGGGGTCCCCAATAATTACGATTTAAATTTGTGTCTCAAAATCTC
TGATGTTACAGCTACCT

**Plasmid pSNW2_14855 -** plasmid for deletion of the gene encoding IS481_14855 restriction endonuclease (6,148 bp).

GCTTTCTCTTTGCGCTTGCGTTTTCCCTTGTCCAGATAGCCCAGTAGCTGACATTCATCCGG
GGTCAGCACCGTTTCTGCGGACTGGCTTTCTACGTGTTCCGCTTCCTTTAGCAGCCCTTGCG
CCCTGAGTGCTTGCGGCAGCGTGAAGCTAATTCCCATGTCAGCCGTTAAGTGTTCCTGTGTC
ACTCAAAATTGCTTTGAGAGGCTCTAAGGGCTTCTCAGTGCGTTACATCCCTGGCTTGTTGT
CCACAACCGTTAAACCTTAAAAGCTTTAAAAGCCTTATATATTCTTTTTTTTCTTATAAAAC
TTAAAACCTTAGAGGCTATTTAAGTTGCTGATTTATATTAATTTTATTGTTCAAACATGAGA
GCTTAGTACGTGAAACATGAGAGCTTAGTACGTTAGCCATGAGAGCTTAGTACGTTAGCCAT
GAGGGTTTAGTTCGTTAAACATGAGAGCTTAGTACGTTAAACATGAGAGCTTAGTACGTGAA
ACATGAGAGCTTAGTACGTACTATCAACAGGTTGAACTGCTGATCTTCAGATCCTCTACGCC
GGACGCATCGTGGCCGTTTTCCGCTGCATAACCCTGCTTCGGGGTCATTATAGCGATTTTTT
CGGTATATCCATCCTTTTTCGCACGATATACAGGATTTTGCCAAAGGGTTCGTGTAGACTTT
CCTTGGTGTATCCAACGGCGTCAGCCGGGCAGGATAGGTGAAGTAGGCCCACCCGCGAGCGG
GTGTTCCTTCTTCACTGTCCCTTATTCGCACCTGGCGGTGCTCAACGGGAATCCTGCTCTGC
GAGGCTGGCCGGCTACCGCCGGCGTAACAGATGAGGGCAAGCGGATGGCTGATGAAACCAAG
CCAACCAGGAAGGGCAGCCCACCTATCAAGGTGTACTGCCTTCCAGACGAACGAAGAGCGAT
TGAGGAAAAGGCGGCGGCGGCCGGCATGAGCCTGTCGGCCTACCTGCTGGCCGTCGGCCAGG
GCTACAAAATCACGGGCGTCGTGGACTATGAGCACGTCCGCGAGCTGGCCCGCATCAATGGC
GACCTGGGCCGCCTGGGCGGCCTGCTGAAACTCTGGCTCACCGACGACCCGCGCACGGCGCG
GTTCGGTGATGCCACGATCCTCGCCCTGCTGGCGAAGATCGAAGAGAAGCAGGACGAGCTTG
GCAAGGTCATGATGGGCGTGGTCCGCCCGAGGGCAGAGCCATGACTTTTTTAGCCGCTAAAA
CGGCCGGGGGGTGCGCGTGATTGCCAAGCACGTCCCCATGCGCTCCATCAAGAAGAGCGACT
TCGCGGAGCTGGTGAAGTACATCACCGACGAGCAAGGCAAGACCGACCAAAGCGGCCATCGT
GCCTCCCCACTCCTGCAGTTCGGGGGCATGGATGCGCGGATAGCCGCTGCTGGTTTCCTGGA
TGCCGACGGATTTGCACTGCCGGTAGAACTCCGCGAGGTCGTCCAGCCTCAGGCAGCAGCTG
AACCAACTCGCGAGGGGATCGAGCCCCATTCGCCATTCAGGCTGCGCAACTGTTGGGAAGGG
CGATCGGTGCGGGCCTCTTCGCTATTACGCCAGCTGGCGAAAGGGGGATGTGCTGCAAGGCG
ATTAAGTTGGGTAACGCCAGGGTTTTCCCAGTCACGACGTTGTAAAACGACGGCCAGTATAG
GGATAACAGGGTAATCTGAATTCGAGCTCGGTACCCATGTCTAGCTGCTCGCTGAAGACTTG
ACGATATGTTTTCATTGCTTCGACCGCTTCAATCTGCATCAGCGCTTGCGCTTCGAGCGTAG
TGGGCTCTGTTGACTCGTTGCAAAACGCGAGTTTGAACAATGTTCTCAGTATGTTTTTGCTG

TACTCATCGAAGTTGCTCATGTCAAGCAACTCCATGTACGCAGAGTGAGTCTCAGCGCAAGC

GCACATATACGAGGCGAAACGCTCGTCATCGAGTTCACGCTCAAGCGTTCTGCGCGGCAGAT

TCTCGAGCGCTTTTGCTCGCTGCTTGTAGACAAGCTGAAGCTGAAATCGCGCTTCGTCTTCA

TCGAGCAGCTTGCCGCCGTCGCGCAGCGCACGAGAACGCAAGTAGTCAATGTCGTCTTGCGA

GACTTCTCTAGTAGCAAGTTCAGCTGCTTCACGCGCCGCGCGTGCTTTGATGTAAGCAAGCG

CAGCTTCTCTGTCAAACTTGTGCGGGCGAATGCCCGCGATGTAGATGCGGCGAATTCGCTTT

GCAGCGTCAACGAGAAACTTCCGCGCGTCAGTCAGCGCCTGCGCCTTCGGCGCGTGGCGGTT

ACCTGCCAGGTGTTGAGGGGCAATGCGAGGAGCGCGAGATGTAGCGAGAGTGCTGTTCACTA

TTTGTCTCCAATGAGAGAGTTGTTCATGTCCTGTATAGATTTGAGTTTTATACGCTTTGGCG

CTTTTGTAGATTTATTTTGAATGTAGATATGAGTCTAAATAAGTGACGGTCATATGAGGGCT

GTTCTGAGACGGGACTTGTCGCATCACGGCTGCAGCAGTGGCCACGGCGCGGGCAGCGCGTG

CTGAAGCTAGGCAGCGGCGGGCTTCGCGTTGCGTACAAAAGAAAAAGCCCCGCGATGGCGGG

GCTTTTTGGGGACAGAGCGCGCTGCGTGCTAGCGTCAAGCTAGAGATGGGTTTAGCTGACCG

CAACAGACCGCTCGATTTCGACGTCGAAACCAGCTAGGCTACATGTCTAGCTGCTCGCTGAA

GACTTGACGATATGTTTTCATTGCTTCGACCGCTTCAATCTGCATCAGCGCTTGCGCTTCGA

GCGTAGTGGGCTCTGTTGACTCGTTGCAAAACGCGAGTTTGAACAATGTTCTCAGTATGTTT

TTGCTGTACTCATCGAAGTTGCTCATGTCAAGCAACTCCATGTACGCAGAGTGAGTCTCAGC

GCAAGCGCACATATACGAGGCGAAACGCTCGTCATCGAGTTCACGCTCAAGCGTTCTGCGCG

GCAGATTCTCGAGCGCTTTTGCTCGCTGCTTGTAGACAAGCTGAAGCTGAAATCGCGCTTCG

TCTTCATCGAGCAGCTTGCCGCCGTCGCGCAGCGCACGAGAACGCAAGTAGTCAATGTCGTC

TTGCGAGACTTCTCTAGTAGCAAGTTCAGCTGCTTCACGCGCCGCGCGTGCTTTGATGTAAG

CAAGCGCAGCTTCTCTGTCAAACTTGTGCGGGCGAATGCCCGCGATGTAGATGCGGCGAATT

CGCTTTGCAGCGTCAACGAGAAACTTCCGCGCGTCAGTCAGCGCCTGCGCCTTCGGCGCGTG

GCGGTTACCTGCCAGGTGTTGAGGGGCAATGCGAGGAGCGCGAGATGTAGCGAGAGTGCTGT

TCACTATTTGTCTCCAATGAGAGAGTTGTTCATGTCCTGTATAGATTTGAGTTTTATACGCT

TTGGCGCTTTTGTAGATTTATTTTGAATGTAGATATGAGTCTAAATAAGTGACGGTCATATG

AGGGCTGTTCTGAGACGGGACTTGTCGCATCACGGCTGCAGCAGTGGCCACGGCGCGGGCAG

CGCGTGCTGAAGCTAGGCAGCGGCGGGCTTCGCGTTGCGTACAAAAGAAAAAGCCCCGCGAT

GGCGGGGCTTTTTGGGGACAGAGCGCGCTGCGTGCTAGCGTCAAGCTAGAGATGGGTTTAGC

TGACCGCAACAGACCGCTCGATTTCGACGTCGAAACCAGCTAGGCTATCTAGAGTCGACCTG

CAGGCATGCAAGCTTCTAGGGATAACAGGGTAATCCGGCGTAATCATGGTCATAGCTGTTTC

CTGTGTGAAATTGTTATCCGCTCACAATTCCACACAACATACGAGCCGGAAGCATAAAGTGT

AAAGCCTGGGGTGCCTAATGAGTGAGCTAACTCACATTAATTGCGTTGCGCTCACTGCCCGC

TTTCCAGTCGGGAAACCTGTCGTGCCAGCTGCATTAATGAATCGGCCAACGCGCGGGGAGAG

GCGGTTTGCGTATTGGGGGGTGGGCGAAGAACTCCAGCATGAGATCCCCGCGCTGGAGGATC
ATCCAGCCGGCGTCCCGGAAAACGATTCCGAAGCCCAACCTTTCATAGAAGGCGGCGGTGGA
ATCGAAATCTCGTGATGGCAGGTTGGGCGTCGCTTGGTCGGTCATTTCGAACCCCAGAGTCC
CGCTCAGAAGAACTCGTCAAGAAGGCGATAGAAGGCGATGCGCTGCGAATCGGGAGCGGCGA
TACCGTAAAGCACGAGGAAGCGGTCAGCCCATTCGCCGCCAAGCTCTTCAGCAATATCACGG
GTAGCCAACGCTATGTCCTGATAGCGGTCCGCCACACCCAGCCGGCCACAGTCGATGAATCC
AGAAAAGCGGCCATTTTCCACCATGATATTCGGCAAGCAGGCATCGCCATGGGTCACGACGA
GATCCTCGCCGTCGGGCATGCGCGCCTTGAGCCTGGCGAACAGTTCGGCTGGCGCGAGCCCC
TGATGCTCTTCGTCCAGATCATCCTGATCGACAAGACCGGCTTCCATCCGAGTACGTGCTCG
CTCGATGCGATGTTTCGCTTGGTGGTCGAATGGGCAGGTAGCCGGATCAAGCGTATGCAGCC
GCCGCATTGCATCAGCCATGATGGATACTTTCTCGGCAGGAGCAAGGTGAGATGACAGGAGA
TCCTGCCCCGGCACTTCGCCCAATAGCAGCCAGTCCCTTCCCGCTTCAGTGACAACGTCGAG
CACAGCTGCGCAAGGAACGCCCGTCGTGGCCAGCCACGATAGCCGCGCTGCCTCGTCCTGCA
GTTCATTCAGGGCACCGGACAGGTCGGTCTTGACAAAAAGAACCGGGCGCCCCTGCGCTGAC
AGCCGGAACACGGCGGCATCAGAGCAGCCGATTGTCTGTTGTGCCCAGTCATAGCCGAATAG
CCTCTCCACCCAAGCGGCCGGAGAACCTGCGTGCAATCCATCTTGTTCAATCATGCGAAACG
ATCCTCATCCTGTCTCTTGATCAGATCTTGATCCCCTGCGCCATCAGATCCTTGGCGGCAAG
AAAGCCATCCAGTTTACTTTGCAGGGCTTCCCAACCTTACCAGAGGGCGCCCCAGCTGGCAA
TTCCGGTTCGCTTGCTGTCCATAAAACCGCCCAGTCTAGCTATCGCCATGCCCATTGACAAG
GCTCTCGCGGCCAGGTATAATTGCACGAGGGCCCAAGTTCACTTAAAAAGGAGATCAACAAT
GAAAGCAATTTTCGTACTGAAACATCTTAATCATGCTAAGGAGGTTTTCTAATGCGTAAAGG
TGAAGAACTGTTCACCGGTGTTGTTCCGATCCTGGTTGAACTGGATGGTGATGTTAACGGCC
ACAAATTCTCTGTTCGTGGTGAAGGTGAAGGTGATGCAACCAACGGTAAACTGACCCTGAAA
TTCATCTGCACTACCGGTAAACTGCCGGTTCCATGGCCGACTCTGGTGACTACCCTGACCTA
TGGTGTTCAGTGTTTTTCTCGTTACCCGGATCACATGAAGCAGCATGATTTCTTCAAATCTG
CAATGCCGGAAGGTTATGTACAGGAGCGCACCATTTCTTTCAAAGACGATGGCACCTACAAA
ACCCGTGCAGAGGTTAAATTTGAAGGTGATACTCTGGTGAACCGTATTGAACTGAAAGGCAT
TGATTTCAAAGAGGACGGCAACATCCTGGGCCACAAACTGGAATATAACTTCAACTCCCATA
ACGTTTACATCACCGCAGACAAACAGAAGAACGGTATCAAAGCTAACTTCAAAATTCGCCAT
AACGTTGAAGACGGTAGCGTACAGCTGGCGGACCACTACCAGCAGAACACTCCGATCGGTGA
TGGTCCGGTTCTGCTGCCGGATAACCACTACCTGTCCACCCAGTCTAAACTGTCCAAAGACC
CGAACGAAAAGCGCGACCACATGGTGCTGCTGGAGTTCGTTACTGCCGCAGGTATCACGCAC
GGCATGGATGAACTCTACAAATAAATGACTAGTCTTGGACTCCTGTTGATAGATCCAGTAAT
GACCTCAGAACTCCATCTGGATTTGTTCAGAACGCTCGGTTGCCGCCGGGCGTTTTTTATTG

GTGAGAATCCAGGGGTCCCCAATAATTACGATTTAAATTTGTGTCTCAAAATCTCTGATGTT
ACAGCTACCT

**Plasmid pSNW2_14025 -** plasmid for deletion of the gene encoding IS481_14025 restriction endonuclease (6,521 bp).

GCTTTCTCTTTGCGCTTGCGTTTTCCCTTGTCCAGATAGCCCAGTAGCTGACATTCATCCGG
GGTCAGCACCGTTTCTGCGGACTGGCTTTCTACGTGTTCCGCTTCCTTTAGCAGCCCTTGCG
CCCTGAGTGCTTGCGGCAGCGTGAAGCTAATTCCCATGTCAGCCGTTAAGTGTTCCTGTGTC
ACTCAAAATTGCTTTGAGAGGCTCTAAGGGCTTCTCAGTGCGTTACATCCCTGGCTTGTTGT
CCACAACCGTTAAACCTTAAAAGCTTTAAAAGCCTTATATATTCTTTTTTTTCTTATAAAAC
TTAAAACCTTAGAGGCTATTTAAGTTGCTGATTTATATTAATTTTATTGTTCAAACATGAGA
GCTTAGTACGTGAAACATGAGAGCTTAGTACGTTAGCCATGAGAGCTTAGTACGTTAGCCAT
GAGGGTTTAGTTCGTTAAACATGAGAGCTTAGTACGTTAAACATGAGAGCTTAGTACGTGAA
ACATGAGAGCTTAGTACGTACTATCAACAGGTTGAACTGCTGATCTTCAGATCCTCTACGCC
GGACGCATCGTGGCCGTTTTCCGCTGCATAACCCTGCTTCGGGGTCATTATAGCGATTTTTT
CGGTATATCCATCCTTTTTCGCACGATATACAGGATTTTGCCAAAGGGTTCGTGTAGACTTT
CCTTGGTGTATCCAACGGCGTCAGCCGGGCAGGATAGGTGAAGTAGGCCCACCCGCGAGCGG
GTGTTCCTTCTTCACTGTCCCTTATTCGCACCTGGCGGTGCTCAACGGGAATCCTGCTCTGC
GAGGCTGGCCGGCTACCGCCGGCGTAACAGATGAGGGCAAGCGGATGGCTGATGAAACCAAG
CCAACCAGGAAGGGCAGCCCACCTATCAAGGTGTACTGCCTTCCAGACGAACGAAGAGCGAT
TGAGGAAAAGGCGGCGGCGGCCGGCATGAGCCTGTCGGCCTACCTGCTGGCCGTCGGCCAGG
GCTACAAAATCACGGGCGTCGTGGACTATGAGCACGTCCGCGAGCTGGCCCGCATCAATGGC
GACCTGGGCCGCCTGGGCGGCCTGCTGAAACTCTGGCTCACCGACGACCCGCGCACGGCGCG
GTTCGGTGATGCCACGATCCTCGCCCTGCTGGCGAAGATCGAAGAGAAGCAGGACGAGCTTG
GCAAGGTCATGATGGGCGTGGTCCGCCCGAGGGCAGAGCCATGACTTTTTTAGCCGCTAAAA
CGGCCGGGGGGTGCGCGTGATTGCCAAGCACGTCCCCATGCGCTCCATCAAGAAGAGCGACT
TCGCGGAGCTGGTGAAGTACATCACCGACGAGCAAGGCAAGACCGACCAAAGCGGCCATCGT
GCCTCCCCACTCCTGCAGTTCGGGGGCATGGATGCGCGGATAGCCGCTGCTGGTTTCCTGGA
TGCCGACGGATTTGCACTGCCGGTAGAACTCCGCGAGGTCGTCCAGCCTCAGGCAGCAGCTG
AACCAACTCGCGAGGGGATCGAGCCCCATTCGCCATTCAGGCTGCGCAACTGTTGGGAAGGG
CGATCGGTGCGGGCCTCTTCGCTATTACGCCAGCTGGCGAAAGGGGGATGTGCTGCAAGGCG

ATTAAGTTGGGTAACGCCAGGGTTTTCCCAGTCACGACGTTGTAAAACGACGGCCAGTATAG
GGATAACAGGGTAATCTGAATTCGCAATCTGATCTTCGAGCACGTTGGAAGGTATCGACCCG
CCCGACTCAATCTTGTCAAACGACGAGTTGATGTTGTGACCGGAGGGTCGCACCGTTGTCTT
CACGCCCTTTTTCGCCAGCCGGAGCATGTCCGCGCTGTATGGGCGAAGTACGTTGCGGTTGT
TCGCCTTCGGGTGAGGTGTCTTGGACAGCCACCAGACGTGCTCAACTGAATCCTTGATCCGA
ACCCTGCGCACCGTCACCCACTCCGCGGGCACGGGCATCTTTGCCGGGTTGTACCAGTAGCA
TTCTTGGGCGAGGTGAAAGCCCAGCTCCTCTACAAGCGCGATCATCAGCTTGTAGTGGTACA
GCGACCGCGTGGGCGACCCCGGATTCCAGCTCCCCCCGATATTCAAGACGAAGCTGCCGTCA
TCGGTCAGCACGCGTTGAATCTCGCGTGCGAACGGCAGGAACCAGTCCACGTAGTCCGCCTT
GCTGACGTTTCCGTACTCCTTCTTGAAGTGGAGCGCATAGGGTGGCGACGTGAATGCCAGAT
TGACACAGCCGTCCGGCATCGCCCTCAAGGTATCAAGCGAATCGCCCAAATAGGCCGCCCCC
GCTGCGGTGCGGTAGAACGGCTTGAGCTTCCGAACCAGCTTGGCGGGGTGTTCCTGCGGCTT
GGTGAACAGCTCGGGTTGTGGGGTTTCAATCTTCATTCTTGTCTTGGGCACCGGGGCAACCG
TTGGTAACATGCATCTCTGTGAGGGTAAGTATAGCAAGAAATCCATTGACAAAGTCAACCAA
TAAATCCGTTTCATTCGACCCTTCGGACTGGATCACTCAGGCCGAAGCCGCACGCCTACGCC
GCGTCACGAGACAGGCTATCGCCCGGCTGGTCCAGAACGGTCGCCTGCGCACGCTGGAAATA
GGCGGGCGATCATTCGTCAATCGTTCGGATGTGCTGGCGTTTGAGCCCAGTCCCCCGGGAAG
GCCGAGGACCTCGAATCATGTGAAATAAAGAAGCCGCCCGAGGGCGGCTTTTTTCATCGTCT
CACAGGGCCTTACCCCATCGCATCCCATCACCTCGAGGAAGGCGGGTTTTTAGGCGGGTTTA
GCGCCAATGCCCTGCGGTAAAGCCAGTGAAACTGGCAAATATGGCGGAAAGGGAGGGATTCG
AACCCTCGTCACGGGAAAACCCGTGAACCGGATTTCGAGTCCGGCGCATTCGACCACTCTGC
CACCTTTCCTTGTGTTCTCGTGGTCGCTACAAGAGCGAAGCTCGCCATTGTAGCAGGTTATG
GGCGCAGTTCGGTAAGGCCGCCCATGTAGGGGCGCAGCACCTCCGGCACGGTGACGCTGCCG
TCGGCGTTCTGGTAGTTCTCCAGCACCGCCACCAGCGCACGGCCGACGGCCAGGCCCGAGCC
GTTGAGGGTGTGCACCAGCTCGGGCTTGCCCTTCTCGTTCTTGAAGCGGGCCTGCATGCGGC
GCGCCTGGAAGGCCTCGCAGTTGGAGCAGGAGCTGATCTCGCGGTAGGTACCCTGCGCCGGG
ATCCAGACCTCCAGGTCGTAGGTCTTGGCCGCGCCGAAGCCCATGTCGCCGGTGCACAGCGT
CACGACGCGGTACGGCAGGCCGAGTTTCTGCAGGATGGCCTCGGCGTGGCCGACCATCTCTT
CCAGCGCCTCGTAGCTCTTCTGCGGGTGCACGATCTGCACCATCTCGACCTTGTCGAACTGG
TGCTGGCGGATCATGCCGCGCGTGTCCTTGCCGTAGCTGCCGGCCTCGGAGCGGAAGCAGGG
GCTGTGCGCGGTCAGCTTGATCGGCAGCTGCTCGGCGGCGAGGATCTGCTCGCGCACCGAGT
TGGTCAGCGAGATCTCCGAGGTGGAGATCAGGTACTGCACCAACCCGCTGTCGTCGCCCCCG
CGCGTGACCCAGAACATGTCCTCCTTGAACTTGGGCAGCTGGCCGGTGCCGACCAGCACGTC
GGCGTTGACGATGTAGGGCGTGTAGCACTCGGTGTAGCCGTGCTCGCCGGTCTGCACGTCGA

GCATGAACTGCGCGAGCGCGCGGTGCAGGCGCGCCACCGGGCCGCGCAGGAAGGCGAAGCGC

GCGCCCGAGAGCCTGGCGCCGGTGTCGAAGTCCAGCCCCAGCGGCGCGCCCAGGTCGACGTG

GTCCTTGACCGGGAAGTCGAACTGGCGCGGCGTGCCCCAGCGGCGCACTTCCACGTTGCCGG

TCTCGTCGGCGCCGACCGGCACGCTCTCGTGCGGCAGGTTGGGCACCTGCAGCAGCATCTCG

TTCAGCTCGGACTGGATCGCCTCGAGCCGCTCGGCGGAGGCCTTCAGCTCGTCGCCGATGCC

GCCGACCTCGGCCATCACGGCGGAGGTGTCCTCGCCCTTGGCCTTGAGTCTAGAGTCGACCT

GCAGGCATGCAAGCTTCTAGGGATAACAGGGTAATCCGGCGTAATCATGGTCATAGCTGTTT

CCTGTGTGAAATTGTTATCCGCTCACAATTCCACACAACATACGAGCCGGAAGCATAAAGTG

TAAAGCCTGGGGTGCCTAATGAGTGAGCTAACTCACATTAATTGCGTTGCGCTCACTGCCCG

CTTTCCAGTCGGGAAACCTGTCGTGCCAGCTGCATTAATGAATCGGCCAACGCGCGGGGAGA

GGCGGTTTGCGTATTGGGGGGTGGGCGAAGAACTCCAGCATGAGATCCCCGCGCTGGAGGAT

CATCCAGCCGGCGTCCCGGAAAACGATTCCGAAGCCCAACCTTTCATAGAAGGCGGCGGTGG

AATCGAAATCTCGTGATGGCAGGTTGGGCGTCGCTTGGTCGGTCATTTCGAACCCCAGAGTC

CCGCTCAGAAGAACTCGTCAAGAAGGCGATAGAAGGCGATGCGCTGCGAATCGGGAGCGGCG

ATACCGTAAAGCACGAGGAAGCGGTCAGCCCATTCGCCGCCAAGCTCTTCAGCAATATCACG

GGTAGCCAACGCTATGTCCTGATAGCGGTCCGCCACACCCAGCCGGCCACAGTCGATGAATC

CAGAAAAGCGGCCATTTTCCACCATGATATTCGGCAAGCAGGCATCGCCATGGGTCACGACG

AGATCCTCGCCGTCGGGCATGCGCGCCTTGAGCCTGGCGAACAGTTCGGCTGGCGCGAGCCC

CTGATGCTCTTCGTCCAGATCATCCTGATCGACAAGACCGGCTTCCATCCGAGTACGTGCTC

GCTCGATGCGATGTTTCGCTTGGTGGTCGAATGGGCAGGTAGCCGGATCAAGCGTATGCAGC

CGCCGCATTGCATCAGCCATGATGGATACTTTCTCGGCAGGAGCAAGGTGAGATGACAGGAG

ATCCTGCCCCGGCACTTCGCCCAATAGCAGCCAGTCCCTTCCCGCTTCAGTGACAACGTCGA

GCACAGCTGCGCAAGGAACGCCCGTCGTGGCCAGCCACGATAGCCGCGCTGCCTCGTCCTGC

AGTTCATTCAGGGCACCGGACAGGTCGGTCTTGACAAAAAGAACCGGGCGCCCCTGCGCTGA

CAGCCGGAACACGGCGGCATCAGAGCAGCCGATTGTCTGTTGTGCCCAGTCATAGCCGAATA

GCCTCTCCACCCAAGCGGCCGGAGAACCTGCGTGCAATCCATCTTGTTCAATCATGCGAAAC

GATCCTCATCCTGTCTCTTGATCAGATCTTGATCCCCTGCGCCATCAGATCCTTGGCGGCAA

GAAAGCCATCCAGTTTACTTTGCAGGGCTTCCCAACCTTACCAGAGGGCGCCCCAGCTGGCA

ATTCCGGTTCGCTTGCTGTCCATAAAACCGCCCAGTCTAGCTATCGCCATGCCCATTGACAA

GGCTCTCGCGGCCAGGTATAATTGCACGAGGGCCCAAGTTCACTTAAAAAGGAGATCAACAA

TGAAAGCAATTTTCGTACTGAAACATCTTAATCATGCTAAGGAGGTTTTCTAATGCGTAAAG

GTGAAGAACTGTTCACCGGTGTTGTTCCGATCCTGGTTGAACTGGATGGTGATGTTAACGGC

CACAAATTCTCTGTTCGTGGTGAAGGTGAAGGTGATGCAACCAACGGTAAACTGACCCTGAA

ATTCATCTGCACTACCGGTAAACTGCCGGTTCCATGGCCGACTCTGGTGACTACCCTGACCT

ATGGTGTTCAGTGTTTTTCTCGTTACCCGGATCACATGAAGCAGCATGATTTCTTCAAATCT
GCAATGCCGGAAGGTTATGTACAGGAGCGCACCATTTCTTTCAAAGACGATGGCACCTACAA
AACCCGTGCAGAGGTTAAATTTGAAGGTGATACTCTGGTGAACCGTATTGAACTGAAAGGCA
TTGATTTCAAAGAGGACGGCAACATCCTGGGCCACAAACTGGAATATAACTTCAACTCCCAT
AACGTTTACATCACCGCAGACAAACAGAAGAACGGTATCAAAGCTAACTTCAAAATTCGCCA
TAACGTTGAAGACGGTAGCGTACAGCTGGCGGACCACTACCAGCAGAACACTCCGATCGGTG
ATGGTCCGGTTCTGCTGCCGGATAACCACTACCTGTCCACCCAGTCTAAACTGTCCAAAGAC
CCGAACGAAAAGCGCGACCACATGGTGCTGCTGGAGTTCGTTACTGCCGCAGGTATCACGCA
CGGCATGGATGAACTCTACAAATAAATGACTAGTCTTGGACTCCTGTTGATAGATCCAGTAA
TGACCTCAGAACTCCATCTGGATTTGTTCAGAACGCTCGGTTGCCGCCGGGCGTTTTTTATT
GGTGAGAATCCAGGGGTCCCCAATAATTACGATTTAAATTTGTGTCTCAAAATCTCTGATGT
TACAGCTACCT

**Plasmid pAI_ThSacB -** plasmid for deletion of *phaC* gene via homologous recombination and thermostable SacB-mediated sucrose counter-selection (9,086 bp).

GCTTTCTCTTTGCGCTTGCGTTTTCCCTTGTCCAGATAGCCCAGTAGCTGACATTCATCCGG
GGTCAGCACCGTTTCTGCGGACTGGCTTTCTACGTGTTCCGCTTCCTTTAGCAGCCCTTGCG
CCCTGAGTGCTTGCGGCAGCGTGAAGCTAATTCCCATGTCAGCCGTTAAGTGTTCCTGTGTC
ACTCAAAATTGCTTTGAGAGGCTCTAAGGGCTTCTCAGTGCGTTACATCCCTGGCTTGTTGT
CCACAACCGTTAAACCTTAAAAGCTTTAAAAGCCTTATATATTCTTTTTTTTCTTATAAAAC
TTAAAACCTTAGAGGCTATTTAAGTTGCTGATTTATATTAATTTTATTGTTCAAACATGAGA
GCTTAGTACGTGAAACATGAGAGCTTAGTACGTTAGCCATGAGAGCTTAGTACGTTAGCCAT
GAGGGTTTAGTTCGTTAAACATGAGAGCTTAGTACGTTAAACATGAGAGCTTAGTACGTGAA
ACATGAGAGCTTAGTACGTACTATCAACAGGTTGAACTGCTGATCTTCAGATCCTCTACGCC
GGACGCATCGTGGCCGTTTTCCGCTGCATAACCCTGCTTCGGGGTCATTATAGCGATTTTTT
CGGTATATCCATCCTTTTTCGCACGATATACAGGATTTTGCCAAAGGGTTCGTGTAGACTTT
CCTTGGTGTATCCAACGGCGTCAGCCGGGCAGGATAGGTGAAGTAGGCCCACCCGCGAGCGG
GTGTTCCTTCTTCACTGTCCCTTATTCGCACCTGGCGGTGCTCAACGGGAATCCTGCTCTGC
GAGGCTGGCCGGCTACCGCCGGCGTAACAGATGAGGGCAAGCGGATGGCTGATGAAACCAAG
CCAACCAGGAAGGGCAGCCCACCTATCAAGGTGTACTGCCTTCCAGACGAACGAAGAGCGAT
TGAGGAAAAGGCGGCGGCGGCCGGCATGAGCCTGTCGGCCTACCTGCTGGCCGTCGGCCAGG
GCTACAAAATCACGGGCGTCGTGGACTATGAGCACGTCCGCGAGCTGGCCCGCATCAATGGC
GACCTGGGCCGCCTGGGCGGCCTGCTGAAACTCTGGCTCACCGACGACCCGCGCACGGCGCG

GTTCGGTGATGCCACGATCCTCGCCCTGCTGGCGAAGATCGAAGAGAAGCAGGACGAGCTTG
GCAAGGTCATGATGGGCGTGGTCCGCCCGAGGGCAGAGCCATGACTTTTTTAGCCGCTAAAA
CGGCCGGGGGGTGCGCGTGATTGCCAAGCACGTCCCCATGCGCTCCATCAAGAAGAGCGACT
TCGCGGAGCTGGTGAAGTACATCACCGACGAGCAAGGCAAGACCGACCAAAGCGGCCATCGT
GCCTCCCCACTCCTGCAGTTCGGGGGCATGGATGCGCGGATAGCCGCTGCTGGTTTCCTGGA
TGCCGACGGATTTGCACTGCCGGTAGAACTCCGCGAGGTCGTCCAGCCTCAGGCAGCAGCTG
AACCAACTCGCGAGGGGATCGAGCCCCATTCGCCATTCAGGCTGCGCAACTGTTGGGAAGGG
CGATCGGTGCGGGCCTCTTCGCTATTACGCCAGCTGGCGAAAGGGGGATGTGCTGCAAGGCG
ATTAAGTTGGGTAACGCCAGGGTTTTCCCAGTCACGACGTTGTAAAACGACGGCCAGTATAG
GGATAACAGGGTAATCTGAATTCCATAGACCAGCGGTTCGCCGTCGCAGCGCAGCACGACCT
CGCGGGCGTGCGTGCGGCCGCGGCAGGCGGGCAGGAGGCTTCGTTCGTTCCGACGCAGGGGG
CTGCTGCCCTGGCGGACCGGCTGCACGGCGTAGTGCTCGCACACGGCTTGCAGCCGCGCGCT
CAGGGACCCGGCCCCGGTCAGCCAGTGCCGCAGTCGGCACCAGGGCAGGGGACGGGCAAACC
AGGATGGCATGGACGATGGGAACAGGATGGGGACCGTCAAAGGGGCGCCATCATAACGACGG
GGTAGGGATCGGCGCGGCCGGCGAAGCTGGCTACACTGCCGCCACATGAAGCTGCACAACTA
CTTCCGGTCTTCCGCTTCGTTCCGCGTGCGCATCGCGCTGGCGCTCAAGGGCCTGGACTACG
AGTACGTGCCCGTCCACCTGGTCAAGGGCGAGCAGCTGCAGGCGCCGTTTGCCCAGCTCTCC
CCGGAGCGGCTGGTGCCGGTGCTGCAGGACGGCGACCAGACGCTCTCGCAGTCGCTGGCCAT
CATCGAGTACCTCGACGAAACCCACCCCGAGCCGCCGCTGCTGCCGGCCGACCCGCCGGGCC
GCGCGCGGGTGCGGGCGCTGGCGCTGGACATCGCCTGCGAGATCCACCCGCTCAACAACCTG
CGCGTGCTGCGCTACCTGGTGCGCCAGCTGGGCGTGAGCGACGAGGCCAAGAACGGCTGGTA
CCGGCACTGGGTGGAAACCGGCCTGGAGGCGGTGGAGCGCCAGCTGGCCGGGCACCCGGCCA
CCGGCCGCTACTGCCACGGCGACACCCCCACGCTCGCGGACTGCGTGCTGGTGCCGCAGATC
TTCAACGCGCAGCGCTTCGACTGCCGGCTGGACCACGTGCCGACCGTCATGAAGGTGTTCGA
GCACTGCATGCAGCACCCGGCCTTCATCGCCGCGCAGCCGTCGCGCTGCCCCGACGCCGAGG
CCTGAGGCGGCGATGGGCGAGGTCGACGCCACGCGGCCGGACACGGCCTGGTTGCGCCCCGA
GTGGCCGGCGCCGCCGGGCGTGCGGGCGCTGATGAGCACCCGACAGGGGGGCGTCAGCCGCC
CGCCTTACGATGGGCTGAACCTGGGCGACCACGTCGGCGACGACGCCGAGGCGGTGCGGCGC
AACCGCGAGCGCTTCGTCGCGGCGCTGCAGGCGCAGCCGGTGTTCCTGCAGCAGGTGCACGG
CACCACCGTGGTGCGGCTGGGACCGGACGACCTGCGCCGTGCCCGGCCGCACGAGGCCGATG
CGGCGATCACGACCGAGCCGGGCATCGCCTGCACGGTCATGGTGGCCGACTGCCTGCCGGTG
CTGTTCGCCAGCGCCGACGGCCGCGCGGTGGGTGCCGCCCATGCAGGCTGGCGCGGGCTGTG
CGCCGGCGTGCTGGAGCGCACCGTGCGCGCGCTGTGCGAGGCAGCCGGGTGCGAGCCGGCGC
GGCTGCTCGCCTGGCTCGGACCTTGCATCGGAGCCGATCGGTTCGAGGTGGGCGACGAGGTG

CGCCAGGCGTTCGTCGCGGTGCAGGACCGCGCCGGCGCCCGCTTCCGGCCGGGGGCGGTGGC
GGGCAAGTGGTGGGCCGACCTGCCCGGGCTGGCGAGGGACCGCCTGGCGGCCGCGGGCGTGA
CCGCGGTCAGCGGCGGGCACTGGTGCACGGTGTCCGACCGCTCAAGGTTCTTTTCGTTCCGG
CGCGACGGGGTCACGGGGCGCATGGCGGCTGCCGTCTGGCGGGTCGCCGGCGCCGGGGACTA
GCGGGGACGCGGCGGCTTGCGCCTGCTCGGCGCGACGCCGGGCCTTGCGGCGCGCAGGTGTA
CCCAACAGATACAGCACCAGTGCGACGGGGCCGAGCCCGTACAGAATGAAGGTGAAGATCGC
GCCGAGCACGGTTCCCTGGCTGCTGAAGGCCTCGGCCACGGCCATCATCAGCGCGACGTAGA
GCCAGGTGATTGCGACGAGATAGAGCACGCGTGGTTCCAGCACCGGGCGGCAGGCCCGATTG
ACAACGATCAGAGAGGACCGAGCATTATGGGAGTCAGATGCAGGGCCGCGGGCAGCGCGGGC
GCCAGAGGGAGGCAAACGCCCGGACACCCGGTTTCGGGCGGCCCGGCTCGCGTTCACTGACA
CGAGGAGAGGATGTGAAGGCGCCTTCCGGCATGAACAGATACCCGAACCCTTCCAAGGAGCT
TGAACATGTCTGACATCGTCATCGTTTCCGCCGCGCGAACGGCGGTCGGCAAGTTCGGCGGC
ACGCTGGCGAAGACGCCGGCTGCCGAGCTGGGGGCCACCGTGATCAAGGAGGTGCTGCGCCG
CGCCGGCCTTTCGGGCGAGCAGGTCAGCGAGGTGATCATGGGCCAGGTGCTGCAGGCCGGCT
GCGGGCAGAACCCGGCGCGACAGGCGGTCATCAAGGCCGGGTTGCCGGAAGGCGTGCCGGCG
ATGACCATCAACAAGGTGTGCGGCTCGGGCCTGAAGGCCGTGATGCTGGCGGCGCAGGCCAT
CCGCGACGGCGACGCCGACATCGTCGTGGCCGGCGGGCAGGAGAACATGAGCCTGGCGCCCC
ACGTGCTGCTCGGCTCGCGCGAGGGCCAGCGCATGGGTGACTGGAAGATGGTCGACTCGATG
ATCACCGACGGCCTGTGGGACGTCTACAACCAGTACCACATGGGCATCACCGCCGAGAACGT
CGCGAAGAAGTACGGCATCAGCCGCGAGGAGCAGGACGCGCTGGCGCTGGCCTCGCAGCAGA
AGGCCGCCGCCGCGCAGGACGCCGGGCGCTTCAAGGACGAGATCGTGCCGGTGGTGATCCCC
CAGAGGAAGGGCGACCCGGTGGTGTTCGACACCGACGAGTTCATCAACCGCAAGACCAGCGC
CGAGGCGCTGGCCGGGCTGCGCCCGGCCTTCGACAAGGCGGGCACGGTGACCGCGGGCAATG
CCTCGGGCATCAACGACGGCGCGGCCGCGGTGGTGGTGATGAGCGCGAAGCGCGCCGAGCAG
CTGGGCCTCAAGCCGCTGGCGCGCATCGCCTCCTATGCCAGCGCCGGCCTGGATCCGGCCTA
CATGGGCATGGGCCCGGTGCCGGCGGCGCGCAAGGCGCTGGACCGCGCCGGCTGGAAGCCGG
CCGACCTCGACCTGCTCGAGATCAACGAGGCCTTCGCGGCGCAGGCCTGCGCGGTGCACAAG
GAAATGGGCTGGGACACCAGCAAGGTCAACGTCAACGGCGGCGCGATCGCGATCGGGCACCC
GATCGGCGCGTCCGGCTGCCGCATCCTGGTCACGCTGCTGCACGAGATGCAGCGGCGCGACG
CCCGCAAGGGCATCGCCTCGCTGTGCATCGGCGGCGGCATGGGCGTGGCACTGACCGTCGAG
CGCTGAACGTTGCGGCAATGGCCCGTCCGGTGACGGGGGAGGGGGAACCTGGGCTTGACCCG
TGGGGCCTTTGCGGCAACCCTCGTCACCGGCACAGACGAAACAGATACATCAGGAGCAGAAC
ATGGCACAGAAAGTTGCGTACGTCACCGGCGGCATGGGCGGTATCGGCACCGCGATCTGCCA
GCGCCTGGCACGCGATGGGTTCAAGGTCATCGCCGGCTGCGGGCCGAACCGCGACTACCAGA

AGTGGCTCGACCAGCAGAAGGAGCTGGGCTACACCTTCTACGCCTCGGTGGGCAACGTGGCC
GACTGGGATTCGACGGTGGCCGCCTTCGCCAAGGCCAAGGCCGAGCACGGGCCGATCGACGT
GCTGGTCAACAACGCCGGCATCACCCGCGACCGCATGTTCCTGAAGATGACGCCGGAGGACT
GGCACGCGGTGATCAACACCAACCTCAACAGCATGTTCAACGTCACCAAGCAGGTGGTGCCG
GACATGGTGGAGCGGGGCTGGGGCCGCATCATCCAGATCTCCTCGGTCAACGGCGAGAAGGG
CCAGGCCGGGCAGACCAACTACTCGGCGGCCAAGGCCGGCATGCACGGCTTCACGATGGCGC
TGGCGCAGGAGCTGGCCTCCAAGGGCGTGACGGTCAACACCGTGAGCCCCGGCTACATCGGC
ACCGACATGGTCCGCGCGATCAAGCCCGAGATCCTGGAGAAGATCATCGCCACGATCCCGGT
GCGGCGCCTGGGCACGCCGGAGGAAATCGCCTCCATCGTGTCCTGGGTGGCGTCGGAGGAAT
CGGGCTTCGCGACCGGTGCCGATTTCTCGATCAACGGCGGCCTGCACATGGGCTGAAGCTCG
CCAGCCCTGCCGATGCGAAGAAGTCTAGAGTCGACCTGCAGGCATGCAAGCTTCTAGGGATA
ACAGGGTAATCCGGCGTAATCATGGTCATAGCTGTTTCCTGTGTGAAATTGTTATCCGCTCA
CAATTCCACACAACATACGAGCCGGAAGCATAAAGTGTAAAGCCTGGGGTGCCTAATGAGTG
AGCTAACTCACATTAATTGCGTTGCGCTCACTGCCCGCTTTCCAGTCGGGAAACCTGTCGTG
CCAGCTGCATTAATGAATCGGCCAACGCGCGGGGAGAGGCGGTTTGCGTATTGGGGGGTGGG
CGAAGAACTCCAGCATGAGATCCCCGCGCTGGAGGATCATCCAGCCGGCGTCCCGGAAAACG
ATTCCGAAGCCCAACCTTTCATAGAAGGCGGCGGTGGAATCGAAATCTCGTGATGGCAGGTT
GGGCGTCGCTTGGTCGGTCATTTCGAACCCCAGAGTCCCGCTCAGAAGAACTCGTCAAGAAG
GCGATAGAAGGCGATGCGCTGCGAATCGGGAGCGGCGATACCGTAAAGCACGAGGAAGCGGT
CAGCCCATTCGCCGCCAAGCTCTTCAGCAATATCACGGGTAGCCAACGCTATGTCCTGATAG
CGGTCCGCCACACCCAGCCGGCCACAGTCGATGAATCCAGAAAAGCGGCCATTTTCCACCAT
GATATTCGGCAAGCAGGCATCGCCATGGGTCACGACGAGATCCTCGCCGTCGGGCATGCGCG
CCTTGAGCCTGGCGAACAGTTCGGCTGGCGCGAGCCCCTGATGCTCTTCGTCCAGATCATCC
TGATCGACAAGACCGGCTTCCATCCGAGTACGTGCTCGCTCGATGCGATGTTTCGCTTGGTG
GTCGAATGGGCAGGTAGCCGGATCAAGCGTATGCAGCCGCCGCATTGCATCAGCCATGATGG
ATACTTTCTCGGCAGGAGCAAGGTGAGATGACAGGAGATCCTGCCCCGGCACTTCGCCCAAT
AGCAGCCAGTCCCTTCCCGCTTCAGTGACAACGTCGAGCACAGCTGCGCAAGGAACGCCCGT
CGTGGCCAGCCACGATAGCCGCGCTGCCTCGTCCTGCAGTTCATTCAGGGCACCGGACAGGT
CGGTCTTGACAAAAAGAACCGGGCGCCCCTGCGCTGACAGCCGGAACACGGCGGCATCAGAG
CAGCCGATTGTCTGTTGTGCCCAGTCATAGCCGAATAGCCTCTCCACCCAAGCGGCCGGAGA
ACCTGCGTGCAATCCATCTTGTTCAATCATGCGAAACGATCCTCATCCTGTCTCTTGATCAG
ATCTTGATCCCCTGCGCCATCAGATCCTTGGCGGCAAGAAAGCCATCCAGTTTACTTTGCAG
GGCTTCCCAACCTTACCAGAGGGCGCCCCAGCTGGCAATTCCGGTTCGCTTGCTGTCCATAA
AACCGCCCAGTCTAGCTATCGCCATGCCCATTGACAAGGCTCTCGCGGCCAGGTATAATTGC

ACGAATTACATATTGAAAAAGGGAGGAAATATTGATGAACATCAAGAACATCGCCAAGAAGG
CCAGCGCCCTGACCGTGGCCGCCGCCCTGCTGGCCGGCGGCGCCCCGCAGACCTTCGCCAAG
GAGACCCAGGACTACAAGAAGAGCTACGGCTTCAGCCACATCACCCGCCACGACATGCTGAA
GATCCCGGAGCAGCAGAAGAGCGAGCAGTTCAAGGTGCCGCAGTTCGACCCGAAGACCATCA
AGAACATCCCGAGCGCCAAGGGCTACAACAAGAACGGCGAGCTGATCGACCTGGACGTGTGG
GACAGCTGGCCGCTGCAGAACGCCGACGGCACCGTGGCCACCTACCACGGCTACAACCTGGT
GTTCGCCCTGGCCGGCGACCCGAAGGACGTGGACGACACCAGCATCTACCTGTTCTACCAGA
AGAAGGGCGAGACCAGCATCGACAGCTGGAAGAACGCCGGCCGCGTGTTCAAGGACAGCGAC
AAGTTCGTGCCGGACGACCCGTACCTGAAGCACCAGACCCAGGAGTGGAGCGGCAGCGCCAC
CCTGACCAAGGACGGCAAGGTGCGCCTGTTCTACACCGCCTTCAGCGGCACCCAGTACGGCA
AGCAGACCCTGACCACCGCCCAGGTGAACTTCAGCCAGCCGGACAGCGACACCCTGAAGATC
GACGGCGTGGAGGACCACAAGAGCGTGTTCGACGGCGCCGACGGCACCGTGTACCAGAACGT
GCAGCAGTTCATCGACGAGGGCAACTACAGCAGCGGCGACAACCACACCATGCGCGACCCGC
ACTACGTGGAGGACCGCGGCCACAAGTACCTGGTGTTCGAGGACAACACCGGCACCAAGACC
GGCTACCAGGGCGAGGACAGCCTGTTCAACCGCGCCTACTACGGCGGCAGCAAGAAGTTCTT
CAAGGAGGAGAGCAGCAAGCTGCTGCAGGGCGCCAACAAGAAGAACGCCAGCCTGGCCAACG
GCGCCCTGGGCATCATCGAGCTGAACAACGACTACACCCTGAAGAAGGTGATGAAGCCGCTG
ATCGCCAGCAACACCGTGACCGACGAGATCGAGCGCGCCAACCTGTTCAAGATGAACGGCAA
GTGGTACCTGTTCACCGACAGCCGCGGCAGCAAGATGACCATCGACGGCATCGGCAGCAAGG
ACATCTACATGCTGGGCTACGTGAGCGGCAGCCTGACCGGCCCGTTCAAGCCGCTGAACAAG
AGCGGCCTGGTGCTGCACATGGACCAGGACTACAACGACATCACCTTCACCTACAGCCACTT
CGCCGTGCCGCAGAAGAAGGGCGACGAGGTGGTGATCACCAGCTACATCACCAACCGCGGCA
TCAGCAACGAGCACCACGCCACCTTCGCCCCGAGCTTCCTGCTGAAGATCAAGGGCAGCAAG
ACCAGCGTGGTGAAGAACAGCATCCTGGAGCAGGGCCAGCTGACCGTGAACAAGTGACTTGG
ACTCCTGTTGATAGATCCAGTAATGACCTCAGAACTCCATCTGGATTTGTTCAGAACGCTCG
GTTGCCGCCGGGCGTTTTTTATTGGTGAGAATCCAGGGGTCCCCAATAATTACGATTTAAAT
TTGTGTCTCAAAATCTCTGATGTTACAGCTACCT
